# Phosphorylation regulates tau’s phase separation behavior and interactions with chromatin

**DOI:** 10.1101/2023.12.21.572911

**Authors:** Lannah S. Abasi, Nesreen Elathram, Manasi Movva, Amar Deep, Kevin D. Corbett, Galia T. Debelouchina

## Abstract

Tau is a microtubule-associated protein often found in neurofibrillary tangles (NFTs) in the brains of patients with Alzheimer’s disease (AD). Beyond this context, mounting evidence suggests that tau localizes into the nucleus, where it may play a role in DNA protection and heterochromatin regulation. Models of tau depletion or pathology show loss of genetically silent heterochromatin, aberrant expression of heterochromatic genes, and transposable element activation. The molecular mechanisms behind these observations are currently unclear. Using *in vitro* biophysical experiments, here we demonstrate that tau can undergo liquid-liquid phase separation (LLPS) with DNA, mononucleosomes, and reconstituted nucleosome arrays under low salt conditions. Low concentrations of tau promote chromatin compaction and protect DNA from digestion. While the material state of samples at physiological salt is dominated by chromatin oligomerization, tau can still associate strongly and reversibly with nucleosome arrays. These properties are driven by tau’s strong interactions with linker and nucleosomal DNA, while magic angle spinning (MAS) solid-state NMR experiments show that tau binding does not drastically alter nucleosome structure and dynamics. In addition, tau co-localizes into droplets formed by nucleosome arrays and phosphorylated HP1α, a key heterochromatin constituent thought to function through an LLPS mechanism. Importantly, LLPS and chromatin interactions are disrupted by aberrant tau hyperphosphorylation. These biophysical properties suggest that tau may directly impact DNA and chromatin accessibility and that loss of these interactions could contribute to the aberrant nuclear effects seen in tau pathology.

## Introduction

Tau is a neuron-specific microtubule-associated protein, well known for forming abnormally hyperphosphorylated neurofibrillary tangles (NFTs) within the brains of patients with Alzheimer’s disease (AD) and other neurodegenerative diseases called tauopathies.^1^ While it is known that the presence of NFTs correlates with cognitive decline,^2^ the mechanistic link between tau dysfunction and pathology remains unclear. From a biophysical perspective, tau’s ability to form insoluble β-sheet rich fibrils within NFTs has been perplexing since tau is highly charged and ordinarily exists as a soluble intrinsically disordered protein. Tau consists of an amino-terminal projection domain, a proline-rich sequence (PRD), a microtubule binding domain (MTBD), and a C-terminal domain **(Fig. S1a).** In neurons, tau can be expressed as one of six alternatively spliced isoforms (0N3R, 0N4R, 1N3R, 1N4R, 2N3R, or 2N4R), which are characterized according to the number of N-terminal inserts in the projection domain (0N, 1N, 2N) and the number of pseudorepeats in the MTBD (3R or 4R). Tau’s sequence is enriched in polar and charged residues that are distributed in a dipolar pattern: the N-terminus is negatively charged while positive charge is concentrated in the middle and C-terminal regions. This results in an overall net positive charge of ∼ 6 at pH 7.2. Tau is extensively regulated by phosphorylation, with 85 known phosphorylatable sites in its largest isoform (2N4R).^3^

Tau’s lack of a defined fold and its charge patterning give it a capacity to form weak multivalent interactions and predispose it to liquid-liquid phase separation (LLPS). Tau’s phase separation has been widely reported in the presence of RNA, polyanions, and crowding agents *in vitro* at physiological protein concentrations.^1,4–11^ It has been established that this process is electrostatically-driven,^4,5^ and that it can be enhanced by phosphorylation and the tauopathy-associated P301L mutant.^6,11^ The biological role of tau LLPS is currently being explored and may be connected to tau’s canonical microtubule polymerization role. In LLPS promoting conditions, tau droplets have the ability to assemble tubulin into stable microtubule bundles *in vitro,*^7^ a process that is hindered by tau phosphorylation.^12^ In addition, there are numerous reports that liquid droplets can progress into gel-like states and aggregates, which has led to speculation that LLPS could promote pathological fibrillization.^6,11,13^ While these observations are intriguing, the cellular functions of tau phase separation have not been established and remain unclear.

While most tau studies so far have been focused on its microtubule polymerization function and aberrant aggregation, a smaller pool of tau (∼16% of total tau)^14^ is nuclear localized, accumulating primarily in the nucleolus and in chromatin.^14–21^ Despite lacking a nuclear localization sequence, tau has been found to be associated with nuclear proteins,^16,22^ has strong interactions with nucleic acids,^5,23–27^ and has been suggested to play a role in DNA and RNA protection,^19,20,28–30^ as well as heterochromatin regulation.^20,31^ Stress conditions such as heat shock and oxidative stress induce an increased translocation of tau into the nucleus, which is associated with a DNA-protective effect, while tau depletion is associated with genomic instability.^29,30,32^ In addition, a heterochromatin regulatory role has been implied by loss of function studies. Tau has been found to associate with DNA within pericentromeric and perinucleolar heterochromatin in mouse neurons,^19,31^ rDNA nucleolar heterochromatin^20^ in human neuroblastoma cells, and heterochromatin in human neurons.^17^ Intriguingly, depletion of tau severely disrupts the distribution of epigenetic marks and proteins associated with transcriptional repression, including DNA methylation, histone 3 (H3) K9 methylation, and heterochromatin protein 1α (HP1α), and correlates with aberrant expression of heterochromatically silenced genes.^20,31^ This can be reversed with overexpression of nuclear-targeted tau.^31^ Heterochromatin relaxation has similarly been observed in Drosophila and mouse tauopathy models and in AD patient brains.^33,34^ In other studies, loss of heterochromatin and transposable element activation have been observed in AD patient brain tissue and are correlated with NFT burden, further implicating pathological tau with aberrant gene expression.^35,36^ These processes have been linked to the interactions of cytoplasmic tau with the actin cytoskeleton, causing nuclear envelope invaginations and destabilization of the lamin nucleoskeleton.^34,37–40^ While it is unknown if nuclear tau influences heterochromatin stability through its localization within heterochromatin^19^ and through its DNA and RNA binding properties, ^5,23–27^ loss of nuclear tau and changes in its phosphorylation state have been observed with AD progression.^17,41,42^ These observations suggest that the loss of function of nuclear tau may be a pathologically relevant event.

While the mechanism of tau’s nuclear function remains enigmatic, tau’s ability to interact with nucleic acids has been well-established both *in vitro* and in cells. Tau can preferentially interact with tRNA in cells,^5^ and tau aggregates can co-localize with small nuclear RNAs (snRNAs), small nucleolar RNAs (snoRNAs), mRNA, and tRNA.^43^ In addition, tau can phase separate with multiple types of RNA.^5^ It has also been demonstrated that tau can bind DNA both *in vitro*^23–27^ and in cells,^19,20,44^ through interactions with the DNA minor groove.^27^ Studies have implicated tau’s PRD and MTBD in both DNA^25^ and RNA^45^ binding, and this binding is thought to be sequence non-specific.^25,46^

Tau’s interactions with nucleic acids in the heterochromatin context are especially intriguing, since this is an environment thought to form and function through phase separation.^47,48^ In addition, tau co-localizes with heterochromatin protein HP1α, which promotes gene repression using a mechanism which is hypothesized to involve LLPS.^47,49–53^ Inspired by these observations, here we sought to explore the question of whether tau plays a role in heterochromatin organization through a phase separation mechanism. Using biophysical assays, we characterize the interactions of tau with DNA, mononucleosomes, and nucleosome arrays *in vitro* and assess its ability to induce LLPS under different ionic conditions and in the presence and absence of phosphorylated HP1α (pHP1α). We also characterize tau’s interactions with nucleosomes using magic angle spinning (MAS) NMR spectroscopy, determine the contributions of the different domains of tau, and explore the effects of AD-relevant tau phosphorylation on these interactions. Our data provide new insights into the enigmatic function of tau in the nucleus and the potential impact of hyperphosphorylation in disrupting tau’s nuclear interaction network.

## Results

### Tau undergoes LLPS with chromatin and DNA at low salt

Tau is found in chromatin environments and is known to interact with both DNA and RNA. However, it is unknown whether tau modulates DNA protection and heterochromatin regulation in a manner that involves direct interactions with chromatin. To explore this idea, we characterized tau’s interactions with both DNA and chromatin starting with low salt conditions where chromatin is less likely to undergo self-association. We expressed and purified 1N4R tau (**Fig. S1b-d**), the isoform thought to be enriched in the nucleus,^54^ and combined it with DNA, mononucleosomes, or reconstituted chromatin polymers containing 12 nucleosomes (12mer arrays)^55^ (schem atic of constructs in **Fig. S2a** and **Fig. 1a**). The 12mer arrays were prepared using recombinant histone octamers and a 2.1 kbp DNA template containing 12 tandem repeats of a strong nucleosome positioning sequence (“Widom 601”) with 30 bp of linker DNA (177 bp per tandem repeat).^55^ Upon combination of tau with DNA, mononucleosomes, or 12mer arrays in a buffer containing 25 mM NaCl, tau spontaneously formed spherical liquid droplets (**Fig. 1a**), which could fuse and wet the glass surface. The localization of tau was visualized by adding 5% Cy5-labeled tau (**Fig. 1a)**, while the DNA-intercalating fluorophore YOYO-1 was used to report on the localization of 12mer arrays, revealing that they are concentrated within the droplets (**Fig. S3a**).

**Figure 1.**
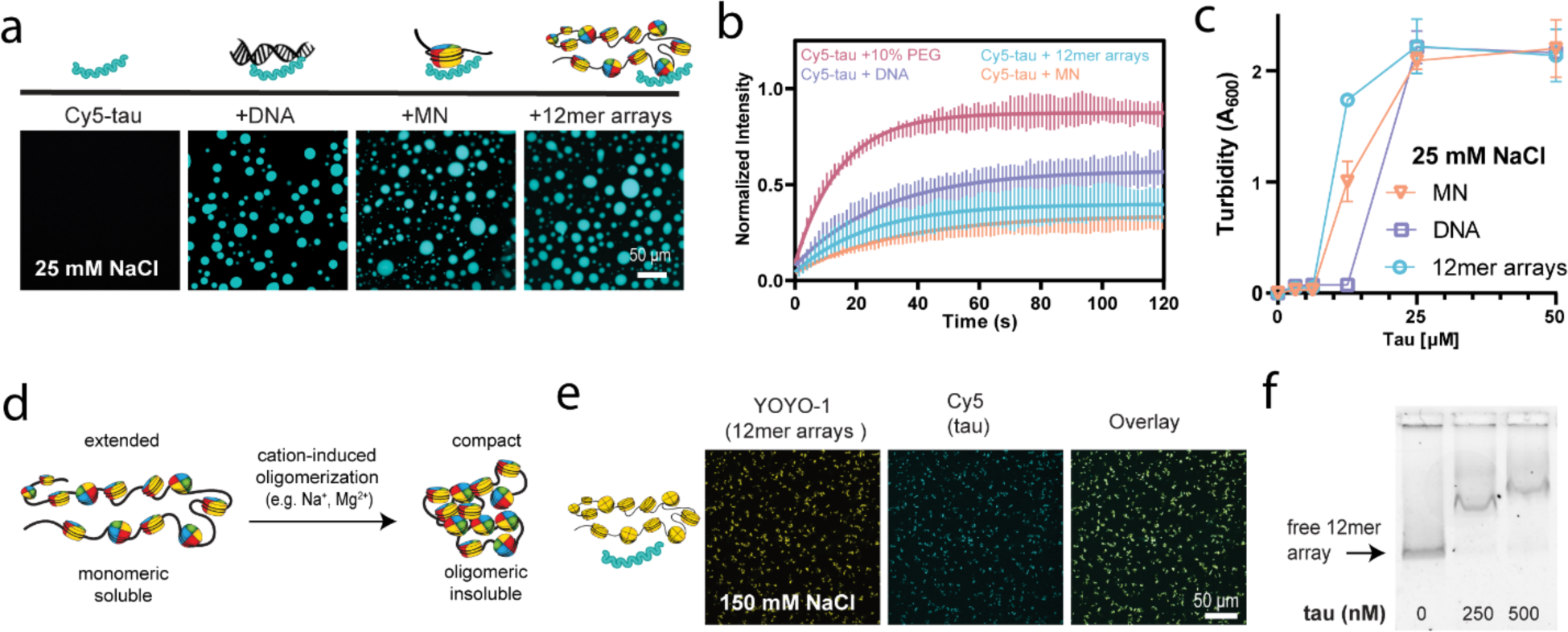
Tau undergoes phase separation and associates with 12mer arrays. a) Confocal microscopy images of droplets of 50 μM tau with 80 nM 12mer arrays (1 μM equivalent of mononucleosomes), 1 μM mononucleosomes, or 80 nM 2.1 kbp DNA (1 μM equivalent of ‘601’ DNA sites), visualized with 5% Cy5-tau (scale bar = 50 μm). Studies were conducted in 20 mM HEPES buffer, 25 mM NaCl, 0.5 mM TCEP, pH 7.2. b) FRAP analysis of 50 μM tau droplets with different LLPS inducers in the same low salt buffer. The standard deviation and the best monoexponential fit curve are plotted for each data set. Quantification of maximum recovery (%) and half-life (s) are tabulated in Table S1. c) Turbidity at A600 of tau solutions with different LLPS inducers in the same low salt buffer. d) Schematic of the compaction behavior of 12mer arrays in the presence of cations, including physiological salt. e) Confocal microscopy images of 50 μM tau (with 5% Cy5-labelled tau) with 80 nM 12mer arrays at 150 mM NaCl (scale bar = 50 μm) in the same buffer. YOYO-1 was added to visualize the arrays. f) EMSA of 4 nM 12mer arrays with and without tau, in 20 mM HEPES buffer, 150 mM NaCl, 0.5 mM TCEP, 0.1% Tween, pH 7.2. MN – mononucleosomes.

Within these droplets, we also sought to determine the mobility of different components using fluorescence recovery after photobleaching (FRAP) experiments. Although tau exhibited mobility characteristic of liquid droplet environments when phase separation was induced with crowding agent (10% PEG) similarly to published results,^4,11^ we found that nucleic acid-containing droplets showed relatively slow recovery after bleaching (**Fig. 1b**). While the maximum recovery percentage, or mobile fraction, was high in droplets induced by a crowding agent in the absence of DNA (∼87%), tau mobility was lower with DNA (∼57%), and even lower in the presence of nucleosomes (∼40% and 34% for 12mer arrays and mononucleosomes, respectively; values tabulated in **Table S1**). Similar low mobility has been observed in tau droplets induced with RNA,^56^ where it is known that tau binds tRNA with high affinity.^5^ These observations suggest that tau might interact strongly with DNA in the phase separated environment.

To compare the phase separation of tau with DNA, mononucleosomes, and 12mer arrays, we performed turbidity measurements using light scattering at 600 nm to quantify droplet formation across a variety of concentrations. Intriguingly, phase separation was induced at a lower tau concentration with 12mer arrays than with 2.1 kbp DNA (12 vs 25 μM tau) (**Fig. 1c**). This implies that the charge patterning of chromatin, with both positively charged histones and negatively charged DNA, enhances tau phase separation, and enables tau to phase separate at lower concentrations. Phase separation of tau with RNA has previously been observed at concentrations as low as 5 μM tau.^5,9^ It is unclear if the differences in LLPS critical concentration are due to the differences in our nucleic acid construct (length and sequence) and concentrations used, or an inherent preference for phase separation with RNA over DNA. Variation in behavior (extent of phase separation and droplet salt sensitivity) has been observed even among different types of RNA,^5^ and different studies use different tau constructs (e.g., 2N4R, and truncated versions), so we cannot directly make conclusions through these comparisons. Nonetheless, our data indicate that tau has the capacity to undergo LLPS with DNA at low salt and in the absence of a crowding agent, while mononucleosomes and nucleosome arrays enhance this transition.

### Tau associates with compact chromatin at physiological salt

While phase separation occurs under low salt conditions, it is also important to test physiological salt concentrations, which screen weak electrostatic interactions and typically lead to compact chromatin states. Under low salt conditions, 12mer arrays are monomeric and soluble, and present in an extended “beads-on-a-string” conformation with nucleosomes spaced by exposed DNA (**Fig. 1d**). Meanwhile, at higher salt (e.g.,150 mM NaCl), 12mer arrays undergo cation-induced oligomerization and form compact, higher order structures, with charge neutralization of DNA enabling closer packing of nucleosomes^57^ (**Fig. 1d, S4a-c**). This *in vitro* chromatin state is consistent with the solid-like behavior of highly compact chromatin in cells.^58^ While phase separated droplets could not be formed under these conditions, tau remained associated with the compacted 12mer arrays (**Fig. 1e**). We could similarly observe this with 12mer arrays compacted by Mg^2+^, an additional control for cation-induced oligomerization (**Fig. S4d**). This suggests that tau can remain associated with highly compacted chromatin, despite DNA being less exposed, and that this interaction is not screened by physiological salt concentrations. We verified that the observed results were not an artefact of using YOYO-1 by repeating key experiments with 12mer arrays made with fluorescein-labelled H2A (**Fig. S5**). In addition, we verified tau’s association with 12mer arrays at 150 mM NaCl using an electrophoretic mobility shift assay (EMSA) and showed that as low as 250 nM tau can induce a band shift of 12mer arrays, indicating a strong binding interaction (**Fig. 1f**). This co-association can be observed by microscopy with tau concentrations as low as 3 uM (**Fig. S3e**). This association is reversible, and tau can be removed from the 12mer arrays by the addition of naked DNA to the sample (**Fig. S6**).

While we cannot observe phase separation under these conditions, published studies indicate that crowding agents are necessary to induce tau LLPS at physiological salt. For example, tau-RNA LLPS can be screened by salt concentrations ranging from 25 to 100 mM NaCl,^5,9^ and LLPS at physiological salt (150 mM NaCl) requires the addition of PEG.^9^ Consistent with this, we observed that tau could phase separate with DNA at physiological salt only in the presence of crowding agent at concentrations as low as 6 uM tau (**Fig. S3b-e**), which is consistent behavior with the published literature. Meanwhile, tau remains fully associated but not phase separated with 12mer arrays at physiological salt in the presence of PEG, similar to its behavior without crowding agent (**Fig. S3b-e**). A complication to the use of crowding agents in our system is that the addition of PEG to 12mer arrays at high salt results in enhanced oligomerization. For instance, when centrifuged, 32% of 12mer arrays were soluble in 150 mM NaCl, and 17% when 10% PEG was also added (**Fig. S4a**). For this reason, we chose to avoid PEG altogether and to conduct most of our studies at low salt conditions, which gives us an opportunity to study a soluble and more accessible chromatin state and to focus on the interaction trends between tau, DNA, and nucleosome arrays. In summary, while tau cannot undergo LLPS with chromatin at physiological salt, it can reversibly bind and remain co-localized with compacted 12mer arrays, interactions that can have important consequences for DNA accessibility.

### Tau-chromatin binding and phase separation are DNA driven

We next sought out to characterize tau’s binding to DNA and mononucleosomes, to determine whether tau binds the linker DNA on 12mer arrays or can interact with the DNA wrapped around the histone octamer. To interrogate tau’s binding preferences, we compared tau’s binding to free DNA and mononucleosome constructs made with 147 bp and 177 bp DNA (147 bp mononucleosomes and 177 bp mononucleosomes, respectively) using a gel shift assay. 147 bp nucleosomes have the minimum amount of DNA to wrap around the histone octamer nucleosome core, whereas the 177 bp nucleosomes have 30 bp of ‘linker’ DNA. We did this comparison at low salt, since mononucleosomes at low concentrations can be unstable under high salt conditions^59^ (**Fig. S7a**), and we wanted to differentiate between nucleosome and free DNA binding. Since tau can bind DNA lengths of 13 bp or longer,^27^ we expected that multiple tau molecules would be able to bind, and consistent with this, we observed bands with increasingly larger mobility shifts corresponding to different tau: DNA or nucleosome stoichiometries (**Fig. 2**). There is precedent for observing cooperative binding of multiple tau molecules to RNA.^5^ In support of this observation, the apparent DNA-tau complex size continued to migrate higher at molar ratios of tau that far exceed the expected 1 tau to 13 bp of DNA and we achieved a higher quality curve fit when using a binding equation with a Hill slope coefficient. For that reason, we opted to fit the data to an equation for one site specific binding with a Hill Slope fitting for all constructs. While this fitting choice is a simplification, given that tau binding is non-specific and likely involves multiple binding events at multiple sites, we nonetheless present dissociation constants as an imperfect estimate to enable us to report on the observed trends.

**Figure 2.**
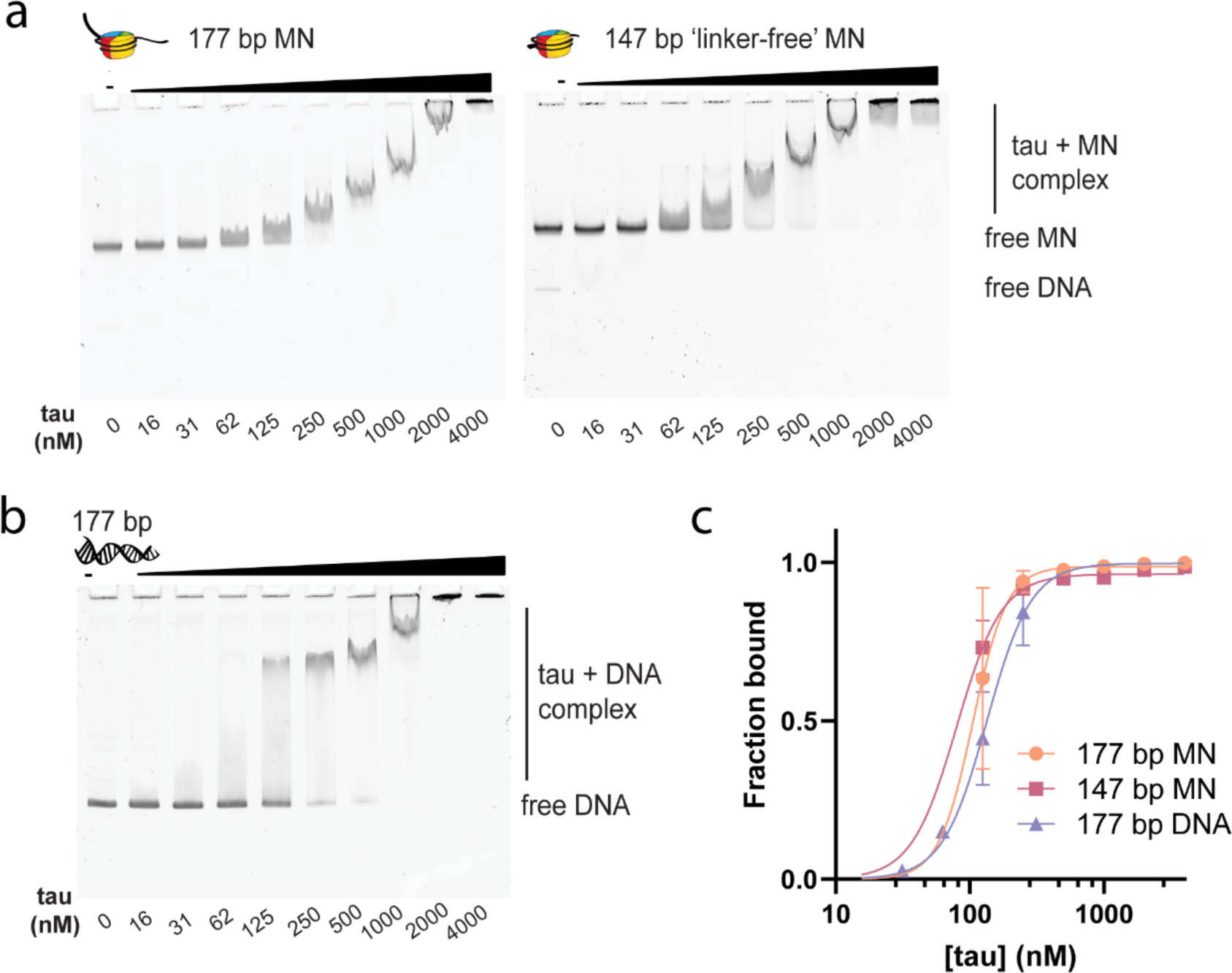
Tau binds DNA and nucleosomes. a) EMSA of 20 nM 177 bp mononucleosomes and 147 bp ‘linker-free’ mononucleosomes with various concentrations of tau, in 20 mM HEPES buffer, 10 mM NaCl, 0.5 mM TCEP, 0.1% Tween, pH 7.2. b) EMSA of 20 nM 177 bp DNA with various concentrations of tau, in the same low salt buffer. c) Quantification of binding propensity based on the intensity of the unbound DNA or mononucleosome bands in Fig. 2a, b. Error bars represent the standard deviation from three independent EMSA experiments. 177 bp DNA, 147 bp mononucleosomes, and 177 bp mononucleosomes were quantified in triplicate for the following data points: 0, 125, 250, 500, and 1000 nM. 16-62 nM data points were excluded for the mononucleosome binding due to the smearing and complex bands overlapping with the unbound probe. Curves were fit to a one site specific binding with Hill Slope equation. MN – mononucleosomes.

Comparison of these constructs revealed that tau can bind 177 bp DNA, 147 bp mononucleosomes, and 177 bp mononucleosomes with similar affinity, with equilibrium dissociation constant (Kd) values of 134.6 ± 8.7, 83.4 ± 13.0, and 105.5 ± 14.9 nM, respectively (**Fig. 2c**). This may indicate a small preference for mononucleosomes over DNA, implying that the positive charge of histones enhances tau’s binding. However, we interpret this difference with caution, due to the greater challenge of quantifying the band intensities of lower tau concentration data points in mononucleosome EMSAs. Significantly, tau’s ability to efficiently bind linker-free mononucleosomes reveals that tau can still access DNA wrapped around the nucleosome core. This explains tau’s ability to associate with highly compacted chromatin states in the presence of high salt and Mg^2+^ concentrations.

We verified that tau could bind DNA at physiological salt, yielding a Kd value of 188.3 ± 9.7 nM for 177 bp DNA (**Fig. S7b**). We also verified this binding on a 22 bp DNA previously shown to bind tau and expected to interact with one-to-one stoichiometry,^25^ and observed a Kd of 112.3 ± 3.7 nM by EMSA (**Fig. S7c).** We independently confirmed a low nanomolar Kd of 6.3 ± 0.5 nM for 22 bp DNA by fluorescence anisotropy (**Fig. S7d**). These values are consistent with published literature. For comparison, previously reported *Kd* values include: ∼ 40 nM by surface plasmon resonance,^60^ and 300 to 900 nM (DNA sequence dependent) by kinetic capillary electrophoresis methods.^24^ In addition, tau is known to bind tRNA with a dissociation constant of 460 nM by EMSA and 735 nM by isothermal titration calorimetry.^5^

Tau’s strong affinity for DNA led us to determine the influence of tau’s DNA-binding regions (DBRs) on DNA binding and phase separation. To this aim, we prepared constructs of tau lacking the residues within the DBRs identified to be important for binding by previous NMR titration studies.^25^ These studies show that the DNA binding interaction occurs through simultaneous binding of the second half of tau’s PRD (R209-A246) and MTBD R2 (K267-S289) of 2N4R tau, and that both regions can independently bind DNA with equivalent affinity.^25^ To test the impact of these independent DBRs, we prepared tau constructs lacking the regions with the strongest chemical shift perturbations within the PRD (ΔPRD-DBR; T190-K210, 1N4R nomenclature) and MTBD (ΔMTBD-DBR; G241-K264, 1N4R nomenclature) (**Fig. S8a**). Both constructs reduced the binding affinity for 177 bp DNA to a similar extent (**Fig. S8b, c**), resulting in binding affinities of 410 ± 50 nM (ΔPRD-DBR) and 430 ± 150 nM (ΔMTBD-DBR). More intriguingly, however, these deletions had very different effects on phase separation propensity. The MTBD-DBR-deletion significantly reduced the degree of LLPS with 12mer arrays, while the PRD-DBR-deletion abrogated phase separation (**Fig. S8d, e**). This revealed an unexpected degree of specificity to tau’s phase separation, most likely driven by the net positive charge of each DBR. The PRD-DBR deletion yielded a net charge of +1, while the MTBD-DBR deletion has a charge of +2, both lower than full-length tau’s charge of +6. Notably, neither deletion reduced tau’s phase separation behavior in the presence of crowding agent (**Fig. S8e**). This shows that tau’s phase separation with 12mer arrays can be regulated by tau’s DNA binding affinity and charge. In summary, we show that tau has a strong affinity for DNA and mononucleosomes and can bind DNA directly on the nucleosome surface. In addition, reducing tau’s ability to bind DNA can disrupt phase separation.

### Low micromolar concentrations of tau compact open chromatin

Given the nanomolar affinity of tau for DNA and mononucleosomes, we were curious what effect low micromolar concentrations of tau have on 12mer arrays, and whether tau has an influence on the formation of higher order chromatin structures. We performed these studies at low salt, where 12mer arrays are open, extended, and soluble. To assess the effect of tau on the oligomerization state of chromatin, we first examined the architecture of 12mer arrays using negative-stain transmission electron microscopy (TEM) images. As expected, 12mer arrays presented the characteristic ‘beads-on-a-string’ extended fiber conformation (**Fig. 3a**). As a control for compaction, low Mg^2+^ concentrations were added, and the 12mer arrays underwent cation-induced oligomerization and formed compacted structures with decreased inter-nucleosome distances. Higher concentrations of Mg^2+^ and physiological salt were avoided, because the oligomerized 12mer arrays become large enough to be visible by microscopy (**Fig. S4b-d)** and cannot efficiently adhere to the TEM grid. We found that the 12mer arrays also adopted a compact configuration when incubated with tau (**Fig 3a**). A molar ratio of 1 tau per nucleosome (0.136 μM tau) resulted in reduced inter-nucleosome distances within the array, and with ten molar equivalents (1.36 μM tau), tau produced oligomerized chromatin structures reminiscent of the Mg^2+^-compacted control (**Fig. 3a**), suggesting that tau can compact chromatin in a concentration dependent manner. This is not unexpected, given tau is cationic, and cation-DNA charge neutralization is a known mechanism to enable closer nucleosome packing. These concentrations are lower than those of tau phase separation but above the Kd, implying that compaction may be a property of tau binding to chromatin.

**Figure 3.**
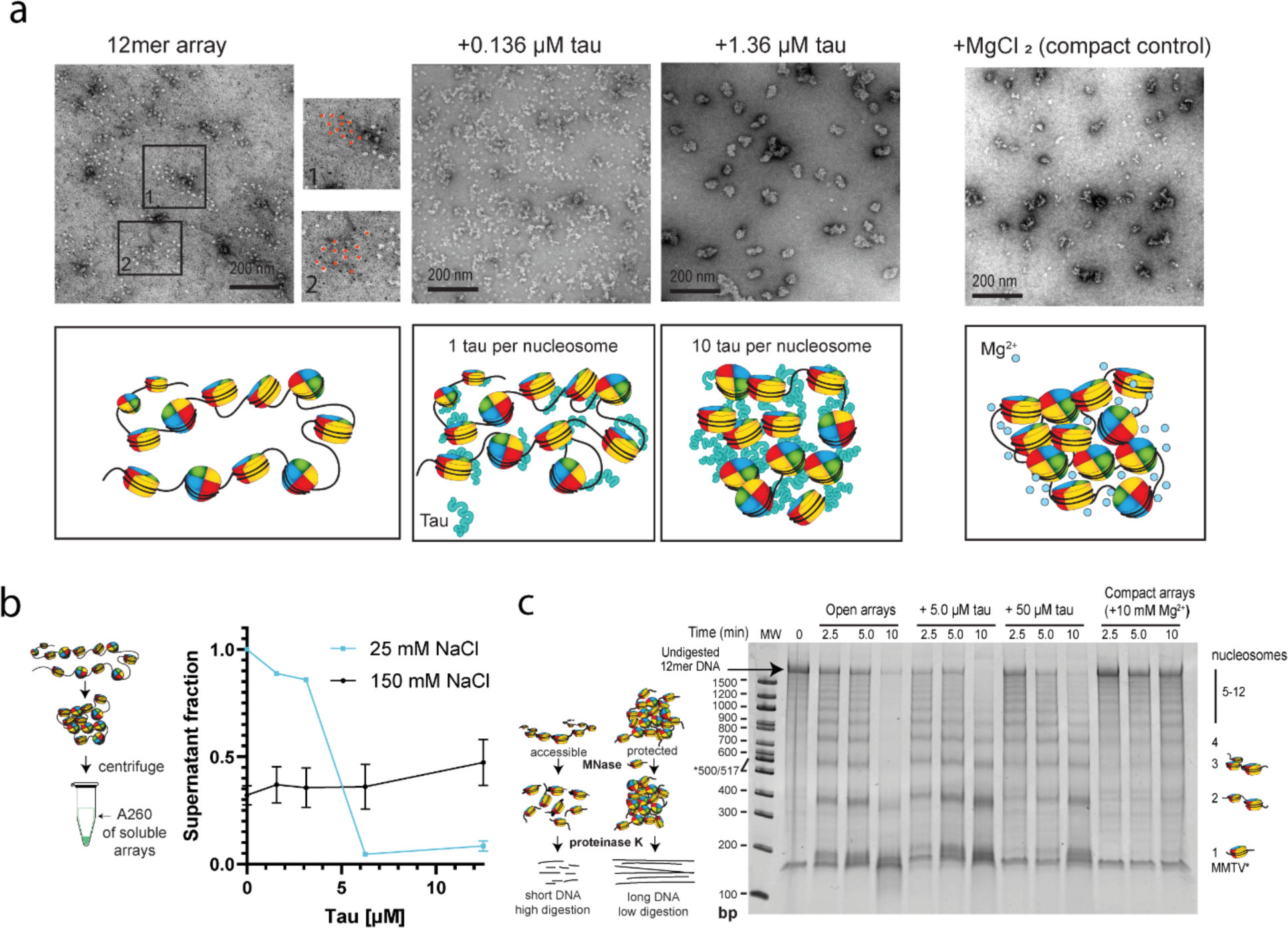
Tau can compact open chromatin under low salt conditions. a) TEM images and schematic of cation-induced compaction of 12mer arrays (11.3 nM of 12mer array which is equivalent to 136 nM of mononucleosomes) with MgCl2, as compaction control, and increasing concentrations of tau (0.136 μM and 1.36 μM), conducted in 20 mM HEPES buffer, 10 mM NaCl, 0.5 mM TCEP, pH 7.2. b) (left) Schematic of the chromatin oligomerization assay. 12mer arrays were incubated in different buffer conditions, subjected to low-speed centrifugation, and the A260 of the supernatant was measured. The soluble fraction reflects the degree of self-association of the 12mer arrays. (right) Increasing concentrations of tau were incubated with 30 nM 12mer arrays (for an A260 ∼ 1) in different salt conditions in the same buffer. c) (left) Schematic of the MNase digestion assay to probe for DNA protection within 12mer arrays. (right) Time course of MNase digestion of 2 nM 12mer arrays with varying concentrations of tau and MgCl2, in the same buffer. All lanes contain a band at ∼ 150 bp which results from a small amount of MMTV nucleosomes present in the sample.

To further confirm this effect, we used differential centrifugation-based oligomerization assays. When 12mer arrays oligomerize, they become insoluble enough to pellet under low-speed centrifugation, and the degree of oligomerization can be measured by the decrease in the fraction of 260 nm absorbance in the supernatant.^61^ Interestingly, at low salt, tau concentrations below the critical concentration for phase separation were sufficient to cause significant array oligomerization (**Fig. 3b**). All 12mer arrays transitioned to the pellet fraction at 3 to 6 μM tau, which is slightly below the critical concentration (6 - 12 μM) required for phase separation as determined by our turbidity assay (**Fig. 1c**). While no droplets were visible by microscopy at these tau concentrations, we cannot exclude that the transition to the pellet represents “mesoscopic” phase separation, which has previously been observed by dynamic light scattering experiments at tau concentrations below the critical concentration for LLPS.^9^ On the other hand, at 150 mM NaCl, tau could not promote further self-association of 12mer arrays, despite being associated with arrays according to microscopy (**Fig. S3a,e)** and our earlier EMSA experiments (**Fig. 1f**). This suggests that tau cannot further oligomerize 12mer arrays that are already self-associated due to the presence of cations.

We next sought to survey the effect of this oligomerization on array accessibility by assaying digestion with Micrococcal Nuclease (MNase), a non-specific nuclease used to monitor chromatin accessibility (**Fig. 3c**). Since tau strongly binds DNA (**Fig. 2**) and low concentrations of tau appear to promote oligomerization (**Fig. 3a, b**), we hypothesized that tau may protect linker DNA. MNase would favor digesting the unprotected linker DNA between nucleosomes first, revealing whether chromatin is open and accessible to being digested into shorter segments (predominately mono- and dinucleosomes) or compacted and protected as longer multi-nucleosome species. This digestion is followed by treatment with proteinase K to remove the histone octamers from the DNA. The digested DNA produces multiple bands on a native gel, reflecting the different sizes of nucleosome polymers (mono- to 12mer). When 12mer arrays were incubated with excess (5 and 50 μM) tau over multiple time points, the DNA digestion showed that tau reduced the accessibility of DNA (**Fig. 3c**). The presence of tau enhanced the intensity of bands corresponding to larger fragments and increased the size of the smallest observed fragment, suggesting that tau protects linker DNA and can compact chromatin. This compaction effect is modest relative to array compaction with 10 mM Mg^2+^, which causes chromatin oligomers to fully precipitate out of solution and is a common control for compaction. There was not a major difference between 5 μM, a concentration that shows high self-association of 12mer arrays prior to phase separation (**Fig. 3b**), and 50 μM, a phase separation concentration **(Fig. 1c)**, which may imply that this effect is independent of phase separation. Overall, we observe that tau can promote modest compaction of chromatin at concentrations well below the critical concentration for phase separation.

### Tau has a subtle effect on DNA contact points in the nucleosome core

Tau’s ability to bind and protect DNA within 12mer arrays made us wonder whether it also has a specific structural effect on chromatin. In addition, recent pulldown studies have shown that tau can bind the disordered tails of histones H3 and H4 and has preferences for unmodified H3 and highly acetylated H4,^62^ suggesting some degree of specificity in nucleosome interactions. Detailed residue-specific structural information, specifically about the influence of tau on the nucleosome core and histone tails can be obtained using magic angle spinning solid-state NMR spectroscopy (MAS NMR). MAS NMR has no size limit, making it well-suited to studying the megadalton 12mer array complex.^63,64^ To perform MAS NMR studies, we prepared 12mer arrays containing ^13^C,^15^N-labeled H4, while histone proteins H3, H2A, and H2B remained unlabeled. In this way, the only visible species by NMR within the tau-12mer array assembly was H4, serving as a reporter of structural changes upon addition of tau. We chose H4 as it is the smallest histone protein, it yields ^13^C-^13^C spectra with high resolution, and solid-state assignments are available for the residues within the nucleosome core.^63^ We used these proteins to reconstitute ∼ 6 mg of 12mer arrays, which were precipitated with 6 mM MgCl2, and then combined with 20 mg of tau (∼18 molar equivalents per nucleosome), the same preparation sequence we used for our imaging experiments. It should be noted that Mg^2+^ also has a beneficial effect on the quality of chromatin spectra and previous studies have used up to 20 mM MgCl2 to yield ^13^C-^13^C correlations with excellent resolution.^63,64^ Here, we chose a concentration of 6 mM in order to facilitate rotor packing and obtain good resolution, without compromising tau’s ability to co-partition with 12mer arrays (**Fig. S4**). We then proceeded to record MAS NMR correlation spectra of the 12mer arrays with isotopically labeled H4 in the presence and absence of tau (parameters described in **Fig. S9a**).^65–67^ One of the advantages of MAS NMR is that it allows dynamics-based spectral editing of mobile and rigid components in the same large molecular weight sample.^68^ For example, scalar based experiments such as 1D and 2D ^1^H-^13^C INEPT can be employed to detect the dynamic H4 tail, while dipolar based pulse schemes such as 1D ^1^H-^13^C cross-polarization (CP) and 2D ^13^C-^13^C DARR can be used to characterize the H4 residues that participate in the rigid nucleosome core. For H4 in the context of the nucleosome, residues 1-25 typically appear in the INEPT spectra,^69^ while residues 26 and beyond are represented in the CP and DARR spectra (**Fig. 4a**).

**Figure 4.**
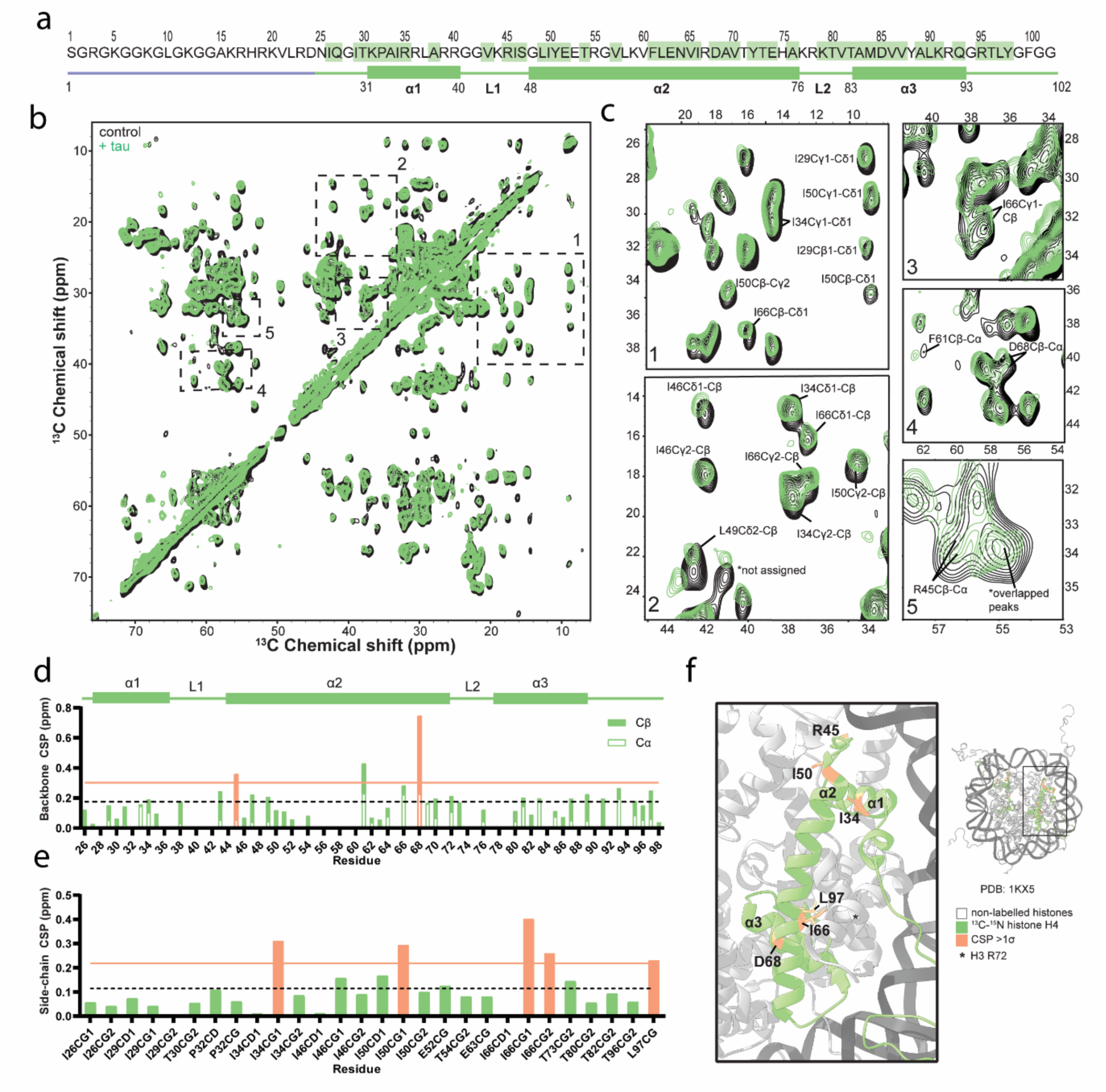
Tau leads to subtle perturbations at DNA contact points on the nucleosome core. a) Schematic of the sequence and secondary structure of histone H4. The residues highlighted in green display resolved correlations that were used in the analysis described below. Rigid regions that appear in the DARR spectrum are shown as green rectangles and lines, while the mobile regions of the INEPT spectrum are shown in purple. b) 2D ^13^C-^13^C DARR spectrum recorded with 20 ms mixing time on labelled histone H4 within 12mer arrays (black) and with 20 mg of added tau (green), in 10 mM Tris, 0.1 mM EDTA, 10 mM KCl, pH 7.5. Conditions were chosen to match those for published chemical shift assignments.^63^ c) Zoomed in regions of the 2D DARR spectrum, showing the largest quantified perturbations. d) Summed chemical shift perturbations (CSPs) of Cα and Cβ chemical shifts per residue, obtained from single-bond correlations. The bars are split into the contribution from the Cα (white) and Cβ (green) CSP. The dotted line is the average value along the sequence, and the orange line is one standard deviation above the mean. Orange shaded bars are residues that are one standard deviation above the mean. e) CSP of side-chain carbons obtained from single bond correlations. The dotted line is the average value along the sequence and the orange line is one standard deviation above the mean. Orange shaded bars are residues that are one standard deviation above the mean. f) Perturbations plotted onto the nucleosome structure. Histone H4 is shown in green and residues containing at least one atom with a CSP above one standard deviation above the mean are shaded in orange and labeled.

In the presence of tau, the majority of the CP-DARR spectrum overlayed with the control spectrum, indicating no major structural rearrangements in the rigid parts of the nucleosome (**Fig. 4b,c**). As a quantitative metric for perturbation, we summed the chemical shift perturbations (CSP) of Cα and Cβ chemical shifts obtained from single-bond correlations (**Fig. 4d**) while the CSPs of individual side-chain atoms were plotted separately (e.g., Cγ, Cδ) (**Fig. 4e**). Long-range correlations (e.g., Cα-Cγ, Cα-Cδ) were excluded from the quantitative analysis, since our experiments were not optimized for acquiring them. Subtle perturbations that were greater than one standard deviation above the mean were observed at the backbone of Arg45 and Asp68 and the sidechains of Ile34, Ile50, Ile66, and Leu97 (**Fig. 4c-e**). Despite being above one standard deviation above the mean, we excluded Tyr61 due to the weak intensity of the Cα and Cβ crosspeak. The observed perturbations loosely map to known histone-DNA contacts observed in the crystal structure of the nucleosome core (PDB: 1KX5^70^). The upper region of perturbation includes Arg45, which has contacts with the DNA backbone, and the lower region is in proximity to H3 DNA contact Arg72. The perturbations support the idea that tau interacts with nucleosomal DNA without capacity for histone recognition or influencing the nucleosome core. The INEPT spectrum, which shows the residues of the mobile histone tail, displays minimal perturbations and no evidence of any specific interaction with tau (**Fig. S9b-d**). This is interesting considering a recent study showing that tau binds unmodified H3 and H4,^62^ but those results do not seem to hold in the context of nucleosomes, where tau’s high DNA-binding affinity favors interactions with nucleosomal DNA. In addition to chemical shifts, we analyzed the intensity differences in both the DARR and INEPT spectra (**Fig. S9d-f**). Although there are observable peak intensity changes in the spectra for some residues with low signal-to-noise ratios (SNR), once SNR is considered, these changes do not appear to be statistically significant. This implies that there are no significant differences in the dynamics of the nucleosome core or the H4 tail in the presence of tau.

### Tau phosphorylation reduces binding and disrupts phase separation with 12mer arrays

Having established a strong affinity of tau for DNA and nucleosomes, we sought out to determine how the interactions are modulated by phosphorylation. Numerous reports indicate the existence of phosphorylated tau in the nucleus.^14,17,28,42,71,72^ Phospho-site specific antibodies and mass spectroscopic (MS) studies revealed that nuclear tau has a specific phosphorylation signature (including T181, T212, S404; 2N4R nomenclature),^28^ and other reports have identified T212/S214 phosphorylation (recognized by the AT100 antibody) to be abundant in chromatin.^17,72^ These phosphorylation signatures are distinct from pathological ones and are speculated to be functional. In contrast, tau hyperphosphorylation is a well-known modification in AD that may result in loss of function in the nucleus. On average, tau is phosphorylated at 2-3 sites in healthy brains and at 5-9 sites when aberrantly hyperphosphorylated in NFTs.^73^ Phosphorylation is known to reduce or abrogate tau’s DNA binding affinity^25,60,74^ and to increase LLPS propensity with crowding agents.^6,8,11^ Lastly, there are reports that the levels of modestly phosphorylated nuclear tau (at the T212/S214 sites) increase with healthy aging and that AD progression correlates with an increase of nuclear tau phosphorylated at S202/T205 (AT8 antibody epitope) and culminates with the exit of tau from the nucleus.^41,42^ It is unclear whether uncoupling of tau from chromatin contributes to epigenetic dysregulation, but in either case, this also strongly implies that phosphorylated tau may have a functional role in the nucleus which gets disrupted by hyperphosphorylation, highlighting the need to understand the interaction of phosphorylated tau with chromatin *in vitro*.

AD and nuclear relevant phosphorylated forms of tau can be produced by incubation with tau’s kinases, including MAPK, cdk5, CAMKII, GSK-3β, and PKA.^75^ Here, we used *in vitro* phosphorylation by PKA and GSK-3β to generate phosphorylated tau species. PKA is known to prime tau for phosphorylation by GSK-3β, which promotes the phosphorylation of numerous AD-associated sites and allowed us to produce hyperphosphorylated tau. Tau was incubated with PKA, GSK-3β, or both for 16 hours, and phosphorylation was confirmed by the upward shift observed by SDS-PAGE (**Fig. 5a**). Intact mass spectrometry (MS) analysis shows that phosphorylation by PKA (PKA-ptau) or GSK-3β (GSK-ptau) alone leads to a heterogenous distribution of 2-5 or 3 phosphorylation events, respectively, while incubation with both kinases (PG-ptau) leads to a hyperphosphorylated state with 9-11 phosphorylated sites (**Fig. 5b, Fig. S10**). Thus, GSK and PKA-phosphorylated tau can provide a model for minimal phosphorylation, while PKA-GSK-phosphorylated tau can serve as a model for hyperphosphorylation. To better characterize the phosphorylated forms, we performed post-translational mapping by trypsin digestion and sequencing using liquid chromatography-tandem mass spectrometry (LC-MS/MS). Tau was phosphorylated at multiple sites which exceeded the average number of detected phosphates (**Fig. S11**), confirming the existence of a heterogenous phosphorylation state (tabulated in **Table S3**). Phosphorylation specificity was low and different sites were detected in independent runs (**Table S3**). For ease of comparison to the bulk of the literature, we will discuss phosphorylation sites using the residue numbering of the 2N4R isoform (1N4R numbering tabulated in **Table S3**). In hyperphosphorylated tau (PG-ptau), MS detected significant phosphorylation throughout the sequence, in the PRD and C-terminus (**Fig. S11, Table S3)**. This includes the nuclear-associated T181, T212, S214, and S404, and NFT-associated S202/T205 and S396/S404 sites, suggesting that it can serve as a model for AD-associated hyperphosphorylated tau.^3,17,28,72,76^ Meanwhile, phosphorylation was detected in the PRD and the C-terminus in both PKA-ptau and GSK-ptau, and sporadically in the MTBD (**Fig. S11**).

**Figure 5.**
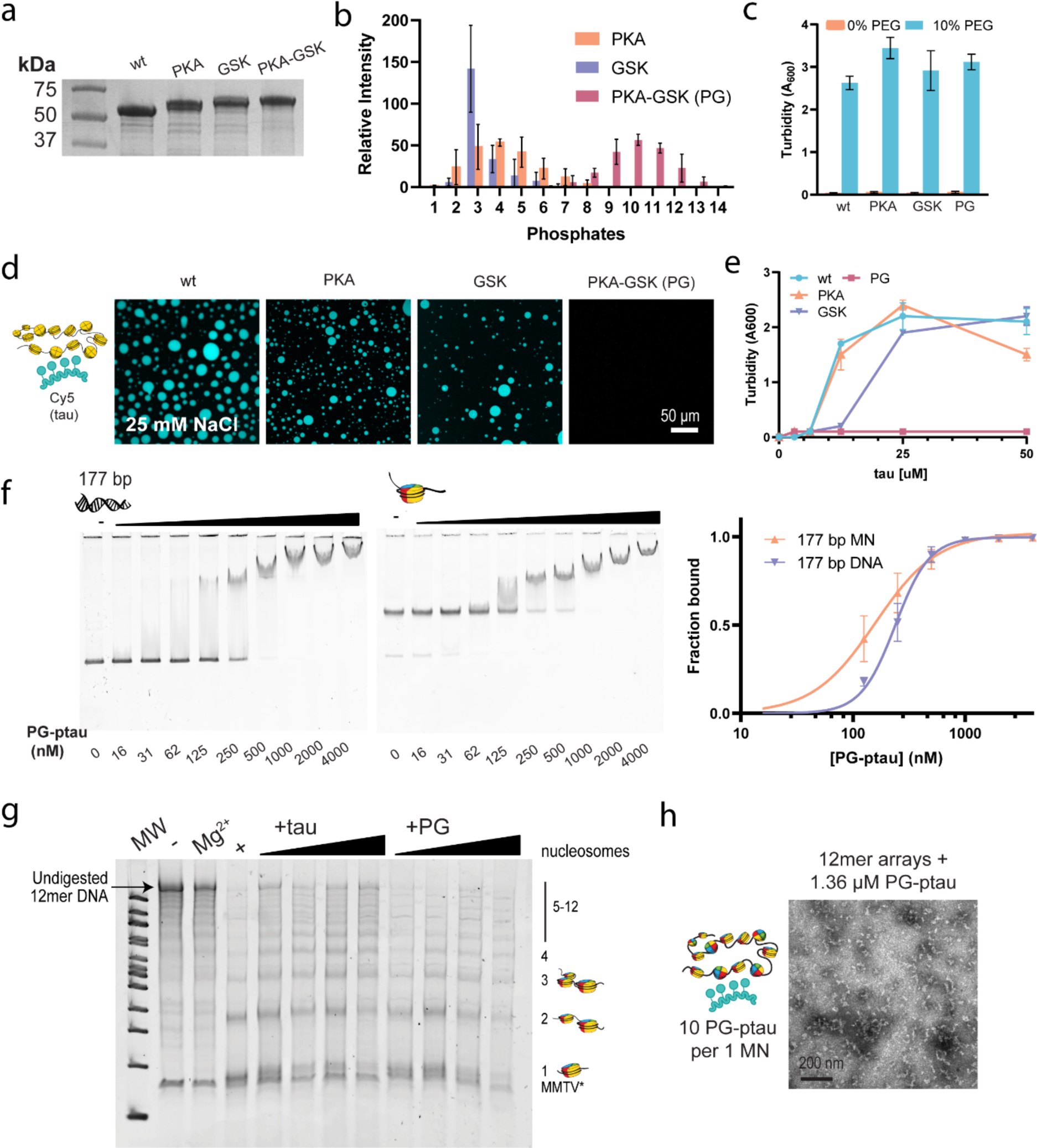
Phosphorylation disrupts tau’s phase separation and compaction of 12mer arrays. a) Gel analysis of tau phosphorylation using PKA, GSK or both (full gel image presented in Fig. S10a). b) Abundance of different masses corresponding to the number of phosphates for each phosphorylated form as detected by intact MS analysis. Mass values were binned to the nearest whole number of phosphates. Error bars reflect standard deviation resulting from independent phosphorylation reactions (n=3 for PKA and GSK tau or 4 with PKA-GSK tau). Individual replicates and non-binned values can be found in Fig. S10b. c) Turbidity (A600) analysis of wild-type and phosphorylated tau constructs, with LLPS induced by 10% PEG-6000, in 20 mM HEPES buffer, 25 mM NaCl, 0.5 mM TCEP, pH 7.2. d) Confocal microscopy images of liquid droplets of 50 μM PKA, GSK or PKA-GSK phosphorylated tau with 12mer arrays, spiked with 5% Cy5-tau (control) or Cy5-ptau (phosphorylated samples), in the same low salt buffer. YOYO-1 was added to visualize the 12mer arrays (scale bar = 50 μm). e) A600 turbidity measurements of the phase separation of PKA, GSK or PKA-GSK phosphorylated tau with 12mer arrays as function of tau concentration. Studies were conducted in the same low salt buffer. f) (left) EMSA to assess the binding propensity of PKA-GSK phosphorylated tau with 20 nM 177 bp DNA, or mononucleosomes in the same buffer containing 10 mM NaCl. Representative gel images from three independent experiments are shown. (right) Quantification of binding propensity based on the intensity of the unbound DNA or mononucleosome bands. Error bars represent the standard deviation from three independent EMSA experiments. g) MNase digestion of 12mer array samples containing varying concentrations of wild-type or PKA-GSK phosphorylated tau. (-) is an undigested control, the lane with Mg^2+^ controls for the effect of compaction, and (+) is a positive control for complete digestion. Protein concentrations used in the gradient are 5, 10, 25, and 50 μM. Studies were conducted in the same buffer containing 10 mM NaCl. h) TEM image of 12mer arrays (11.3 nM, or 136 nM equivalent of mononucleosomes) with 1.36 μM PKA-GSK phosphorylated tau, in the same buffer containing 10 mM NaCl. MN – mononucleosomes; PG – PKA-GSK phosphorylated tau; MW – molecular weight.

We first tested the effect of phosphorylation on tau’s ability to undergo LLPS. Similar to previous reports,^6^ phosphorylated tau could phase separate extensively in the presence of crowding agent (**Fig. 5c**). We then tested how phosphorylation affected tau’s phase separation with 12mer arrays, using Cy5-labelled phosphorylated tau. Minimal phosphorylation was mostly non-perturbative (**Fig. 5d, e**), and similar to non-phosphorylated tau, 12mer arrays partitioned into the droplets (**Fig. S12a**). Meanwhile, hyperphosphorylation completely abrogated phase separation. This was not due to aggregation or loss of phase separation propensity, as phosphorylated tau phase separated extensively when induced with crowding agent (**Fig. 5c**). Hyperphosphorylated tau (PG-ptau) maintained a relatively strong affinity for DNA and mononucleosomes (**Fig. 5f**). In EMSAs, the binding affinities for 177 bp DNA and 177 bp mononucleosomes were 241 ± 14 (∼2X wt tau), and 166 ± 28 nM (∼2X wt tau), respectively (**Fig. 5f**). Similarly, the 22 bp DNA fragment showed lower affinity to DNA than non-phosphorylated tau, with a Kd of 600 ± 160 (∼5X wt tau) by EMSA **(Fig. S12b)** and 76 ± 16 nM (∼13X higher than wt tau) by fluorescence anisotropy (**Fig. S12c**). We verified that binding to DNA could occur at physiological salt and obtained a Kd of 426 ± 40 nM for 177 bp DNA at 150 mM NaCl (**Fig. S12d**).

While these measurements reveal a reduction in binding affinity for PG-ptau, the Kd values remain in the nanomolar range and represent relatively strong interactions with DNA. This is not surprising, given that the post-translational modification mapping detects little phosphorylation in tau’s DNA-binding MTBD region (**Fig. S11a**). To the best of our knowledge, the affinity of a hyperphosphorylated species of tau (9-11 phosphates) prepared by incubation with PKA and GSK to DNA has not been reported before. However, previous reports have explored the binding of various forms of phosphorylated tau to DNA. For example, tau phosphorylated by neuronal cdc2-like kinase (NCLK) ^26^ and in Sf9 insect cells^60^ can still bind DNA to a reduced degree (neither affinity quantified). The Sf9 phosphorylated tau study did not report the degree of phosphorylation, but the modification protocol would be expected to yield 12 phosphorylation sites,^77^ potentially comparable with the number of modifications detected in our study. Meanwhile, tau phosphorylated by mouse brain extract kinases (18 phosphate modifications),^25^ CDK2/CycA3 kinase (2 phosphate modifications),^25^ and GSK (not quantified in study)^74^ cannot bind DNA. These phospho-forms of tau vary significantly by the site and extent of phosphorylation, and different DNA constructs were used, which may explain the degree of variation in literature reports.

We next investigated the functional consequences of phosphorylation on the observed compaction behavior of tau. As shown above, wild-type tau showed increased protection of 12mer array linker DNA in a concentration dependent manner (**Fig. 3c**). In contrast, hyperphosphorylated tau (PG-ptau) displayed a reduced ability to protect linker DNA against digestion with MNase (**Fig. 5g**). In addition, the TEM images show a similar loss in the ability to oligomerize and condense 12mer arrays (**Fig 5h**), suggesting a reduced ability to promote a more compact chromatin state. Hyperphosphorylated tau, although capable of binding mononucleosomes, does not appear to efficiently neutralize DNA charge to promote closer nucleosome packing. In summary, our results show that minimal phosphorylation by PKA or GSK does not dramatically alter the phase separation properties of tau and 12mer arrays. Hyperphosphorylation by using both PKA and GSK, on the other hand, abrogates phase separation and reduces protection of linker DNA. Intriguingly, however, hyperphosphorylated tau still retains a relatively strong ability to bind DNA and mononucleosomes.

### Tau undergoes LLPS with phosphorylated HP1α

Given tau’s strong propensity to interact with chromatin, without apparent DNA or histone binding specificity, we wondered what role protein-protein interactions could play in tau’s localization patterns within the nucleus. One major model for heterochromatin organization suggests that heterochromatin domains form through phase separation of heterochromatin protein 1α (HP1α), and that droplets separate chromocenters from the surrounding euchromatin, keeping out transcriptional machinery and forming a selectively permeable boundary.^47,49,78^ This LLPS process appears to be electrostatically driven and can be modulated by a number of HP1α binding partners.^47,49,51,53,78–80^ Tau’s capacity to phase separate through an electrostatic mechanism and the observation that its depletion or pathology disrupts HP1α and heterochromatin clustering^20,31,33^ prompts the question of a possible interplay between the phase separation of tau and HP1α. To explore this possibility, we reconstituted the heterochromatin environment *in vitro* by combining tau with phosphorylated HP1α (pHP1α, **Fig. S13**), in the presence and absence of 12mer arrays (**Fig. 6a**). We chose to work with HP1α phosphorylated on its N-terminal extension as this modification significantly reduces the saturation concentration required for LLPS *in vitro* in the absence of nucleic acids or chromatin from > 250 μM for unmodified HP1α to ∼ 75 μM for pHP1α (concentrations compared at 75 mM KCl).^79^ Literature suggests that HP1α is phosphorylated by the CK2 kinase and that it is constitutively present in cells, pointing to its biological significance.^47,79,81^ Suitably phosphorylated HP1α can also be produced recombinantly in *E. coli* using co-expression with the CK2. Upon combination of 50 μM tau and 50 μM pHP1α at low salt, the two instantly formed droplets, and 12mer arrays partitioned into these droplets when present (**Fig 6a**). Under these conditions pHP1α does not phase separate alone or with 12mer arrays (**Fig. S14a**), suggesting that tau reduces the pHP1α saturation concentration required for LLPS. The phase separation of tau and pHP1α appears to be driven by interactions of tau’s positively charged regions with the negatively charged phosphates in the N-terminal tail of pHP1α, since tau cannot phase separate with non-phosphorylated HP1α at similar concentrations (**Fig. S15**). The co-localization of tau, pHP1α, and 12mer arrays is maintained at physiological salt where oligomeric chromatin is observed (**Fig. 6b**, overlay in **Fig. S14b**). At 25 mM NaCl, the tau-pHP1α droplets were confirmed to be liquid-like by FRAP experiments (**Fig. 6c**). Under these conditions, tau remained highly mobile, with a maximum recovery percentage of ∼82%, which is comparable to the ∼87% maximum recovery in tau droplets induced by crowding agent. In the presence of pHP1α and 12mer arrays, tau displayed lower recovery at 49%, which is slightly higher than the 40% recovery observed in droplets of tau and 12mer arrays alone. Meanwhile, at 150 mM NaCl, tau does not appear to display liquid-like mobility according to our FRAP experiments (**Fig. S14c**).

**Figure 6.**
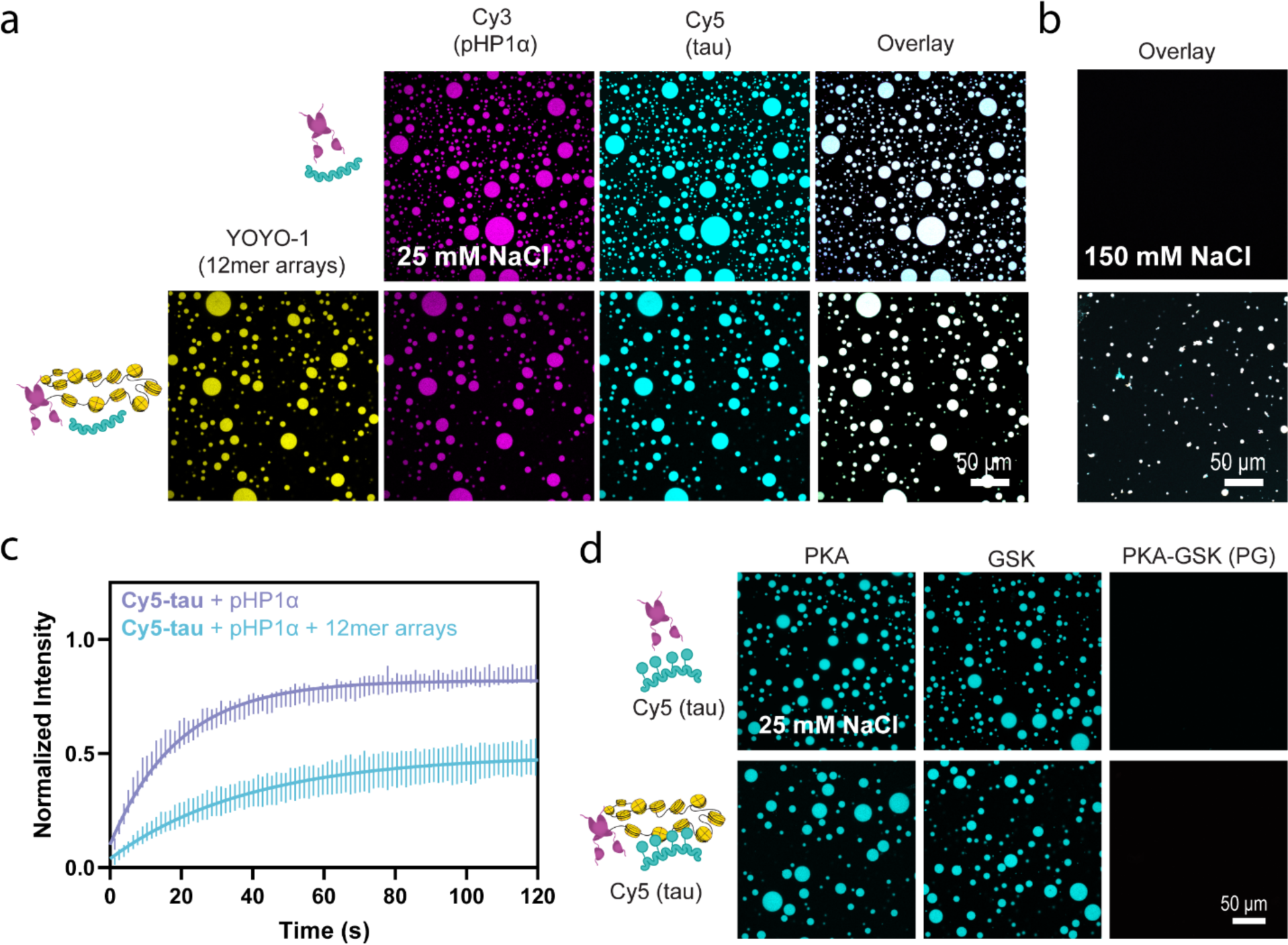
Phase separation of tau with pHP1α and 12mer arrays. a) Confocal microscopy images of droplets of 50 μM tau and 50 μM pHP1α, with and without 80 nM 12mer arrays, imaged using 5% Cy5-tau, 5% Cy3-pHP1α, and YOYO-1 (scale bar = 50 μm), in 20 mM HEPES buffer, 25 mM NaCl, 0.5 mM TCEP, pH 7.2. b) Co-localization of tau, pHP1α, and 12mer arrays in the same buffer containing 150 mM NaCl. See Fig. S14b for the full overlay. c) FRAP analysis of tau mobility in samples containing 50 μM tau with pHP1α alone (purple) or pHP1α and 12mer arrays (blue) in the same low salt buffer. The standard deviation and the best monoexponential fit curve are plotted for each data set. Quantification of maximum recovery (%) and half-life (s) are tabulated in Table S1. d) Confocal microscopy images of 50 μM pHP1α and 50 μM PKA, GSK or PKA-GSK phosphorylated tau (top) and 50 μM pHP1α, 50 μM phosphorylated tau, and 80 nM 12mer arrays (bottom), in the same low salt buffer.

The phase separation of tau and pHP1α alone is robust and is maintained at 100 mM NaCl but screened by 150 mM NaCl (**Fig. S14d**). A phase diagram revealed that the two proteins can co-partition at concentrations as low as 3 μM each (**Fig. S14e, f**). Furthermore, tau and pHP1α can co-partition at these concentrations at physiological salt with crowding agent added (**Fig. S14f**). We did not pursue LLPS experiments with tau, pHP1α, and 12mer arrays at physiological salt in the presence of PEG, due to the enhanced precipitation of the 12mer arrays under these conditions (**Fig. S4a**).

We next investigated the effect of phosphorylation on tau’s ability to phase separate with pHP1α. PKA and GSK phosphorylation had a minimal effect on co-phase separation with pHP1α and did not interfere with co-localization of 12mer arrays and pHP1α (**Fig. 6d, Fig. S16a-c**). Interestingly, in the case of hyperphosphorylated tau, phase separation was completely abrogated with pHP1α, both in the presence and absence of arrays (**Fig. 6d, Fig. S16a-c**). This suggests that tau’s affinity for pHP1α can be regulated by phosphorylation and that pathological hyperphosphorylation may be disruptive to heterochromatin environments. In further support of this, hyperphosphorylated tau could also dissolve pre-made pHP1α-12mer array droplets (**Fig. S16d**).

## Discussion

While tau is canonically known as a microtubule binding protein, an accumulating body of literature has suggested that it may play additional roles in chromatin stability and heterochromatin organization.^20,31,33^ How tau might carry out these potential roles in the nucleus is currently unknown. By using *in vitro* biophysical experiments, here we show that tau can directly impact the material properties of reconstituted heterochromatin environments through a process that involves chromatin binding, compaction and/or LLPS. This process can be modulated by physiologically relevant concentrations of mono- and divalent salts, post-translational modifications, and other heterochromatin components such as pHP1α. We also demonstrate that tau can protect linker DNA on 12mer arrays but does not cause substantial changes in nucleosome structure and dynamics. These observations suggest that tau’s direct interactions with chromatin may modulate chromatin accessibility, which in turn may contribute to a DNA protective function and/or heterochromatin organization role in the nucleus.

While tau has a well-documented ability to undergo LLPS,^1,4–6,8,9,12^ here we show for the first time that it can also form condensates with DNA, mononucleosomes, and 12mer arrays. This is not surprising as both tau and nucleosomes contain distinct patterns of positive and negative charge that can promote electrostatic multivalent interactions that drive phase separation. The primary interaction driving LLPS, however, appears to be tau’s high affinity for DNA encoded in its PRD and MTBD regions. Deletion of these regions reduces or abolishes LLPS, with the removal of the more positively charged PRD region having a more pronounced effect. Strong interactions with linker and nucleosomal DNA would also explain the relatively low mobility of tau in tau-12mer array droplets. The presence of positively charged histones, however, improves the LLPS propensity of tau, suggesting that other regions of the sequence may also contribute through weaker electrostatic interactions. These interactions may help explain the buffering capacity of tau-12mer array droplets, which can accommodate low levels of phosphorylation throughout the tau sequence.

Our experiments suggest that the LLPS propensity of tau and chromatin is salt dependent. Unlike other electrostatically driven LLPS systems where high salt conditions abolish interactions,^4,47,79^ the effect of salt on tau-12mer array samples is complex. Physiological concentrations of monovalent and divalent salts typically lead to a compact chromatin state,^61^ which we also observe both in the presence of NaCl and MgCl2. Importantly, tau remains associated with chromatin under these conditions even though the LLPS process is no longer observed. While salt can screen the weaker electrostatic interactions that help drive LLPS, tau can remain associated to 12mer arrays through its strong DNA binding propensity. These observations suggest that tau has the ability to interact with chromatin under physiological conditions, whether this interaction is LLPS-based or not.

From a chromatin organization point of view, it is important to understand the consequences of these interactions on chromatin structure and dynamics. Taken together, our experiments suggest that tau may lead to a compact chromatin state mediated by interactions with linker and nucleosomal DNA. Our TEM experiments, for example, show compact oligomeric states in the presence of tau alone. Our MNase digestion assays confirm tau’s ability to reduce the accessibility of linker DNA in the 12mer array context. While we could only perform these experiments under low salt conditions, they are consistent with MNase digestion assays conducted in cells that show similarly reduced chromatin accessibility upon inducible tau expression.^62^ The study further determined that tau-expressing cells were more resistant to heterochromatin disruption induced by histone deacetylase inhibitors and suggested that tau compaction could prevent chromatin remodeling and stabilize heterochromatin.^62^ While our work suggests that tau can do this directly, we cannot preclude the involvement of other chromatin effectors. For instance, tau has also been found associated with Tip5, a major component of the repressive nucleolar remodeling complex,^20^ which suggests tau may interact with and impact chromatin through multiple pathways.

Interestingly, our MAS NMR experiments suggest that tau does not substantially perturb the nucleosome core structure and dynamics. While we detect small structural perturbations at a few contact points between histones and DNA, these most likely report on tau’s interactions with nucleosomal DNA and/or a distinct compact state in the presence of tau. Notably, we do not see changes in histone H4 tail dynamics, which implies that it remains accessible and available for post-translational modifications. Thus, tau may be able to function independently from other modes of nucleosome regulation that rely on post-translational modification of the histone tails.

The biophysical properties of tau in heterochromatin environments would not only be influenced by interactions with chromatin but also by the presence of other heterochromatin components including various proteins and regulatory RNA. Here, we investigated the influence of the heterochromatin component pHP1α whose ability to undergo LLPS has been hypothesized to be a driving force for heterochromatin formation and regulation.^47^ We found that tau and pHP1α co-partition into liquid droplets. Our previous work has shown that pHP1α LLPS can be enhanced by positively charged peptides through specific interactions with pHP1α’s PXVXL-binding interface and/or through non-specific electrostatic interactions.^79^ Since tau does not have a PXVXL motif, its mode of interaction most likely involves contacts between its positively charged regions and the negatively charged phosphorylated N-terminal extension on pHP1α. In support of this explanation, wild-type HP1α does not undergo LLPS with equivalent concentrations of tau. Importantly, however, HP1α can sequester tau into droplets that contain 12mer arrays. We note that our 12mer arrays do not contain H3K9me3, which binds to the chromodomain of pHP1α.^82,83^ Therefore, we expect that in our case, the main interactions between pHP1α and chromatin would be through the positively charged hinge, which can interact with DNA, and the negatively charged N-terminal extension which may contact the positively charged histone tails.^79,80^ Remarkably, however, the colocalization of tau-pHP1α-array samples persists in the presence of up to 150 mM NaCl, even if LLPS is not observed under these conditions.

While phosphorylation of tau can enhance its ability to undergo LLPS in the presence of molecular crowders such as PEG,^6,11^ its influence on tau LLPS in the context of chromatin is complex. Minimal phosphorylation (2-3 sites on average) does not appear to substantially affect LLPS with 12mer arrays or pHP1α, while hyperphosphorylation by a combination of the PKA and GSK-3β kinases has a large effect. While tau-array, tau-pHP1α and tau-pHP1α-array droplets may have some buffering capacity that can accommodate a few phosphorylation modifications on tau, the dramatic reversal in charge upon hyperphosphorylation would abolish key electrostatic interactions that drive LLPS. Surprisingly, however, PKA-GSK-phosphorylated tau still retains relatively high affinity for interactions with DNA and mononucleosomes (as compared to wild-type tau). Therefore, the mechanism of LLPS abolishment most likely stems from the disruption of key tau-tau and tau-pHP1α interactions. For example, previous work has shown that negatively charged peptides can completely abolish LLPS of pHP1α and LLPS of HP1α and DNA.^47,79^ While the effects of hyperphosphorylated tau in the nucleus are likely buffered by other components, our data indicate that the buildup of phosphorylation modifications on tau has the potential to disrupt the material properties of heterochromatin environments. Our observations are consistent with previous reports that pathological forms of tau lead to heterochromatin relaxation and loss of chromocenters.^33^ Some groups have hypothesized that chromatin relaxation results from the loss of tau’s ability to protect against DNA damage,^33^ while others have speculated that tau is a chromatin effector.^62^ We propose a hypothesis that this behavior may also stem from a disrupted ability to partition into condensed heterochromatin environments.

In summary, our data support a model where tau can interact with and compact chromatin while also participating in and modulating the LLPS behavior of other chromatin components (**Fig. 7**). Minimal phosphorylation does not significantly impact this behavior while hyperphosphorylation can lead to disruption. Our model is also consistent with observations implying that minimal phosphorylation of tau retains a functional nuclear role,^17,28,72^ while disease-associated hyperphosphorylation may lead to loss of heterochromatin and exit of tau from the nucleus.^41,42^ Such behavior may contribute to or act in parallel to the formation of cytoplasmic NFTs, a well-known hallmark of Alzheimer’s disease. Thus, our study sheds light on how the biophysical properties of tau might influence its function in health and disease, with mechanisms potentially distinct from those proposed by the amyloid hypothesis. In the future, it will be important to investigate the role of other heterochromatin components, including various regulatory RNAs, as RNA has a well-documented ability to modulate tau LLPS.^5^ We are also looking forward to further cellular studies that investigate tau’s properties and interactions in the nuclear environment and directly probe their effects on chromatin stability, organization, and gene regulation.

**Figure 7.**
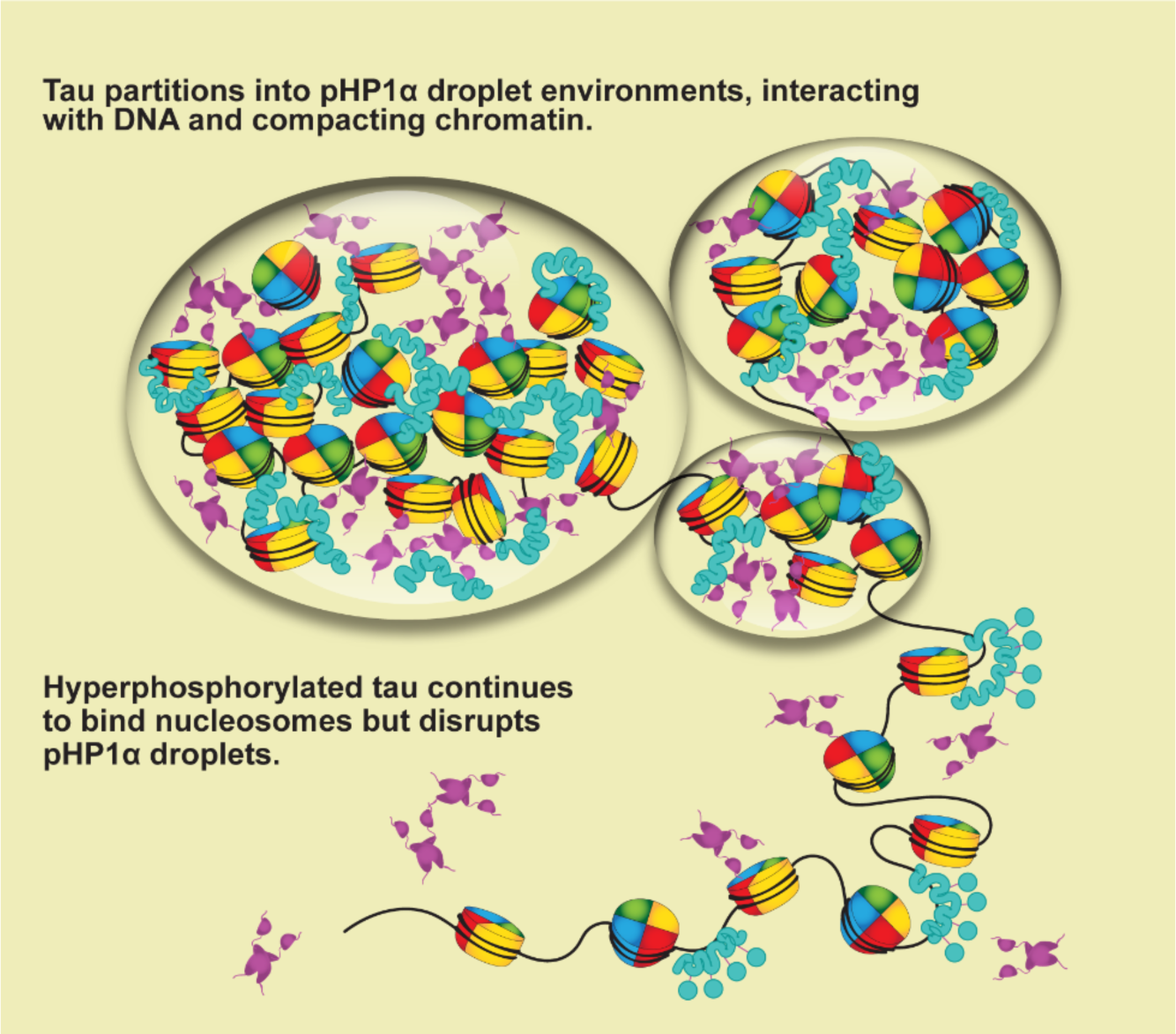
Biophysical model of phosphorylation-dependent phase separation of tau in heterochromatin environments. Minimally or non-phosphorylated tau partitions into pHP1α droplets, interacting with nucleosomal and linker DNA and inducing chromatin compaction. Meanwhile, hyperphosphorylated tau remains chromatin-bound, but cannot phase separate with pHP1α.

## Methods

### Tau expression and purification

Tau was prepared as described previously with some modifications.^84^ Briefly, tau was expressed in BL21 (DE3) Rosetta *Escherichia coli* cells in a pET-2B-T vector. Cells were grown in Luria Bertani media with ampicillin and chloramphenicol at 37°C, induced at an OD600 of 0.6-0.8 using 0.5 mM isopropyl-β-D-thiogalactoside (IPTG), and harvested after 3 hours. The bacterial pellet was resuspended in lysis buffer (20 mM 2-(N-morpholino)ethanesulfonic acid (MES), 500 mM NaCl, 1 mM MgCl2, 1 mM ethylenediaminetetraacetic acid (EDTA), 5 mM dithiothreitol (DTT), pH 6.8) supplemented with Roche EDTA-free Protease Inhibitor Cocktail tablets. Cells were lysed by sonication, boiled for 20 min at 95°C, and the soluble fraction was collected by centrifugation. The supernatant was then dialyzed into a low salt version of the previous buffer (50 mM NaCl), loaded onto a HiTrap SP HP cation exchange column (5 mL) (Cytiva), and tau was eluted using a linear gradient from 0.1 mM to 1M NaCl over 1 hr. Tau truncation products were removed using a Superdex S75 HiLoad column (Cytiva) in 1X phosphate-buffered saline (PBS). Pure fractions were then dialyzed into 20 mM HEPES, 50 mM NaCl, pH 7.2, 0.5 mM tris(2-carboxyethyl) phosphine (TCEP), concentrated to ∼100 μM or higher, aliquoted, and stored at ™80°C until use. The isolated protein was characterized by sodium dodecyl sulfate–polyacrylamide gel electrophoresis (SDS-PAGE), UV-absorbance, reverse-phase high-performance liquid chromatography (RP-HPLC), and mass spectrometry (MS). Tau protein concentration was obtained by using an extinction coefficient εM = 7575 M^−1^cm^−1^ at 280 nm.

### Histone expression and purification

Histones were prepared as previously described.^85^ Briefly, H2A, H2B, H3, and H4 were separately expressed in BL21(DE3) Rosetta *Escherichia coli* cells in Luria Bertani broth at 37°C. For NMR experiments, H4 was expressed in M9 media supplemented with ^15^N ammonium chloride and ^13^C6-glucose. Cells were grown to OD600 of 0.6, induced with 0.5 mM IPTG, and harvested by centrifugation after 2 hours. The bacterial pellet was resuspended in lysis buffer (20 mM Tris-HCl, 200 mM NaCl, 1 mM EDTA, 1 mM DTT, pH 7.6) supplemented with Roche EDTA-free Protease Inhibitor Cocktail, lysed by sonication, and the insoluble pellet was collected for inclusion body extraction. The pellet was resuspended and washed twice with lysis buffer containing 0.1% v/v of Triton-X, washed once with lysis buffer without detergent, and then the pellet was solubilized in 6 M guanidine hydrochloride, 20 mM Tris, 1 mM EDTA, 100 mM NaCl (pH 7.5 at 4°C) for 2 hours at 4°C. The mixture was centrifuged, and then the supernatant was dialyzed against buffer containing 7 M urea, 10 mM Tris, 1 mM EDTA, 100 mM NaCl, 1 mM DTT, pH 7.5 at 4°C, loaded onto a HiTrap SP HP cation exchange column (5 mL) (Cytiva), and eluted using a linear gradient of 0.1 to 1M NaCl. Pure fractions were collected and loaded onto a Waters Xbridge BEH C18 prep-size column and purified by RP-HPLC. Fractions were collected using a gradient of 30-70% acetonitrile in H2O with 0.1% TFA. The isolated protein was characterized by SDS-PAGE, UV-absorbance, RP-HPLC, and MS. Protein concentrations were obtained by using extinction coefficients εM = 4470, 7450, 4470, and 5960 M^−1^cm^−1^ at 280 nm for H2A, H2B, H3, and H4, respectively. Histones were lyophilized and stored at ™80°C until refolding into octamers.

### DNA preparation

12mer array DNA (2.1kbp), which is composed of 12 repeats of the 601 Widom sequence^86^ separated by 30 bp linkers, was prepared as described previously.^55^ Briefly, DH5α cells were transformed with a plasmid containing the 12mer DNA sequence and grown in Terrific Broth for 18 hours at 37°C. The cells were harvested and then resuspended in buffer containing 25 mM Tris-HCl, 50 mM glucose, 10 mM EDTA, pH 8.0. The cells were then lysed with buffer containing 0.2 M NaOH, 1% sodium dodecyl sulfate, and then neutralized with 4 M potassium acetate and 2 N acetic acid. Plasmid DNA was precipitated using isopropanol precipitation and the pellet was then suspended in TE 10/50 buffer (10 mM Tris-HCl, 50 mM EDTA, pH 8.0), and incubated with 20 mg/mL RNase A at 37°C for 12 hours. Proteins were extracted from the mixture using phenol chloroform, and RNA was removed by PEG precipitation. The 12mer DNA sequence was excised from the plasmid using Eco-RV digestion (New England BioLabs), and the excess plasmid was removed via PEG purification. The purified 12mer DNA was concentrated by ethanol precipitation and stored in TE 10/0.1 buffer (10 mM Tris-HCl, 0.1 mM EDTA, pH 8.0), and frozen at ™20°C. 177 bp DNA was prepared by digestion of the 12mer DNA plasmid using ScaI and then purified and stored similarly. The same protocol was used to prepare the 155 bp nucleosome-binding MMTV “buffer DNA”, which is used to sequester excess octamer during 12mer array assembly. 147 bp DNA was prepared using a large-scale PCR amplification starting with the 177 bp DNA as template. The 147 bp DNA was amplified using the following primers: 5’-CTGGAGAATCCCGGTGC-3’ and 5’- GTGTCAGATATATACATCCTGT-3’ and the PCR product was purified using a NucleoSpin Gel and PCR Clean-up Maxi kit (Takara Bio). The sequences of the 147 bp and 177 bp DNA are as follows:

147 bp DNA

CTGGAGAATCCCGGTGCCGAGGCCGCTCAATTGGTCGTAGACAGCTCTAGCACCGCTTAA ACGCACGTACGCGCTGTCCCCCGCGTTTTAACCGCCAAGGGGATTACTCCCTAGTCTCCAGGCACGTGTCAGATATATACATCCTGT

177 bp DNA (<u>with the “601” sequence underlined</u>)

ACTACGCGGCCGCC<u>CTGGAGAATCCCGGTGCCGAGGCCGCTCAATTGGTCGTAGACAGC TCTAGCACCGCTTAAACGCACGTACGCGCTGTCCCCCGCGTTTTAACCGCCAAGGGGATT ACTCCCTAGTCTCCAGGCACGTGTCAGATATATACATCCTG</u>TGCATGTAAGATCCAGT

### Assembly of histone octamers, mononucleosomes, and 12mer arrays

Histone octamers and nucleosomes were assembled as previously described.^55,85^ Lyophilized histones were dissolved at 2 mg/mL in 6 M guanidine hydrochloride, 20 mM Tris-HCl, 5 mM dithiothreitol, pH 7.6 at 4°C at a ratio of 1.0:1.0:0.9:0.9 of H2A:H2B:H3:H4. The octamers were refolded through dialysis into buffer containing 2 M NaCl, 10 mM Tris, 1 mM EDTA, 1 mM dithiothreitol, pH 7.6, and the octamers were purified from histone tetramers and dimers using a Superdex 200 10/30 column (Cytiva). The pure octamers were combined, concentrated, and stored in 50% glycerol at ™20°C. 12mer arrays were reconstituted by combining 1 equivalents 12mer DNA, 0.3 equivalent MMTV DNA,^85,87^ and ∼1.6 equivalent octamers in 2M TEK buffer (10 mM Tris pH 7.5, 0.1 mM EDTA, 2M KCl). The equivalent of octamers can vary and is always optimized by ratio test. The combined DNA and octamer were then subjected to a salt gradient dialysis into 10 mM TEK buffer (10 mM Tris pH 7.5, 0.1 mM EDTA, 10 mM KCl) for ∼24 hours at 4°C. The 12mer arrays were precipitated by addition of 2 mM MgCl2 to purify from MMTV mononucleosomes and excess MMTV DNA and dialyzed into 20 mM HEPES pH 7.2, 10 mM NaCl, 0.5 mM TCEP. The purity of 12mer arrays was confirmed using APAGE (2% acrylamide, 1% Biorad agarose) gels stained with SyBr Gold (Life Technologies). Concentrations were determined using molar extinction coefficient εA260 = 2822957 M^−1^cm^−1^.

### pHP1α expression and purification

pHP1α was prepared as described previously.^67^ BL21(DE3)-Rosetta *Escherichia coli* cells were co-transformed with the His-tagged HP1α and CK2 plasmids to make in-cell phosphorylated HP1α. Cells were grown in Luria Bertani media with ampicillin and streptomycin at 37 °C, induced at an OD600 of 0.6 using 0.5 mM IPTG, transferred to 18 °C, and harvested after 16-24 hours. The bacterial pellet was resuspended in lysis buffer (1x PBS, pH 7.4, 300 mM NaCl, 10% glycerol, 7.5 mM imidazole) supplemented with Roche protease inhibitor tablets and lysed by sonication. The lysate supernatant was then incubated with Ni-NTA resin (5 mL resin per 1 L culture) with rotation at 4 °C for one hour and the lysate-resin was loaded onto a Bio-Rad Econo-Column and washed with wash buffer (1x PBS, pH 7.4, 300 mM NaCl, 10% glycerol, 7.5 mM imidazole), and then eluted with wash buffer containing 400 mM imidazole. The eluate was incubated with TEV-protease during dialysis into imidazole-free buffer overnight at 4 °C. The solution was then adjusted to 6 M guanidine hydrochloride and purified using a Waters XBridge BEH C18 prep-size RP-HPLC column using a gradient of 10-60% acetonitrile in H2O with 0.1% TFA. Pure fractions (assessed by SDS-PAGE) were lyophilized and stored at ™80°C. Protein concentration was obtained by using an extinction coefficient εM = 29160 M^−1^cm^−1^ at 280 nm. HP1α was refolded by resuspending the lyophilized protein in resuspension buffer (20 mM HEPES pH 7.2, 6 M guanidine hydrochloride, 20 mM NaCl, 2 mM DTT) at 1 mg/mL, dialyzing into 20 mM HEPES, 150 mM NaCl, 1M guanidine hydrochloride, at pH 7.2 for 2-3 hours, then dialyzing against guanidinium-free buffer with 300 mM NaCl overnight at 4 °C. 300 mM NaCl is used to prevent phase separation of pHP1α during the concentration step. Unmodified HP1α was expressed and purified by an identical procedure, except that the CK2 plasmid was omitted in the initial transformation step.

### Fluorophore conjugation

Tau was labelled on its 2 native cysteines using Cy5 maleimide. To label, excess Cy5-maleimide was added to 100 μM tau overnight at room temperature with rotation and was then quenched with excess β-mercaptoethanol. Labeled tau was then purified by RP-HPLC from excess Cy5 using a semi-preparative C18 RP-HPLC column using a 20–70% acetonitrile gradient with 0.1% TFA. Single and double-labelled fractions were lyophilized separately. Samples were resuspended in 20 mM HEPES, 50 mM NaCl, 6 M guanidine hydrochloride, 0.5 mM TCEP, pH 7.2, and subjected to dialysis steps into the same buffer without guanidine hydrochloride. Doubly labelled tau was used for microscopy experiments, and conjugation was confirmed by ESI-TOF-MS, SDS-PAGE, and fluorescence was visualized using a Typhoon FLA 9500 (Cytiva) imager.

Fluorescein-labelled 12mer arrays were prepared through conjugation of fluorescein to H2A using maleimide chemistry, prior to incorporation into octamers.^85^ The conjugation was performed at position 110 where the native Asn residue was mutated to Cys. H2A N110C was purified by cation exchange and RP-HPLC as described above, and 9 mg of lyophilized protein was dissolved in 3 mL of labeling buffer (20 mM Tris, pH 7.8, 6 M guanidine hydrochloride, 0.5 mM TCEP) to a final concentration of ∼150 μM. The solution was incubated with TCEP for 1 hour with stirring, followed by the addition of 1.5 mg fluorescein-5-maleimide (dissolved in 50 μL N,N-dimethylformamide). The reaction was allowed to proceed for 1 hour in the dark, with stirring, and was quenched with 1 mM 2- mercaptoethanol. Fluorescently labeled H2A was purified using a semi-preparative C18 RP-HPLC column using a 30–70% HPLC acetonitrile gradient with 0.1% TFA. Conjugation was confirmed by ESI-TOF-MS, SDS-PAGE and fluorescence was visualized using a Typhoon FLA 9500 (Cytiva) imager.

HP1α was labelled with Cy3 as previously described.^47,67^ To avoid perturbing the native cysteines in its folded domains, a GSKCK tag was added to the C-terminus of HP1α and a S133C mutation was introduced. This construct was co-expressed with the CK2 kinase and purified by the same protocol as unmodified pHP1α (see above). To label, excess Cy3-maleimide was added to 100 μM pHP1α, and then quenched with excess β-mercaptoethanol after a 10 second reaction, so that only the most solvent-exposed cysteine could be labelled. The labeled protein was purified by RP-HPLC from excess Cy3 using a gradient of acetonitrile with 0.1% TFA. Pure fractions were lyophilized and refolded similarly to unmodified pHP1α.

### In vitro phosphorylation of tau

Tau was phosphorylated in *vitro* according to previous protocols with some modifications.^12,88^ 50 µL of 100 µM tau was incubated with 2 µg cAMP-dependent Protein Kinase (PKA, New England BioLabs P6000S) and/or 0.4 µg GSK3β (Sigma-Aldrich G4296) in a buffer containing 20 mM HEPES (pH 7.2), 50 mM NaCl, 0.5 mM TCEP, 4 mM adenosine-5′- triphosphate (ATP; Gold Bio), 1 mM phenylmethylsulfonyl fluoride (PMSF), 10 mM MgCl2, at 30°C for 16 hours. Kinases were then heat inactivated for 20 minutes at 65 °C, followed by centrifugation to remove precipitated kinase. Phosphorylated protein was then buffer exchanged into 20 mM HEPES, 50 mM NaCl, pH 7.2, 0.5 mM TCEP using a 30 kDa cutoff centrifugal concentrator (Sartorius) to remove MgCl2 and ATP. Phosphorylation was verified via SDS-PAGE and ESI-TOF-MS, while specific sites were identified using LC-MS/MS.

### Liquid chromatography-tandem mass spectrometry (LC-MS/MS)

For phosphorylated samples, detailed post-translational modification mapping was obtained from LC-MS/MS experiments performed according to the protocol described in Ref.^89^. Briefly, the proteins were analyzed on an SDS-PAGE gel and stained with Coomassie. The relevant protein band was extracted from the gel, reduced with DTT, alkylated with iodacetamide, and trypsin-digested in-gel with ammonium bicarbonate. The trypsin-digested peptides were then analyzed by ultra-high-pressure liquid chromatography (UPLC) coupled with tandem mass spectrometry (LC-MS/MS) using nano-spray ionization, on an Orbitrap Fusion Lumos hybrid mass spectrometer (Thermo) coupled with a nanoscale reversed-phase UPLC (Thermo Dionex UltiMate 3000 RSLC nano System) using a 25 cm, 75-micron ID glass capillary packed with 1.7- µm C18 (130) BEHTM beads (Waters). Peptides were eluted using a 60-minute linear gradient (5–80%) of Buffer A (98% H2O, 2% acetronitrile, 0.1% formic acid) to Buffer B (100% acetonitrile, 0.1% formic acid) at a flow rate of 0.375 mL/min. In the survey scan, MS1 spectra were measured in an m/z range of 400 to 1500 with a resolution of 120000. The spray voltage was set to 2200 V, ion transfer tube temperature of 275 °C, and a maximum injection time of 50 ms. An initial survey scan was followed by data dependent scans of the most abundant ions (charge from +2 to +5), while selecting ions with minimum intensities of 50000 and fragmented with a higher-energy collision dissociation at 30% collision energy. Fragmented masses were analyzed in the mass analyzer with an ion trap scan rate of turbo, first mass m/z of 100, automatic gain control (AGC) target of 5000, maximum injection time of 35 ms, and exclusion time set to 5 seconds.

Three to four independent replicates were obtained for each condition, and phosphorylation sites observed across different replicates and different conditions are tabulated in **Table S3**. All MS/MS data were processed in the MaxQuant software (version 2.0.3.0)^90^ and searched against the *Homo sapiens* Uniprot protein sequence database and common contaminants. Phosphorylation of serine, threonine, and tyrosine residues was searched as a variable modification, and carbamidomethylation of cysteine residues was set as a fixed modification. Peptide mass tolerance was set to 20 ppm, fragment mass tolerance was set to 0.5 Da, and the false discovery rate was set to 1%. The digestion enzyme was set to trypsin/P and a maximum of two missed cleavages were allowed.

### Confocal microscopy

LLPS was initiated by addition of other components (1 μM DNA, 1 μM nucleosome, 80 nM 12mer arrays (equivalent to 1 μM ‘601’ nucleosome sites), 50 μM pHP1α, 50 μM HP1α, or 10% PEG) to tau (1.6 μM-50 μM) in LLPS buffer (20 mM HEPES, 25 mM NaCl, 0.5 mM TCEP, pH 7.2) at room temperature. Fluorophore-labeled protein was mixed with unlabeled protein in a molar ratio of 1:20, and 0.8 μM of YOYO-1 was added to 1 μM eq. of mononucleosomes, 601 sites in 12mer arrays, and DNA. Samples (5 µL) were placed in a µ-Slide 18-well glass bottom multi-well plate (Ibidi) for imaging and were allowed to settle for 5-10 minutes. Images were acquired with a Leica SP8 microscope with a 40X oil immersion objective.

### Fluorescence recovery after photobleaching

**(FRAP)** Droplets were prepared as described above and imaged using a Leica SP8 confocal microscope. In each experiment, 3 to 4 μm regions of interest were bleached at 100% laser power in the center of each droplet. Two pre-bleach frames were acquired, and 100 post-bleaching frames were acquired (1.295 s/frame), corresponding to 2 minutes of recovery. Data were analyzed using Fiji/ImageJ. The FRAP curves were normalized to the maximal prebleach and minimal postbleach intensities, and an unbleached reference was used to correct for photobleaching due to image acquisition. The error bars represent standard deviation from data collected with 20 or more droplets from at least two independent samples and the curve represents the best fit to a one phase decay model.

### Electrophoresis mobility shift assay

**(EMSA)** Reconstituted mononucleosomes (147 bp or 177 bp mononucleosomes), 22 bp DNA, and 177 bp DNA were incubated to a final concentration of 20 nM with various concentrations of tau for 20 minutes at room temperature. Reaction conditions were 20 mM HEPES buffer, 10 mM NaCl, pH 7.2, 0.5 mM TCEP, 0.1% Tween-20 with 5% sucrose. Samples were analyzed on a 5% TBE gel at 90V for 90 min for 177 bp DNA and mononucleosomes, and 60 minutes for 22 bp DNA. DNA was visualized through staining with SYBR Gold or by visualizing fluorescein-conjugated 22 bp DNA using a Typhoon FLA 9500 (Cytiva) imager. The fraction bound was calculated from the amount of remaining free mononucleosome or DNA band in each individual lane. Error bars represent the standard deviation from three independent EMSA experiments. Curves were fitted to a one site specific binding with Hill Slope equation in GraphPad Prism 9.5.

### Fluorescence anisotropy

**(FA)** Fluorescein conjugated 22 bp DNA was prepared by annealing reverse complementary oligonucleotides. Briefly, the forward fluorescein-conjugated oligonucleotide (5’-6-FAM) 5’- ATTTAGAAATGTCCACTGTAGG-3’ (Integrated DNA Technologies) and its non-conjugated reverse complement were resuspended in 20 mM HEPES, 10 mM NaCl, 0.5 mM TCEP, pH 7.2 at 100 μM and then mixed at a 1:1 ratio, denatured for 5 min at 95 °C in a water bath, and then slowly cooled and annealed by turning off the heat. The annealed oligonucleotide was used directly for EMSA and fluorescence polarization experiments. Various concentrations of tau were mixed with 10 nM of fluorescein-labelled DNA in 20 mM HEPES,10 mM NaCl, 0.5 mM TCEP, 0.01% Nonidet P40 substitute, pH 7.2, incubated at room temperature for 10 min. Fluorescence polarization was detected using a Tecan Infinite M1000 PRO fluorescence plate reader. Binding data were fit to a one site specific binding with Hill Slope equation in GraphPad Prism 9.5

### Turbidity assays

Turbidity of protein samples was estimated from the optical density at 600 nm using a NanoDrop spectrophotometer (ThermoFisher). Protein solutions were pre-incubated at room temperature, and proteins were thoroughly mixed at a 10 μL final volume and immediately measured for turbidity. All turbidity measurements were conducted in 20 mM HEPES buffer, 25 mM NaCl, 0.5 mM TCEP, pH 7.2. Three technical replicates were obtained for each sample, as well as three independent replicates per condition.

### Chromatin oligomerization assay

For data shown in Fig. 3b, 30 nM 12mer arrays (at an A260 of ∼1) were mixed with increasing amounts of tau (∼0-12.5 μM) at 25 or 150 mM NaCl, incubated at room temperature for 10 minutes and then centrifuged at 13,000xg for 10 minutes. The A260 of the supernatant was then measured and normalized to the value of fully soluble 12mer arrays. All assays were conducted in 20 mM HEPES buffer, 0.5 mM TCEP, pH 7.2, with 25 or 150 mM NaCl. Three technical replicates were obtained for each sample, as well as three independent replicates per condition. 10 mM MgCl2 (no NaCl in the buffer) or 10% PEG (and 150 mM NaCl) were used as controls for the experiments shown in supplementary Fig. S4a.

### Transmission electron microscopy

**(TEM)** 12mer arrays (at 20 ng/uL, or 11.3 nM), with or without various additives (0.136 μM and 1.36 μM tau, and 0.5 and 1 mM MgCl2), were incubated for 20 minutes in 20 mM HEPES buffer, 10 mM NaCl, 0.5 mM TCEP, pH 7.2. Then, glutaraldehyde was added to a final concentration of 0.1%, and the sample was cross-linked overnight on ice at 4°C. Glutaraldehyde was then removed by dialysis against 20 mM HEPES, 10 mM NaCl, 0.5 mM TCEP, pH 7.2 for 2 hours. Each solution was then directly loaded onto freshly glow-discharged carbon-coated electron microscopy (EM) grids (Electron Microscopy Science) for 10 min, washed with 50 mM NaCl, stained with 2% (w/v) uranyl acetate for 1.5 minutes, and then dried. Images were then acquired using a JEOL 1400 Plus microscope with a bottom-mount Gatan OneView (4k x 4k) camera or a FEI Tecnai Spirit G2 BioTWIN microscope with a bottom mount Eagle (4k x 4k) camera.

### Micrococcal nuclease (MNase) protection assay

12mer arrays (40 ng) were digested with 1 U of MNase (New England Biolabs) in a 10 μL reaction for varying digestion times (2.5 to 10 minutes) in buffer containing 20 mM HEPES, 10 mM NaCl, 0.5 mM TCEP, pH 7.2, supplemented with 1 mM CaCl2. For reactions containing more components (i.e., tau or MgCl2), samples were incubated for 20 minutes at room temperature prior to addition of MNase. After digestion, reactions were quenched with the addition of final concentrations of 15% Proteinase K (New England Biolabs) (∼1.2 U) and 62 mM EDTA and incubated at 50°C for 30 min. The DNA was analyzed on a 5% TBE gel, stained by SYBR Gold, and imaged using a Typhoon FLA 9500 (Cytiva) imager.

### Magic angle spinning nuclear magnetic resonance (MAS NMR) spectroscopy

Large-scale 12mer array assembly was performed as described previously^91^ in a 20 mL total volume, at a concentration of 2.5 μM 601 sites, in 10 mM TEK buffer (10 mM Tris pH 7.5, 0.1 mM EDTA, 10 mM KCl) with ^15^N,^13^C-labeled H4. 12mer arrays were purified by sequential MgCl2 precipitations, and pure pelleted 12mer arrays were resuspended and pooled together. To prepare the control sample for NMR, 10 mg of purified 12mer arrays were precipitated with 6 mM MgCl2, centrifuged, and packed into a rotor. For the tau-array sample, 6 mM MgCl2 and 20 mg of tau (in solution, ∼700 μM) were added to ∼ 6 mg of 12mer arrays, and the sample was similarly centrifuged and packed into a rotor. While 12mer arrays alone were largely solid, tau-12mer arrays were gel-like and could be packed into a rotor using a plunger pipette.

NMR data were collected using a 750 MHz NMR spectrometer, equipped with an AVANCE II Bruker console and a triple resonance (^1^H, ^13^C, ^15^N) 3.2 mm magic angle spinning (MAS) E- free Bruker probe. All experiments were performed using 3.2 mm thin-wall Bruker rotors with an MAS spinning frequency of 11.1 kHz. Spectra were referenced to adamantane (^13^C = 40.49 ppm), and the magic angle was calibrated using KBr.^92^ 2D ^13^C-^13^C CP DARR experiments^93^ were recorded at 275 K with the following acquisition settings: 20 ms mixing, 1024 points and 9 ms acquisition time in the direct dimension and 256 points and 3.2 ms acquisition in the indirect dimension. 144 scans were collected for the tau-array spectrum and 64 scans were collected for the control, due to the difference in sample amount. 2D ^13^H-^13^C INEPT experiments^94^ were recorded at 295 K with the following acquisition settings: 1024 points and 9 ms acquisition time in the direct dimension and 500 points and 3.3 ms acquisition in the indirect dimension. 64 scans were obtained for both control and tau-arrays. NMR spectra were processed with TopSpin 4.0.5 and analyzed using NMRFAM-SPARKY^95^ and CCPNMR 3.0.2.^96^ H4 chemical shift assignments were transferred from published work.^63^ Nucleosome images were rendered using Chimera.^97^

## Author contributions

L.A. and G.T.D. designed the project; L.A. prepared all materials and performed imaging and turbidity experiments, TEM, MS, NMR, and binding studies; N.E. prepared nucleosomes and performed NMR experiments; M.M. performed MNase assays; A.D. performed FA experiments; L.A. and G.T.D. analyzed data and wrote the manuscript with input from all authors; K.C. and G.T.D. supervised the project.

## Acknowledgements

This work was supported by NIH grants R21AG079239 to G.T.D., R35GM144121 to K.C., T32GM139795 fellowship to L.A., T32GM008326 fellowship to N.E., and the UC San Diego Dean of Physical Sciences Undergraduate Summer Research fellowship to M.M. In addition, we thank the UC San Diego core facilities used in this study: Biomolecular and Proteomics Mass Spectrometry Facility (S10 OD021724), the Microscopy Core (NINDS P30NS047101), the Electron Microscopy Core, and the Molecular Mass Spectrometry Facility. We are grateful for the technical assistance of J. Santini, M. Erb, T. Merloo, G. Castillon, Y. Su, and M. Ghassemian from the core facilities. We also thank B. Ackermann for sharing various reagents.

## Materials and Correspondence

All inquiries should be addressed to.

## Data Availability

All relevant data are available in the manuscript or supplementary information. Processed NMR spectra in Sparky format will be shared on Zenodo upon publication.

## References

1. Wegmann, S. Liquid-Liquid Phase Separation of Tau Protein in Neurobiology and Pathology. Adv Exp Med Biol 1184, 341–357 (2019).

2. Nelson, P. T. et al. Correlation of Alzheimer disease neuropathologic changes with cognitive status: A review of the literature. J Neuropathol Exp Neurol 71, 362–381 (2012).

3. Martin, L. et al. Tau protein kinases: Involvement in Alzheimer’s disease. Ageing Res Rev 12, 289–309 (2013).

4. Boyko, S., Qi, X., Chen, T. H., Surewicz, K. & Surewicz, W. K. Liquid-liquid phase separation of tau protein: The crucial role of electrostatic interactions. Journal of Biological Chemistry 294, 11054–11059 (2019).

5. Zhang, X. et al. RNA stores tau reversibly in complex coacervates. PLoS Biol 15, 1–28 (2017).

6. Wegmann, S. et al. Tau protein liquid–liquid phase separation can initiate tau aggregation. EMBO J 37, 1–21 (2018).

7. Hernández-Vega, A. et al. Local Nucleation of Microtubule Bundles through Tubulin Concentration into a Condensed Tau Phase. Cell Rep 20, 2304–2312 (2017).

8. Ambadipudi, S., Biernat, J., Riedel, D., Mandelkow, E. & Zweckstetter, M. Liquid-liquid phase separation of the microtubule-binding repeats of the Alzheimer-related protein Tau. Nat Commun 8, 1–13 (2017).

9. Hochmair, J. et al. Molecular crowding and RNA synergize to promote phase separation, microtubule interaction, and seeding of Tau condensates. EMBO J 41, (2022).

10. Najafi, S. et al. Liquid–liquid phase separation of Tau by self and complex coacervation. Protein Science 30, 1393–1407 (2021).

11. Kanaan, N. M., Hamel, C., Grabinski, T. & Combs, B. Liquid-liquid phase separation induces pathogenic tau conformations in vitro. Nat Commun 11, (2020).

12. Savastano, A. et al. Disease-associated tau phosphorylation hinders tubulin assembly within tau condensates. Angewandte Chemie International Edition 1–6 (2020) doi:10.1002/anie.202011157.

13. Lin, Y., Fichou, Y., Zeng, Z., Hu, N. Y. & Han, S. Electrostatically Driven Complex Coacervation and Amyloid Aggregation of Tau Are Independent Processes with Overlapping Conditions. ACS Chem Neurosci 11, 615–627 (2020).

14. Greenwood, J. A. & Johnson, G. V. W. Localization and in Situ Phosphorylation State of Nuclear Tau. Exp Cell Res 220, 332–337 (1995).

15. Loomis, P. A., Howard, T. H., Castleberry, R. P. & Binder, L. I. Identification of nuclear tau isoforms in human neuroblastoma cells. Proc Natl Acad Sci U S A 87, 8422–8426 (1990).

16. Gunawardana, C. G. et al. The human tau interactome: Binding to the ribonucleoproteome, and impaired binding of the proline-to-leucine mutant at position 301 (P301L) to chaperones and the proteasome. Molecular and Cellular Proteomics 14, 3000–3014 (2015).

17. Gil, L. et al. Aging dependent effect of nuclear tau. Brain Res 1677, 129–137 (2017).

18. Thurston, V. C., Pena, P., Pestell, R. & Binder, L. I. Nucleolar localization of the microtubule-associated protein tau in neuroblastomas using sense and anti-sense transfection strategies. Cell Motil Cytoskeleton 38, 100–110 (1997).

19. Sjöberg, M. K., Shestakova, E., Mansuroglu, Z., Maccioni, R. B. & Bonnefoy, E. Tau protein binds to pericentromeric DNA: A putative role for nuclear tau in nucleolar organization. J Cell Sci 119, 2025–2034 (2006).

20. Maina, M. B. et al. The involvement of tau in nucleolar transcription and the stress response. Acta Neuropathol Commun 6, 70 (2018).

21. Brady, R. M., Zinkowski, R. P. & Binder, L. I. Presence of tau in isolated nuclei from human brain. Neurobiol Aging 16, 479–486 (1995).

22. Tracy, T. E. et al. Tau interactome maps synaptic and mitochondrial processes associated with neurodegeneration. Cell 185, 712–728.e14 (2022).

23. Hua, Q. et al. Microtubule associated protein tau binds to double-stranded but not single-stranded DNA. Cellular and Molecular Life Sciences 60, 413–421 (2003).

24. Krylova, S. M. et al. Tau protein binds single-stranded DNA sequence specifically - The proof obtained in vitro with non-equilibrium capillary electrophoresis of equilibrium mixtures. FEBS Lett 579, 1371–1375 (2005).

25. Qi, H. et al. Nuclear magnetic resonance spectroscopy characterization of interaction of Tau with DNA and its regulation by phosphorylation. Biochemistry 54, 1525–1533 (2015).

26. Hua, Q. & He, R.-Q. Effect of Phosphorylation and Aggregation on Tau Binding to DNA. Protein Pept Lett 9, 349–357 (2005).

27. Wei, Y. et al. Binding to the minor groove of the double-strand, Tau protein prevents DNA damage by peroxidation. PLoS One 3, (2008).

28. Ulrich, G. et al. Phosphorylation of nuclear Tau is modulated by distinct cellular pathways. Sci Rep 8, 1–14 (2018).

29. Sultan, A. et al. Nuclear Tau, a key player in neuronal DNA protection. Journal of Biological Chemistry 286, 4566–4575 (2011).

30. Violet, M. et al. A major role for Tau in neuronal DNA and RNA protection in vivo under physiological and hyperthermic conditions. Front Cell Neurosci 8, 1–11 (2014).

31. Mansuroglu, Z. et al. Loss of Tau protein affects the structure, transcription and repair of neuronal pericentromeric heterochromatin. Sci Rep 6, 1–16 (2016).

32. Bou Samra, E., et al. A role for Tau protein in maintaining ribosomal DNA stability and cytidine deaminase-deficient cell survival. Nat Commun 8, (2017).

33. Frost, B., Hemberg, M., Lewis, J. & Feany, M. B. Tau promotes neurodegeneration through global chromatin relaxation. Nat Neurosci 17, 357–366 (2014).

34. Frost, B. Alzheimer’s disease and related tauopathies: disorders of disrupted neuronal identity. Trends Neurosci 46, 797–813 (2023).

35. Sun, W., Samimi, H., Gamez, M., Zare, H. & Frost, B. Pathogenic tau-induced piRNA depletion promotes neuronal death through transposable element dysregulation in neurodegenerative tauopathies. Nat Neurosci 21, 1038–1048 (2018).

36. Guo, C. et al. Tau Activates Transposable Elements in Alzheimer’s Disease. Cell Rep 23, 2874–2880 (2018).

37. Beckmann, A. et al. Moesin is an effector of tau-induced actin overstabilization, cell cycle activation, and neurotoxicity in Alzheimer’s disease. iScience 26, (2023).

38. Fulga, T. A. et al. Abnormal bundling and accumulation of F-actin mediates tau-induced neuronal degeneration in vivo. Nat Cell Biol 9, 139–148 (2007).

39. Frost, B., Bardai, F. H. & Feany, M. B. Lamin Dysfunction Mediates Neurodegeneration in Tauopathies. Current Biology 26, 129–136 (2016).

40. Sohn, C., Ma, J., Ray, W. J. & Frost, B. Pathogenic tau decreases nuclear tension in cultured neurons. Frontiers in Aging 4, (2023).

41. Gil, L. et al. Perinuclear lamin a and nucleoplasmic lamin B2 characterize two types of hippocampal neurons through Alzheimer’s disease progression. Int J Mol Sci 21, 1–19 (2020).

42. Hernández-Ortega, K., Garcia-Esparcia, P., Gil, L., Lucas, J. J. & Ferrer, I. Altered Machinery of Protein Synthesis in Alzheimer’s: From the Nucleolus to the Ribosome. Brain Pathology 26, 593–605 (2016).

43. Lester, E. et al. Tau aggregates are RNA-protein assemblies that mislocalize multiple nuclear speckle components. Neuron 109, 1675–1691.e9 (2021).

44. Benhelli-Mokrani, H. et al. Genome-wide identification of genic and intergenic neuronal DNA regions bound by Tau protein under physiological and stress conditions. Nucleic Acids Res 46, 11405–11422 (2018).

45. Wang, X. S. et al. The Proline-Rich Domain and the Microtubule Binding Domain of Protein Tau Acting as RNA Binding Domains. Protein Pept Lett 13, 679–685 (2006).

46. Hua, Q. & He, R.-Q. Tau could protect DNA double helix structure. Biochim Biophys Acta Proteins Proteom 1645, 205–211 (2003).

47. Larson, A. G. et al. Liquid droplet formation by HP1α suggests a role for phase separation in heterochromatin. Nature 547, 236–240 (2017).

48. Laflamme, G. & Mekhail, K. Biomolecular condensates as arbiters of biochemical reactions inside the nucleus. Commun Biol 3, 773 (2020).

49. Strom, A. R. et al. Phase separation drives heterochromatin domain formation. Nature 547, 241–245 (2017).

50. Sanulli, S. et al. HP1 reshapes nucleosome core to promote phase separation of heterochromatin. Nature 575, 390–394 (2019).

51. Wang, L., Gao, Y., Li, G., Li, H. & Li, P. Histone Modifications Regulate Chromatin Compartmentalization by Contributing to a Phase Separation Mechanism. Mol Cell 76, 646–659.e6 (2019).

52. Erdel, F. et al. Mouse Heterochromatin Adopts Digital Compaction States without Showing Hallmarks of HP1-Driven Liquid-Liquid Phase Separation. Mol Cell 78, 236 (2020).

53. Grewal, S. I. S. The molecular basis of heterochromatin assembly and epigenetic inheritance. Mol Cell 0, (2023).

54. Liu, C. & Götz, J. Profiling murine tau with 0N, 1N and 2N isoform-specific antibodies in brain and peripheral organs reveals distinct subcellular localization, with the 1N isoform being enriched in the nucleus. PLoS One 8, 1–18 (2013).

55. Dyer, P. N. et al. Reconstitution of Nucleosome Core Particles from Recombinant Histones and DNA. Methods Enzymol 375, 23–44 (2003).

56. Ukmar-Godec, T. et al. Lysine/RNA-interactions drive and regulate biomolecular condensation. Nat Commun 10, (2019).

57. Hansen, J. C. Conformational dynamics of the chromatin fiber in solution: Determinants, mechanisms, and functions. Annu Rev Biophys Biomol Struct 31, 361–392 (2002).

58. Hansen, J. C., Maeshima, K. & Hendzel, M. J. The solid and liquid states of chromatin. Epigenetics Chromatin 14, 50 (2021).

59. Hazan, N. P. et al. Nucleosome Core Particle Disassembly and Assembly Kinetics Studied Using Single-Molecule Fluorescence. Biophys J 109, 1676–1685 (2015).

60. Camero, S. et al. Thermodynamics of the interaction between Alzheimer’s disease related Tau protein and DNA. PLoS One 9, (2014).

61. Schwarz, P. M., Felthauser, A., Fletcher, T. M. & Hansen, J. C. Reversible oligonucleosome self-association: Dependence on divalent cations and core histone tail domains. Biochemistry 35, 4009–4015 (1996).

62. Rico, T. et al. Tau Stabilizes Chromatin Compaction. Front Cell Dev Biol 9, (2021).

63. Shi, X. et al. Structure and Dynamics in the Nucleosome Revealed by Solid-State NMR. Angewandte Chemie - International Edition 57, 9734–9738 (2018).

64. Ackermann, B. E. & Debelouchina, G. T. Emerging Contributions of Solid-State NMR Spectroscopy to Chromatin Structural Biology. Front Mol Biosci 8, 1–12 (2021).

65. Baldus, M. & Meier, B. H. Total correlation spectroscopy in the solid state. the use of scalar couplings to determine the through-bond connectivity. J Magn Reson A 121, 65–69 (1996).

66. Berkeley, R. F., Kashefi, M. & Debelouchina, G. T. Real-time observation of structure and dynamics during the liquid-to-solid transition of FUS LC. Biophys J 120, 1276–1287 (2021).

67. Ackermann, B. E. & Debelouchina, G. T. Heterochromatin Protein HP1α Gelation Dynamics Revealed by Solid-State NMR Spectroscopy. Angewandte Chemie - International Edition 58, 6300–6305 (2019).

68. Matlahov, I. & van der Wel, P. C. A. Hidden motions and motion-induced invisibility: Dynamics-based spectral editing in solid-state NMR. Methods 148, 123–135 (2018).

69. Gao, M. et al. Histone H3 and H4 N-terminal tails in nucleosome arrays at cellular concentrations probed by magic angle spinning NMR spectroscopy. J Am Chem Soc 135, 15278–15281 (2013).

70. Davey, C. A., Sargent, D. F., Luger, K., Maeder, A. W. & Richmond, T. J. Solvent mediated interactions in the structure of the nucleosome core particle at 1.9 Å resolution. J Mol Biol 319, 1097–1113 (2002).

71. Drummond, E. et al. Phosphorylated tau interactome in the human Alzheimer’s disease brain. Brain 143, 2803–2817 (2020).

72. Rossi, G. et al. A new function of microtubule-associated protein tau: involvement in chromosome stability. Cell Cycle 4101, (2008).

73. Kopke, E. et al. Microtubule-associated protein tau. Abnormal phosphorylation of a non-paired helical filament pool in Alzheimer disease. Journal of Biological Chemistry 268, 24374–24384 (1993).

74. Lu, Y. et al. Hyperphosphorylation results in tau dysfunction in DNA folding and protection. Journal of Alzheimer’s Disease 37, 551–563 (2013).

75. Wang, J. Z., Grundke-Iqbal, I. & Iqbal, K. Kinases and phosphatases and tau sites involved in Alzheimer neurofibrillary degeneration. European Journal of Neuroscience 25, 59–68 (2007).

76. Mondragón-Rodríguez, S., Perry, G., Luna-Muñoz, J., Acevedo-Aquino, M. C. & Williams, S. Phosphorylation of tau protein at sites Ser396-404 is one of the earliest events in Alzheimer’s disease and Down syndrome. Neuropathol Appl Neurobiol 40, 121–135 (2014).

77. Tepper, K. et al. Oligomer formation of tau protein hyperphosphorylated in cells. Journal of Biological Chemistry 289, 34389–34407 (2014).

78. Larson, A. G. & Narlikar, G. J. The Role of Phase Separation in Heterochromatin Formation, Function, and Regulation. Biochemistry 57, 2540–2548 (2018).

79. Her, C., Phan, T. M., Jovic, N., Kapoor, U. & Ackermann, B. E. Molecular interactions underlying the phase separation of HP1α: Role of phosphorylation, ligand and nucleic acid binding. 1–21 (2022).

80. Keenen, M. M. et al. HP1 proteins compact DNA into mechanically and positionally stable phase separated domains. Elife 10, 1–38 (2021).

81. Nishibuchi, G. et al. N-terminal phosphorylation of HP1α increases its nucleosome-binding specificity. Nucleic Acids Res 42, 12498–12511 (2014).

82. Bannister, A. J. et al. Selective recognition of methylated lysine 9 on histone H3 by the HP1 chromo domain. Nature 410, 120–124 (2001).

83. Lachner, M., O’Carroll, D., Rea, D., Mechtler, K. & Jenuwein, T. Methylation of histone H3 lysine 9 creates a binding site for HP1 proteins. Nature 410, 116–120 (2001).

84. Barghorn, S. & Mandelkow, E. Toward a unified scheme for the aggregation of tau into Alzheimer paired helical filaments. Biochemistry 41, 14885–14896 (2002).

85. Debelouchina, G. T., Gerecht, K. & Muir, T. W. Ubiquitin utilizes an acidic surface patch to alter chromatin structure. Nat Chem Biol 13, 105–110 (2017).

86. Lowary, P. T. & Widom, J. New DNA Sequence Rules for High Affinity Binding to Histone Octamer and Sequence-directed Nucleosome Positioning. J Mol Biol (1998).

87. Flaus, A. & Richmond, T. J. Positioning and Stability of Nucleosomes on MMTV 3 H LTR Sequences. J Mol Biol 275, 427–441 (1998).

88. Zheng-Fischhöfer, Q. et al. Sequential phosphorylation of Tau by glycogen synthase kinase-3β and protein kinase A at Thr212 and Ser214 generates the Alzheimer-specific epitope of antibody AT100 and requires a paired-helical-filament-like conformation. Eur J Biochem 252, 542–552 (1998).

89. Shevchenko, A., Wilm, M., Vorm, O. & Mann, M. Mass spectrometric sequencing of proteins from silver-stained polyacrylamide gels. Anal Chem 68, 850–858 (1996).

90. Tyanova, S., Temu, T. & Cox, J. The MaxQuant computational platform for mass spectrometry – based shotgun proteomics. Nat Protoc 11, 2301–2319 (2016).

91. Elathram, N., Ackermann, B. & Debelouchina, G. DNP-enhanced solid-state NMR spectroscopy of chromatin polymers. J Magn Reson Open (2022).

92. Morcombe, C. R. & Zilm, K. W. Chemical shift referencing in MAS solid state NMR. Journal of Magnetic Resonance 162, 479–486 (2003).

93. Takegoshi, K., Nakamura, S. & Terao, T. 13C–1H dipolar-assisted rotational resonance in magic-angle spinning NMR. Chemical Physical Letters 344, 631–637 (2001).

94. Morris, G. A. & Freeman, R. Enhancement of nuclear magnetic resonance signals by polarization transfer. J Am Chem Soc 101, 760–762 (1979).

95. Lee, W., Tonelli, M. & Markley, J. L. NMRFAM-SPARKY: Enhanced software for biomolecular NMR spectroscopy. Bioinformatics 31, 1325–1327 (2015).

96. Skinner, S. P. et al. CcpNmr AnalysisAssign: a flexible platform for integrated NMR analysis. J Biomol NMR 66, 111–124 (2016).

97. Pettersen, E. F. et al. UCSF Chimera - A visualization system for exploratory research and analysis. J Comput Chem 25, 1605–1612 (2004).

