## Supplementary figures and tables for "Phosphorylation regulates tau’s phase separation behavior and interactions with chromatin"

**Contents**

Figure S1. Analysis of purified 1N4R tau.
Figure S2. Nucleosome and 12mer array constructs.
Figure S3. Behavior of tau under crowding conditions.
Figure S4. Effect of mono- and divalent salt on 12mer arrays and tau-array LLPS.
Figure S5. Behavior of fluorescein-labelled 12mer arrays (FX-arrays).
Figure S6. Reversibility of tau-12mer array binding.
Figure S7. Extended tau-DNA binding data.
Figure S8. Binding and LLPS behavior of tau DNA-binding region deletion constructs.
Figure S9. Extended tau-array MAS-NMR data.
Figure S10. MS analysis of phosphorylated tau.
Figure S11. Post-translational modification (PTM) mapping of phosphorylated constructs using LC-MS/MS.
Figure S12. Binding and LLPS behavior of phosphorylated tau.
Figure S13. Analysis of HP1α and pHP1α.
Figure S14. LLPS behavior of tau with pHP1α.
Figure S15. Phase separation behavior of unmodified HP1α.
Figure S16. LLPS behavior of phosphorylated tau with pHP1α and 12mer arrays.
Table S1. Maximum recovery (%) and half-time (s) parameters obtained in different FRAP experiments.
Table S2. Dissociation constants (Kd) determined in this study.
Table S3. Phosphorylation sites of tau.

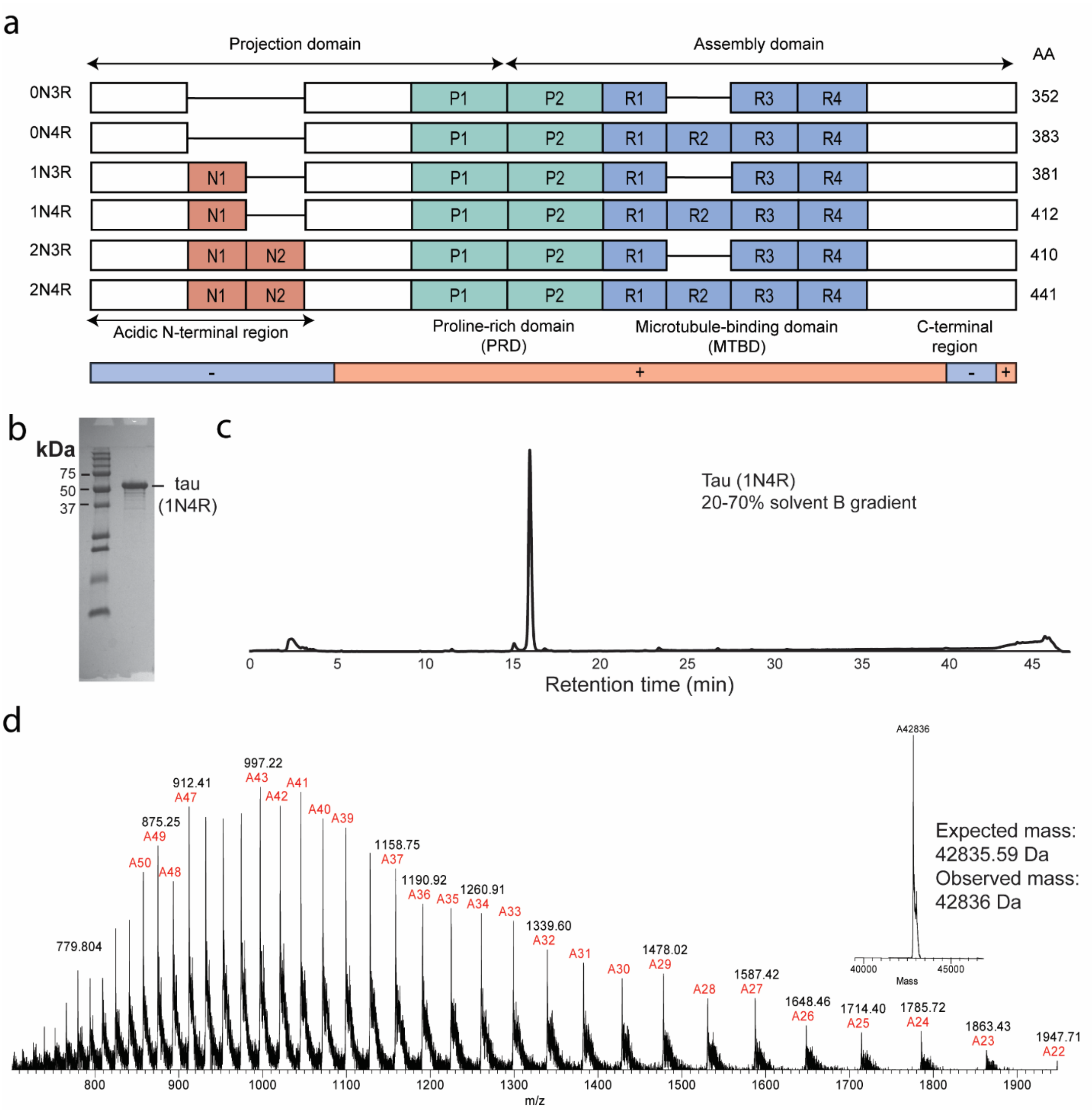


**Figure S1. Analysis of purified 1N4R tau.** a) Domain architecture of the six tau isoforms. b) Purified 1N4R tau analyzed by SDS-PAGE, c) RP-HPLC, and d) mass spectrometry.

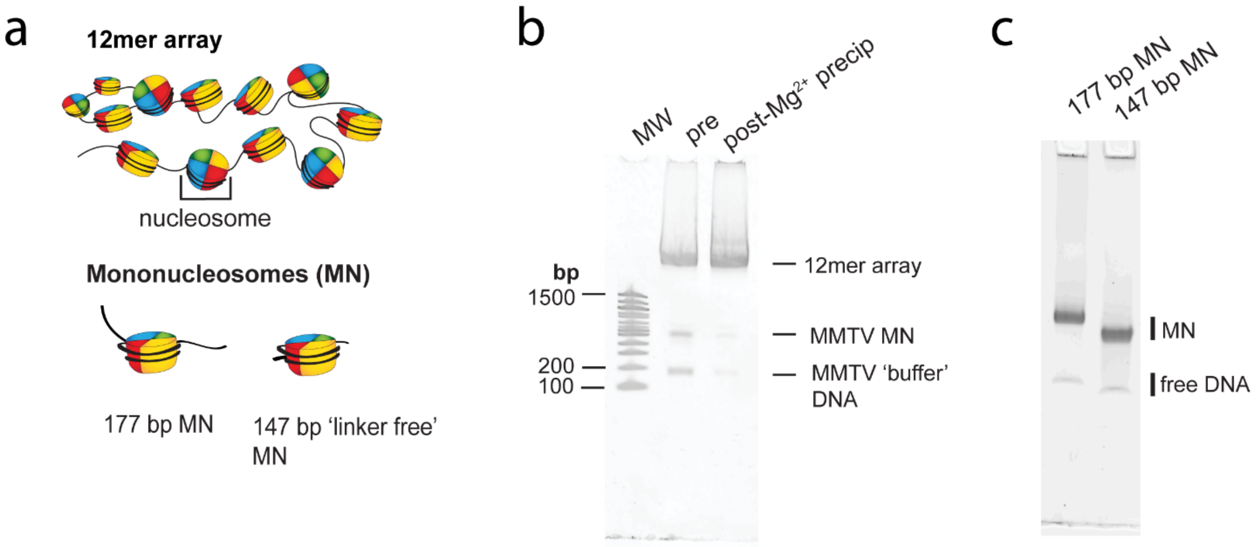


**Figure S2. Nucleosome and 12mer array constructs.** a) Schematic of 12mer array and mononucleosome constructs. b) 12mer array assembly before and after $Mg^{2+}$ purification as analyzed by APAGE gel (2% acrylamide, 1% agarose). c) 147 bp and 177 bp mononucleosomes on a native 5% TBE gel. MW – molecular weight.

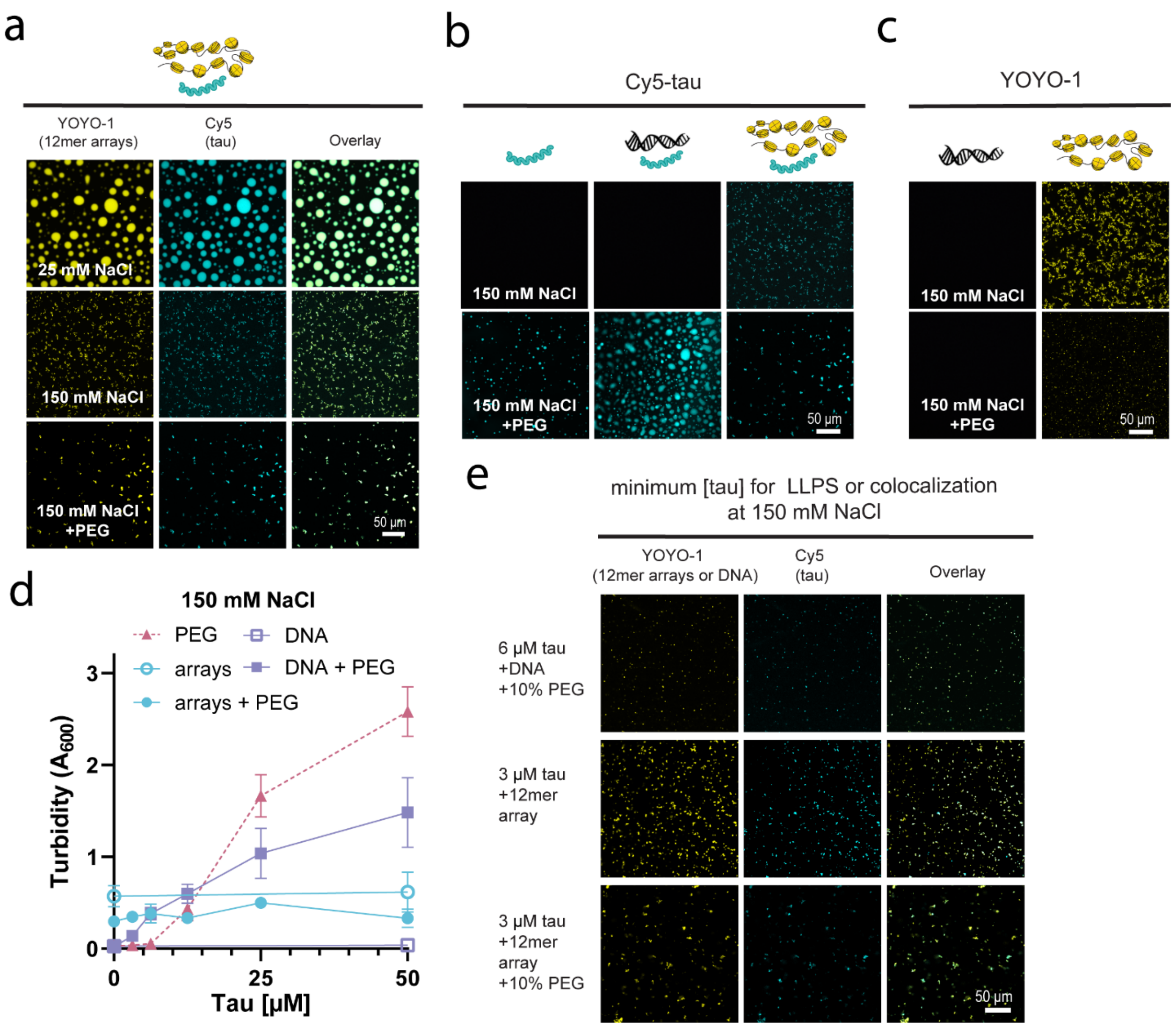


**Figure S3. Behavior of tau under crowding conditions.** a) Confocal microscopy images of 50 µM tau with 80 nM 12mer arrays (1 µM equivalent of mononucleosomes) with 0.8 µM YOYO-1 in 20 mM HEPES buffer, 0.5 mM TCEP, pH 7.2, under different salt and crowding conditions, visualized with 5% Cy5-tau (scale bar = 50 µm). b) Confocal microscopy images of 50 µM tau with 80 nM 12mer arrays (1 µM equivalent of mononucleosomes) or 80 nM 2.1 kbp DNA in the same buffer under different salt and crowding conditions, visualized with 5% Cy5-tau and 0.8 µM YOYO-1 (scale bar = 50 µm). c) Confocal microscopy images of 80 nM 12mer arrays (1 µM equivalent of mononucleosomes) or 80 nM 2.1 kbp DNA with 0.8 µM YOYO-1 in the same buffer, under different salt and crowding conditions, visualized with YOYO-1 (scale bar = 50 µm). d) Turbidity (A600) of tau with 10% PEG-6000, 12mer arrays, and DNA, in the same buffer at 150 mM NaCl. e) Confocal microscopy images of the lowest concentration samples used in the turbidity assay of d) where tau is colocalized or phase separated with 80 nM 12mer array or 80 nM 2.1 kbp DNA.

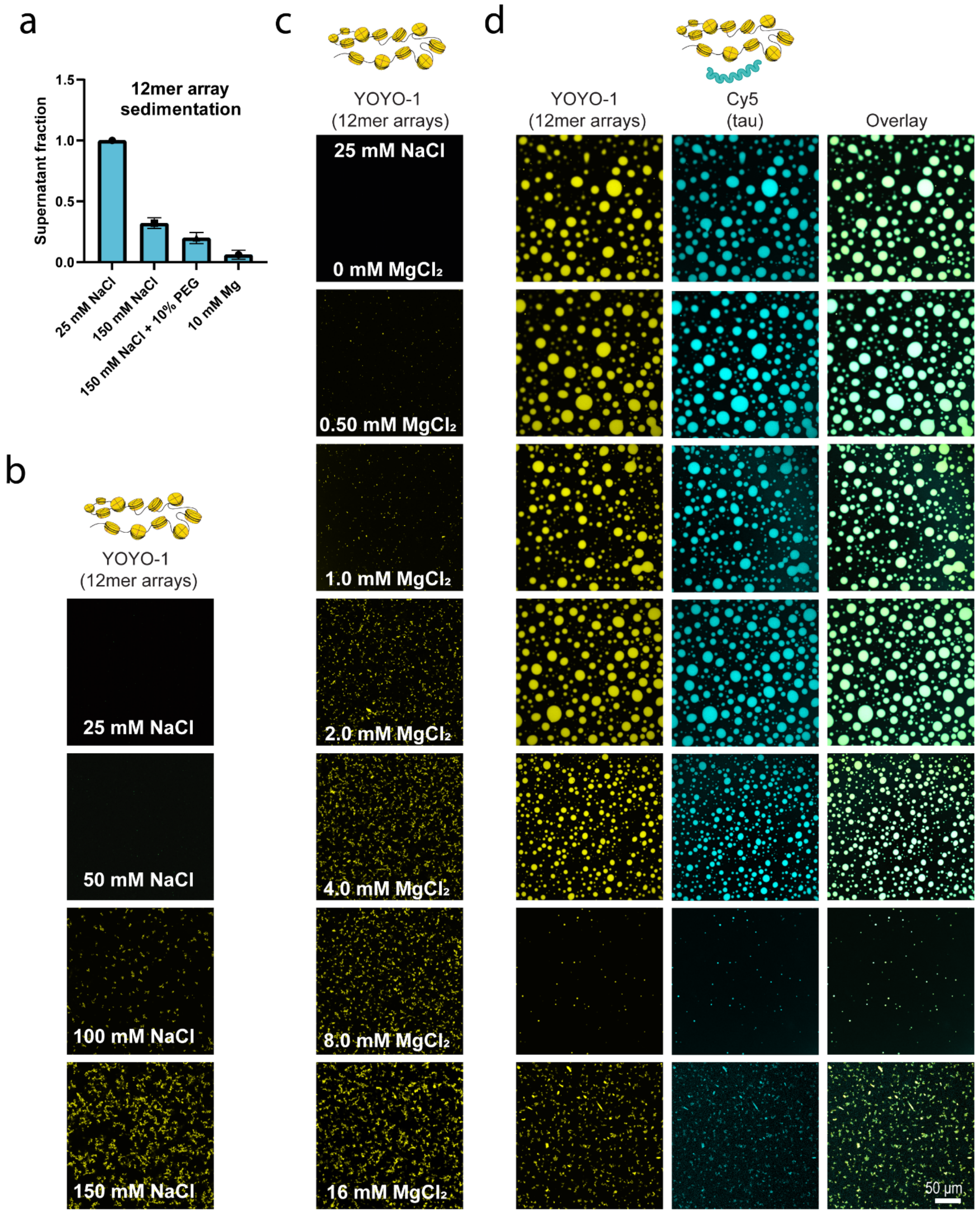


**Figure S4. Effect of mono- and divalent salt on 12mer arrays and tau-array LLPS**. a) Self-association of 12mer arrays under different conditions. 12mer arrays were incubated in different salt or crowding conditions, subjected to low speed centrifugation, and the soluble fraction was calculated by the A260 of the supernatant. The experiment was conducted in 20 mM HEPES buffer, 0.5 mM TCEP, pH 7.2 with the indicated NaCl, PEG or $MgCl_2$ concentrations. b) Confocal microscopy images of 80 nM 12mer arrays (1 μM equivalent of mononucleosomes) with 0.8 μM YOYO-1 with varying NaCl concentrations in the same buffer. c) Confocal microscopy images of 80 nM YOYO-1 labelled arrays with varying $MgCl_2$ concentrations, in the same buffer at 25 mM NaCl. c) Confocal microscopy images of 50 μM tau (5% Cy5-labelled) with 80 nM YOYO-1 labelled arrays with varying $MgCl_2$ concentrations, in the same buffer at 25 mM NaCl. The scale bar in the images denotes 50 μm.

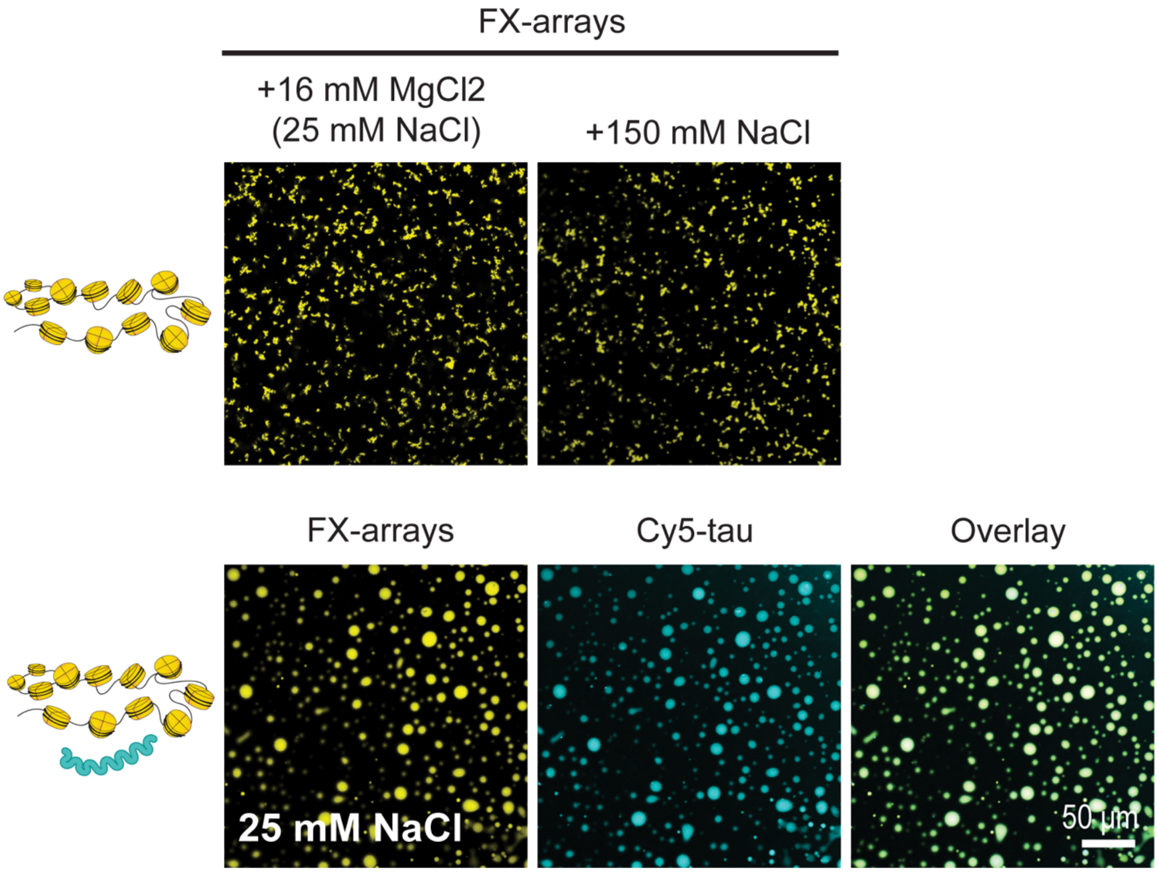


**Figure S5. Behavior of fluorescein-labelled 12mer arrays (FX-arrays).** a) Confocal microscopy images of 80 nM FX-arrays (1 µM equivalent of mononucleosomes), in 20 mM HEPES buffer, 25 mM NaCl, 0.5 mM TCEP, pH 7.2. The upper images of the FX-arrays have 16 mM $MgCl_2$ or 150 mM NaCl added. The lower panel includes 50 µM tau, in the same buffer. The scale bar in the images denotes 50 µm.

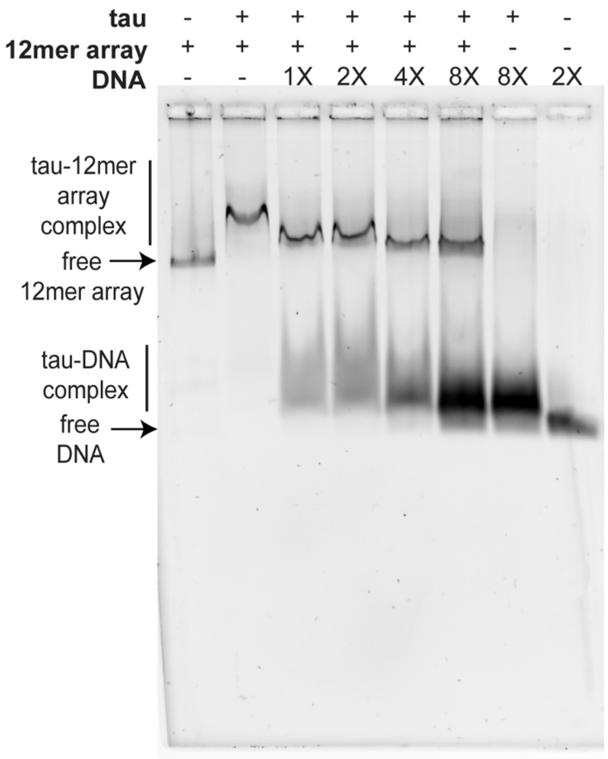


**Figure S6. Reversibility of tau-12mer array binding.** a) 1.7 nM 12mer arrays were incubated with 500 nM tau for 20 minutes, and increasing amounts (20 nM/1X, 40 nM/2X, 80 nM/4X, 160 nM/8X) of competitor 177 bp DNA were added, incubated for 20 minutes, and then subjected to electrophoresis on an APAGE composite gel (2% acrylamide, 1% agarose). All experiments were performed in 20 mM HEPES buffer, 150 mM NaCl, 0.5 mM TCEP, 0.1% Tween, pH 7.2.

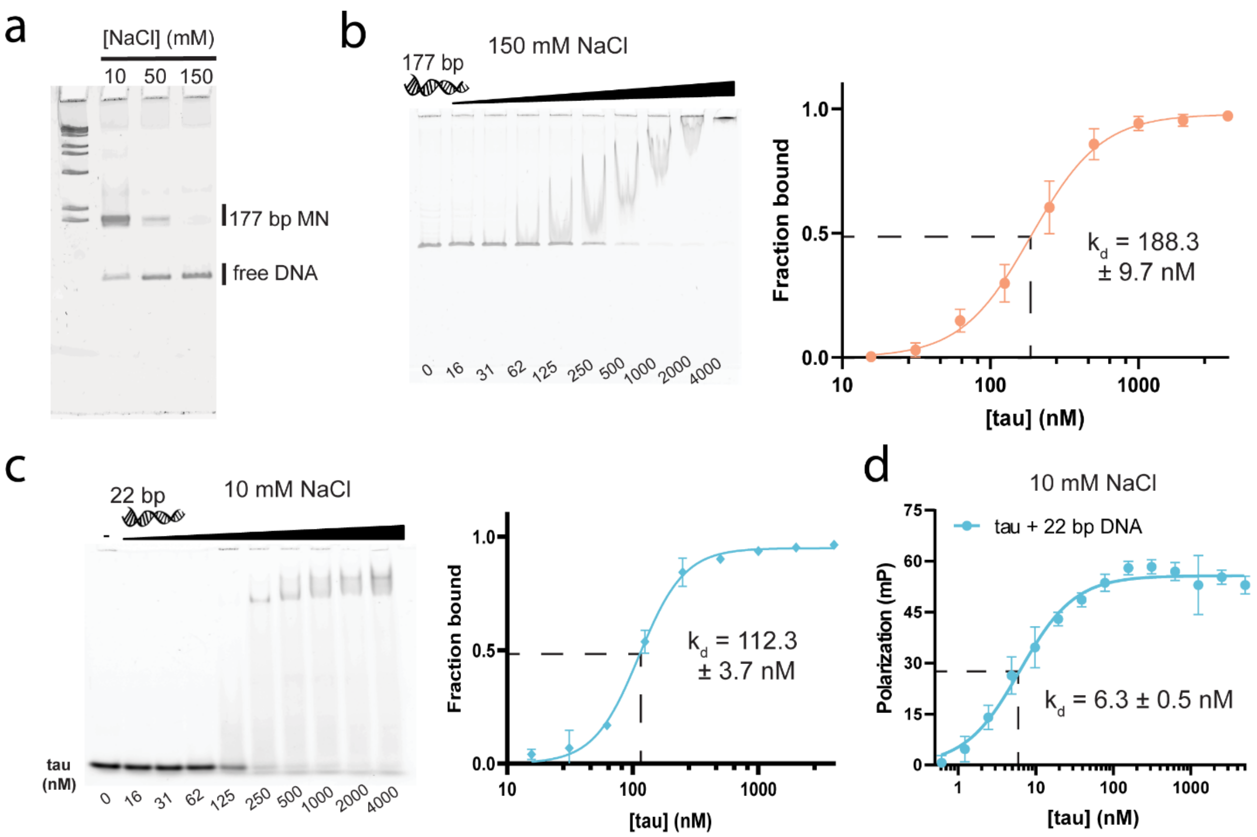


**Figure S7. Extended tau-DNA binding data.** a) Salt-induced dissociation of 20 nM 177 bp mononucleosomes on a native 5% TBE gel, in 20 mM HEPES buffer, 0.5 mM TCEP, pH 7.2, with different NaCl concentrations. High concentrations of NaCl can promote the dissociation of mononucleosomes present at low concentrations into octamer and free DNA. b) (left) EMSA of 20 nM 177 bp DNA with various concentrations of tau, in the same buffer with 0.1% Tween and at 150 mM NaCl. (right) Quantification of binding propensity based on the intensity of the unbound DNA bands. c) (left) EMSA of 20 nM 22 bp DNA with various concentrations of tau, in the same buffer with 0.1% Tween and 10 mM NaCl. (right) Quantification of binding propensity based on the intensity of the unbound DNA bands. d) Fluorescence anisotropy measurements of binding of 10 nM fluorescein-labelled 22 bp DNA with varying concentrations of tau, in the same buffer with 0.01% nonidet P40 substitute and 10 mM NaCl.

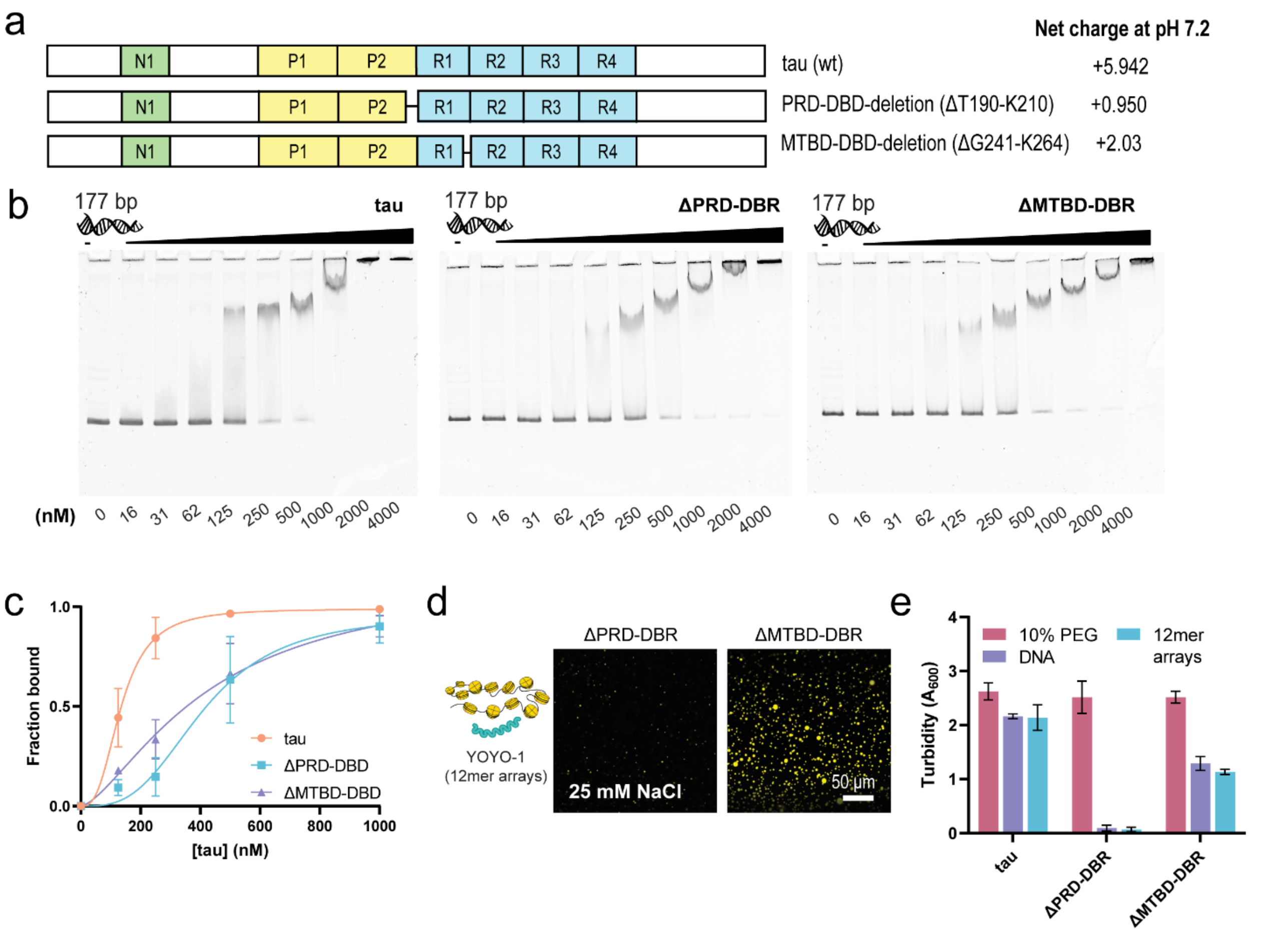


**Figure S8. Binding and LLPS behavior of tau DNA-binding region deletion constructs.** a) Schematic of tau (1N4R) domain architecture and domain deletions. b) EMSA of 20 nM 177 bp DNA with various concentrations of tau and deletion constructs, in 20 mM HEPES buffer, 10 mM NaCl, 0.5 mM TCEP, 0.1% Tween, pH 7.2. c) Quantification of binding in b) calculated using the unbound fraction. d) Confocal microscopy images of 50 μM deletion constructs with 80 nM 12mer arrays visualized with 0.8 μM YOYO-1, in the same buffer with 25 mM NaCl. The scale bar in the images denotes 50 μm. e) Turbidity (A600) of tau and deletion constructs, with LLPS induced by 10% PEG-6000, 12mer arrays, and DNA, in the same low salt buffer with 25 mM NaCl.

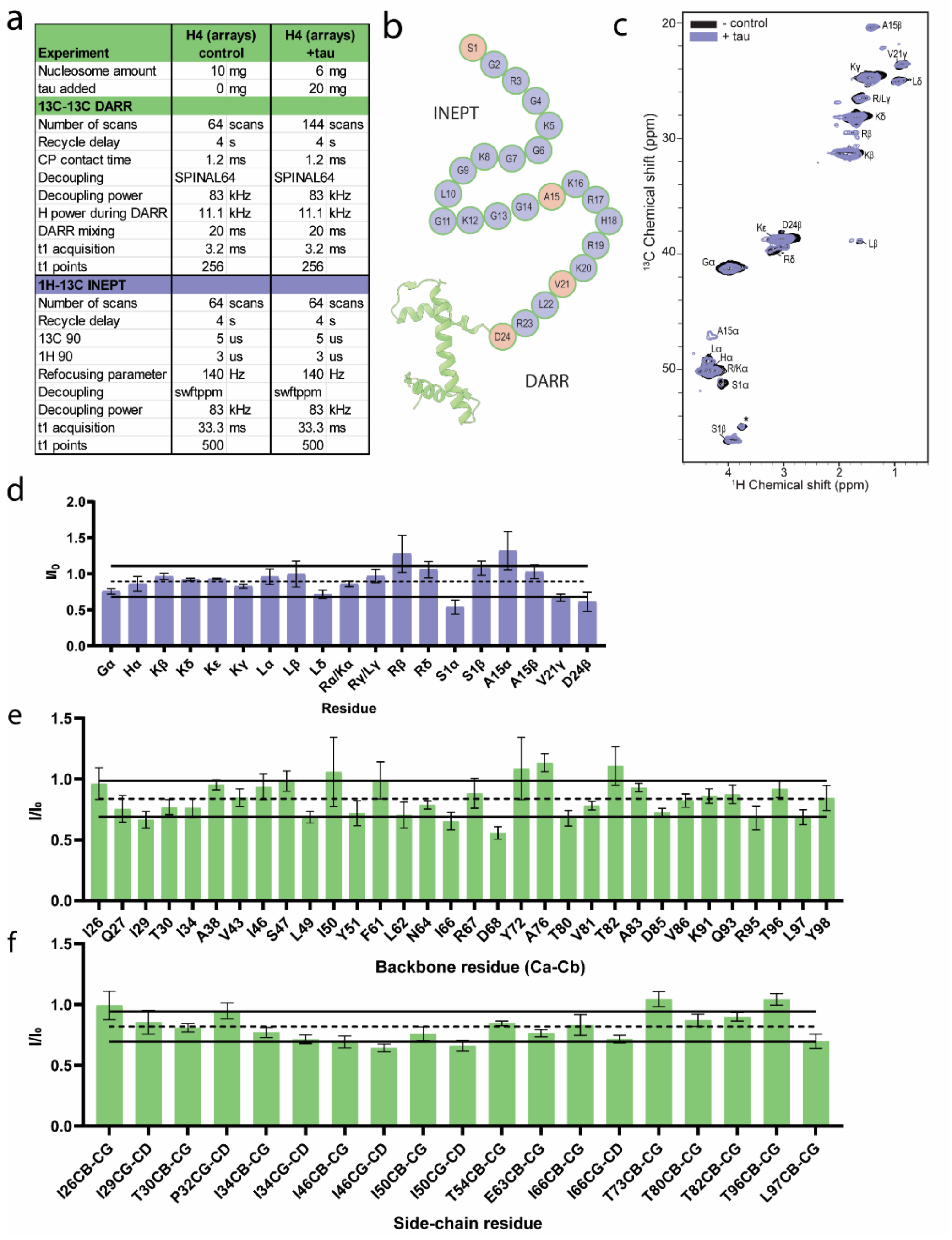

| Experiment | H4 (arrays) control | H4 (arrays) +tau |
|---|---|---|
| Nucleosome amount | 10 mg | 6 mg |
| tau added | 0 mg | 20 mg |
| 13C-13C DARR | | |
| Number of scans | 64 scans | 144 scans |
| Recycle delay | 4 s | 4 s |
| CP contact time | 1.2 ms | 1.2 ms |
| Decoupling | SPINAL64 | SPINAL64 |
| Decoupling power | 83 kHz | 83 kHz |
| H power during DARR | 11.1 kHz | 11.1 kHz |
| DARR mixing | 20 ms | 20 ms |
| t1 acquisition | 3.2 ms | 3.2 ms |
| t1 points | 256 | 256 |
| 1H-13C INEPT | | |
| Number of scans | 64 scans | 64 scans |
| Recycle delay | 4 s | 4 s |
| 13C 90 | 5 us | 5 us |
| 1H 90 | 3 us | 3 us |
| Refocusing parameter | 140 Hz | 140 Hz |
| Decoupling | swftppm | swftppm |
| Decoupling power | 83 kHz | 83 kHz |
| t1 acquisition | 33.3 ms | 33.3 ms |
| t1 points | 500 | 500 |



**Figure S9. Extended tau-array MAS NMR data.** a) Tabulated experimental parameters of the MAS NMR experiments of 12mer arrays with and without tau. b) Amino acid composition of the histone H4 tail. Residues marked in orange have unambiguous assignments in the INEPT spectrum. c) 2D $^{1}H$-$^{13}C$ INEPT spectra of 12mer arrays with tau (purple) and without tau (black). Asterisk (*) indicates the Tris buffer peak. d) Ratio of the peak intensities of the tau spectrum divided over the peak intensities of the control spectrum. e) Intensity ratio (bound/free) of one-bond backbone Cα-Cβ correlation peaks in the 2D $^{13}C$-$^{13}C$ DARR spectrum in Fig. 4b. Intensities were calculated only for non-overlapped peaks. f) Intensity ratio (bound/free) of resolved side-chain one-bond correlation peaks in the 2D $^{13}C$-$^{13}C$ DARR spectrum in Fig. 4b.

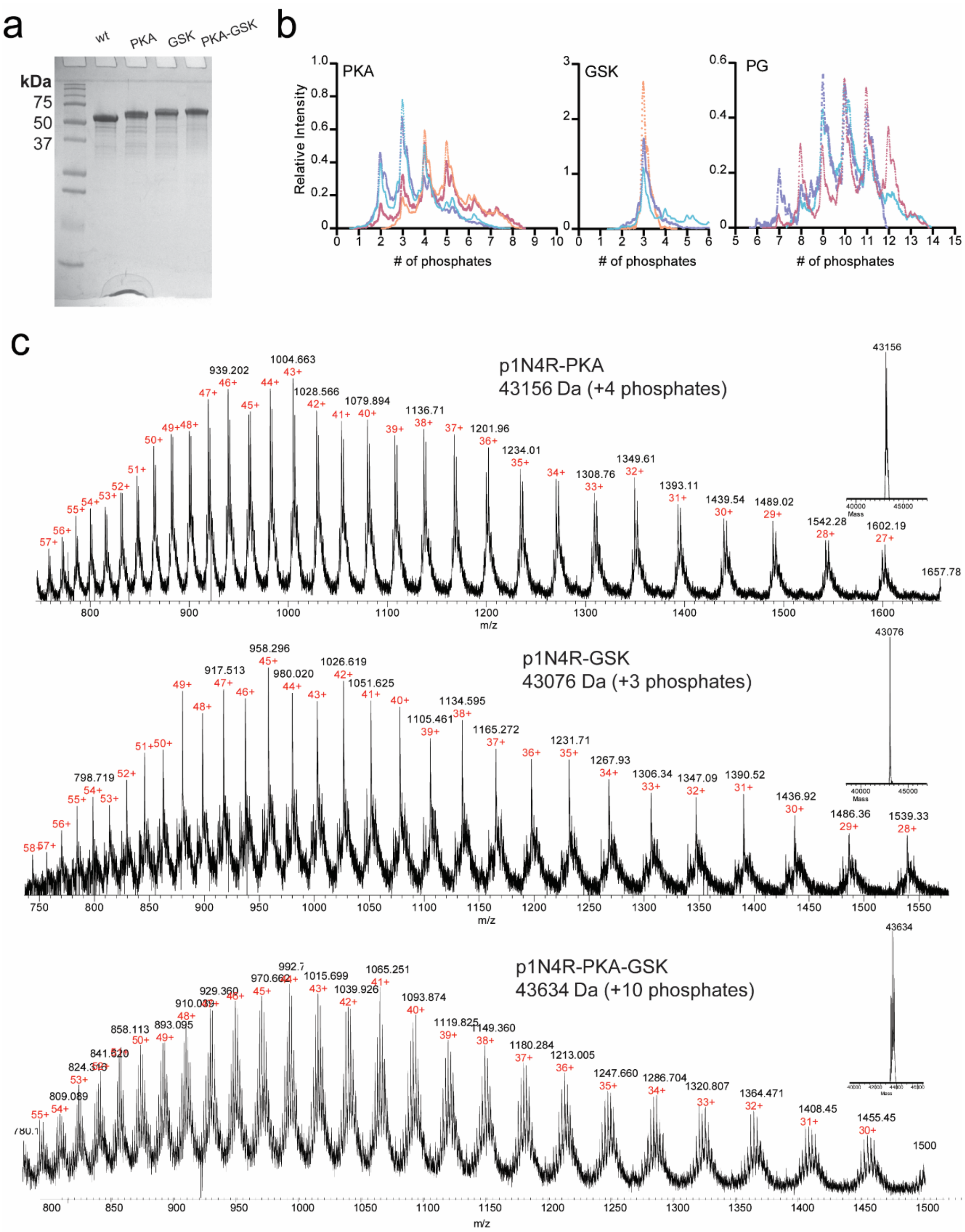


**Figure S10. MS analysis of phosphorylated tau.** a) Full image of the 18% SDS-PAGE gel of phoshorylated tau constructs presented in Fig. 5a. b) Relative abundance of the number of phosphates detected on tau phosphorylated by PKA, GSK, and PKA/GSK, respectively. Individual replicates are shown (n=3 for PKA or GSK phosphorylated tau and 4 with PKA-GSK phosphorylated tau) for independent phosphorylation reactions. Relative intensity is calculated by dividing the raw intensity by the area under the mass spectrum curve as approximated by the trapezoidal rule, to correct for differences in sample. c) Mass spectrometry analysis of a representative replicate for each phosphorylated form of tau.

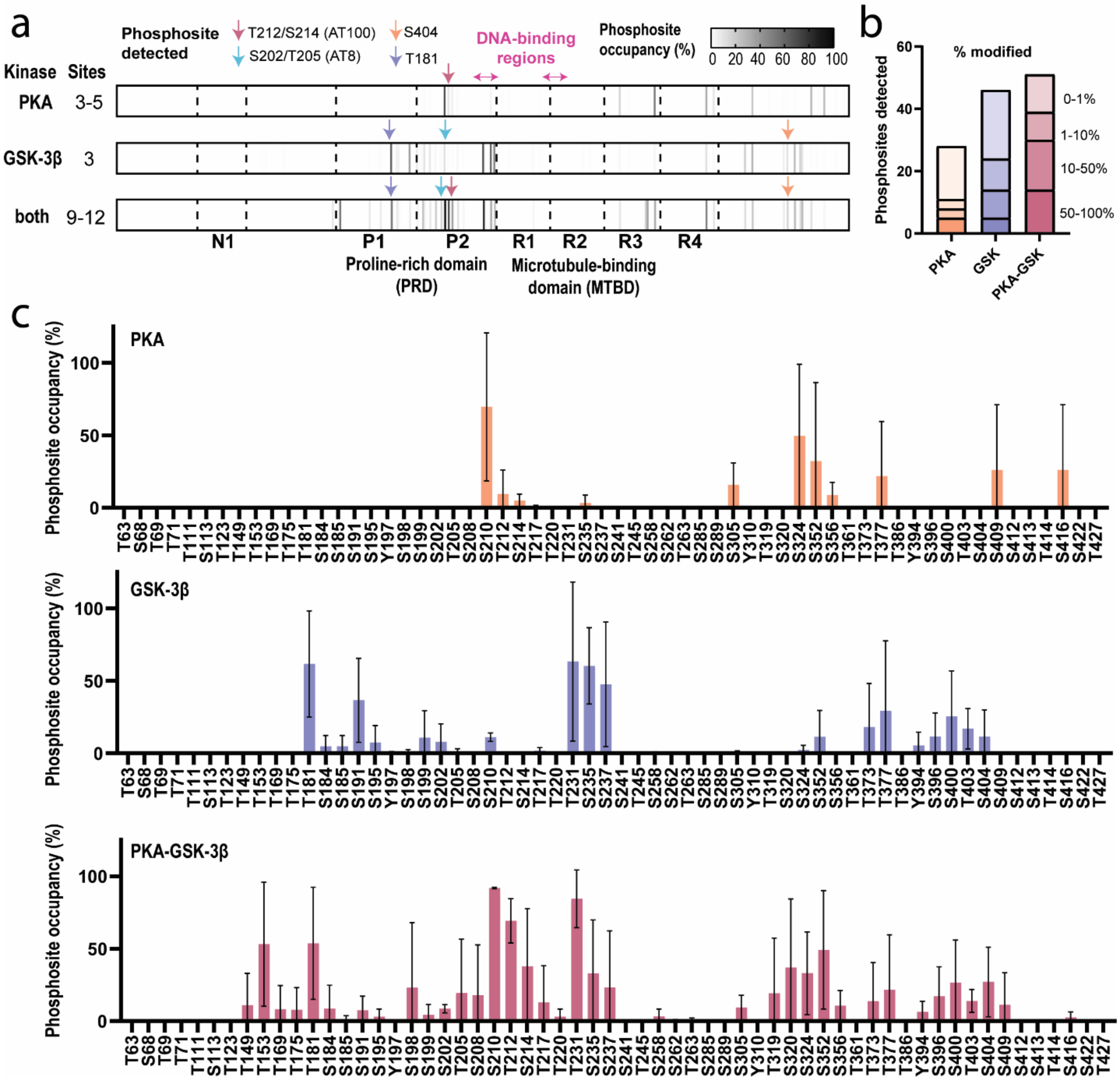


**Figure S11. Post-translational modification (PTM) mapping of phosphorylated constructs using LC-MS/MS.** a) Schematic of tau domain architecture and detected phosphorylation sites, as colored by phosphorylation site relative occupancy percentage. b) Number of phosphorylated sites detected by post-translational modification mapping using digestion with trypsin, as colored by the percent of modified phosphorylated peptide that was detected. c) Quantification of relative phosphorylation site occupancy, as labelled by the 2N4R nomenclature, for greater ease in comparison to the bulk of tau literature. Error bars are standard deviation of three replicates for PKA and GSK phosphorylated tau, and four replicates for PKA-GSK phosphorylated tau. For 1N4R tau nomenclature, see Table S3.

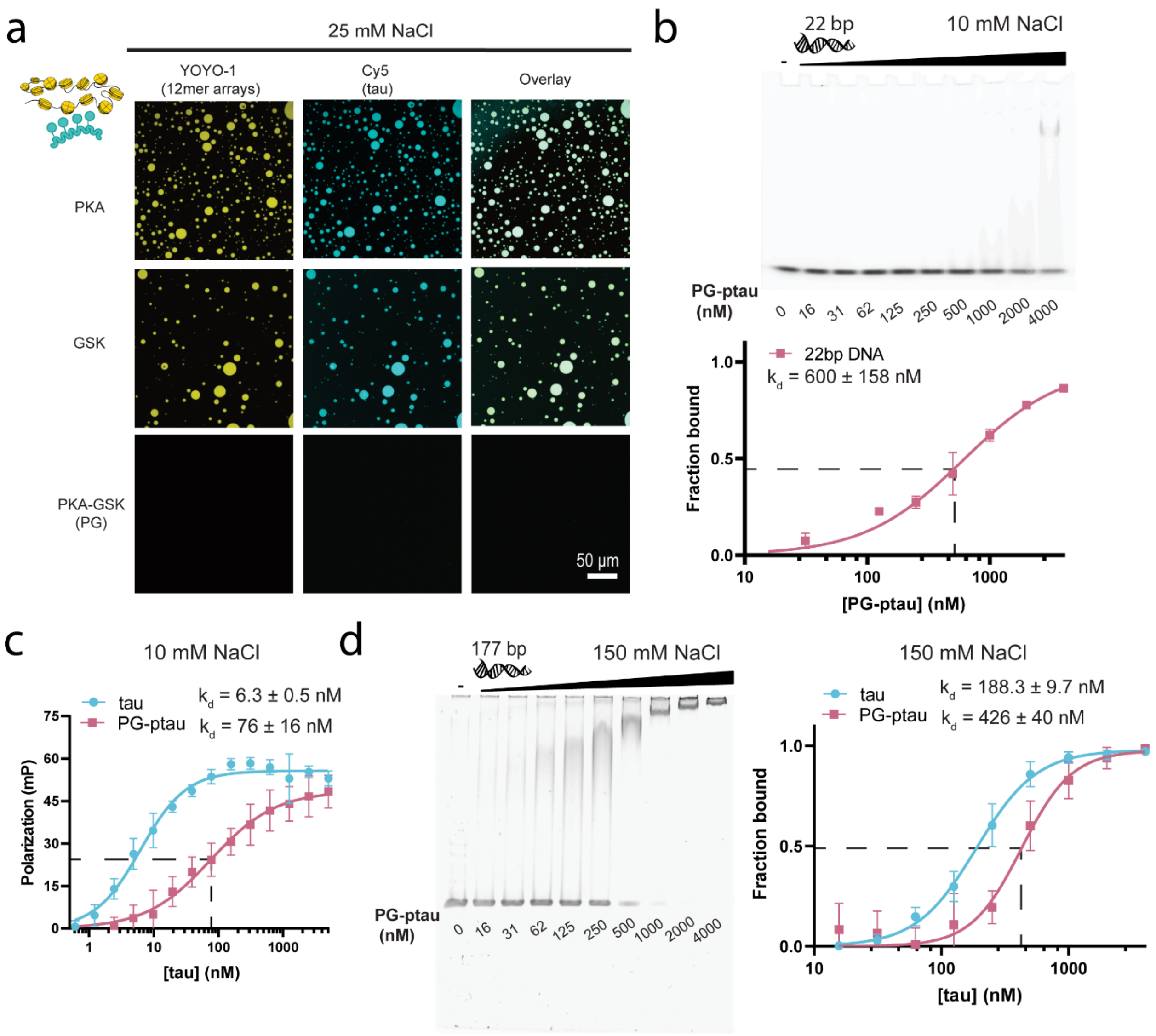


**Figure S12. Binding and LLPS behavior of phosphorylated tau.** a) Confocal microscopy images of 50 µM phosphorylated tau and 80 nM 12mer arrays (1 µM mononucleosome equivalent), spiked with 5% Cy5-labeled phosphorylated tau and 0.8 µM YOYO-1 (scale bar = 50 µm). Studies were conducted in 20 mM HEPES buffer, 25 mM NaCl, 0.5 mM TCEP, pH 7.2. b) (top) EMSA to assess the binding propensity of PKA-GSK phosphorylated tau with 20 nM 22 bp DNA in the same buffer with 0.1% Tween at 10 mM NaCl. (bottom) Quantification of binding propensity based on the intensity of the unbound DNA. Error bars represent the standard deviation from three independent EMSA experiments. c) Fluorescence anisotropy measurements of binding of 10 nM fluorescein-labelled 22 bp DNA with varying concentrations of wild-type or PKA-GSK phosphorylated tau, with 0.01% nonidet P40 substitute at 10 mM NaCl. The tau binding curve data (blue) is repeated from Fig. S7d. d) (left) EMSA of 20 nM 177 bp DNA with various concentrations of tau, with 0.1% Tween at 10 mM NaCl. (right) Quantification of binding propensity based on the intensity of the unbound DNA bands. Error bars represent the standard deviation from three independent EMSA experiments. The tau binding curve data (blue) is repeated from Fig. S7b. PG-ptau – tau phosphorylated by a combination of PKA and GSK-3β.

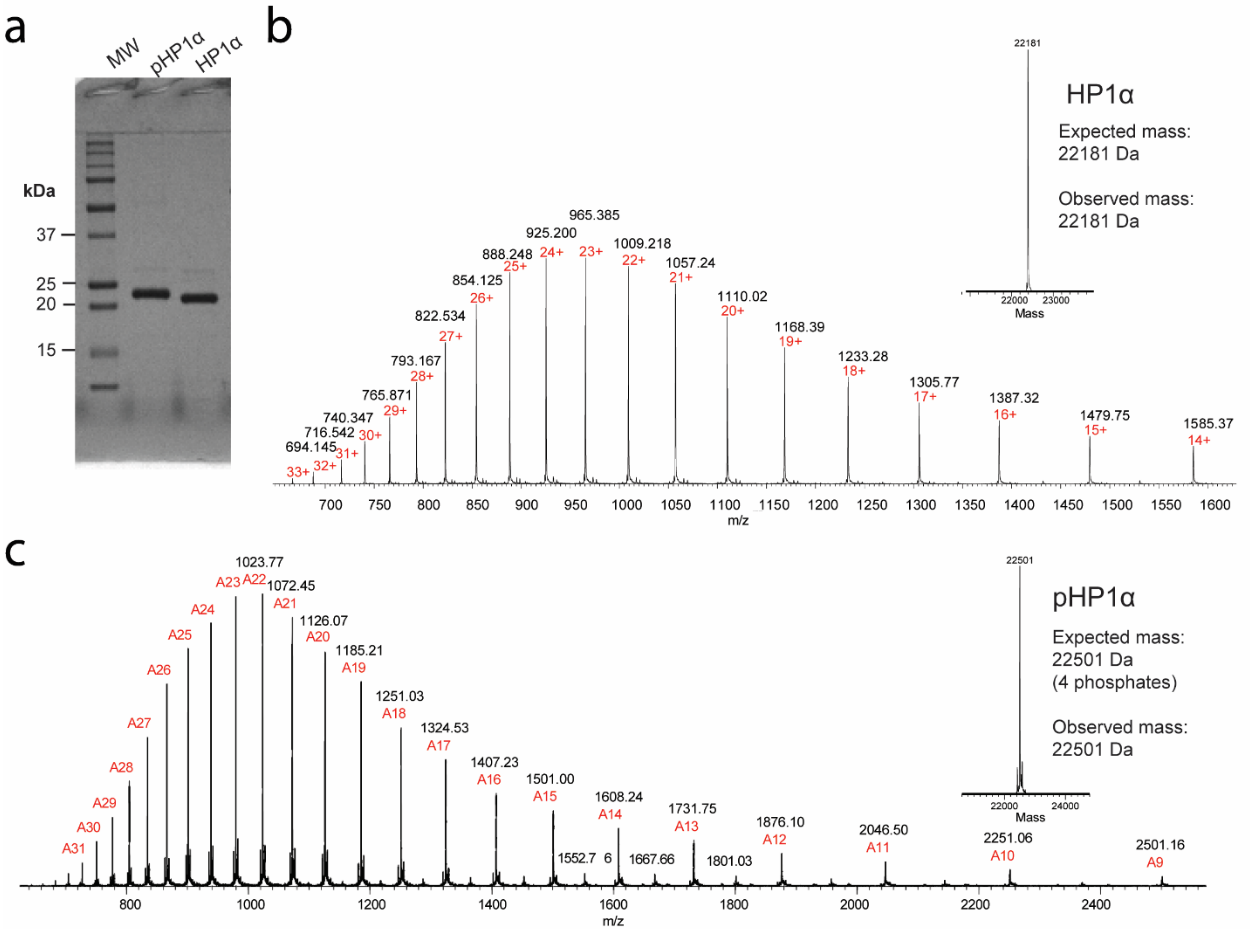


**Figure S13. Analysis of HP1α and pHP1α.** a) Coomassie-stained 12% SDS-PAGE gel of pHP1α and HP1α. b) ESI-MS analysis of HP1α. c) ESI-MS analysis of pHP1α.

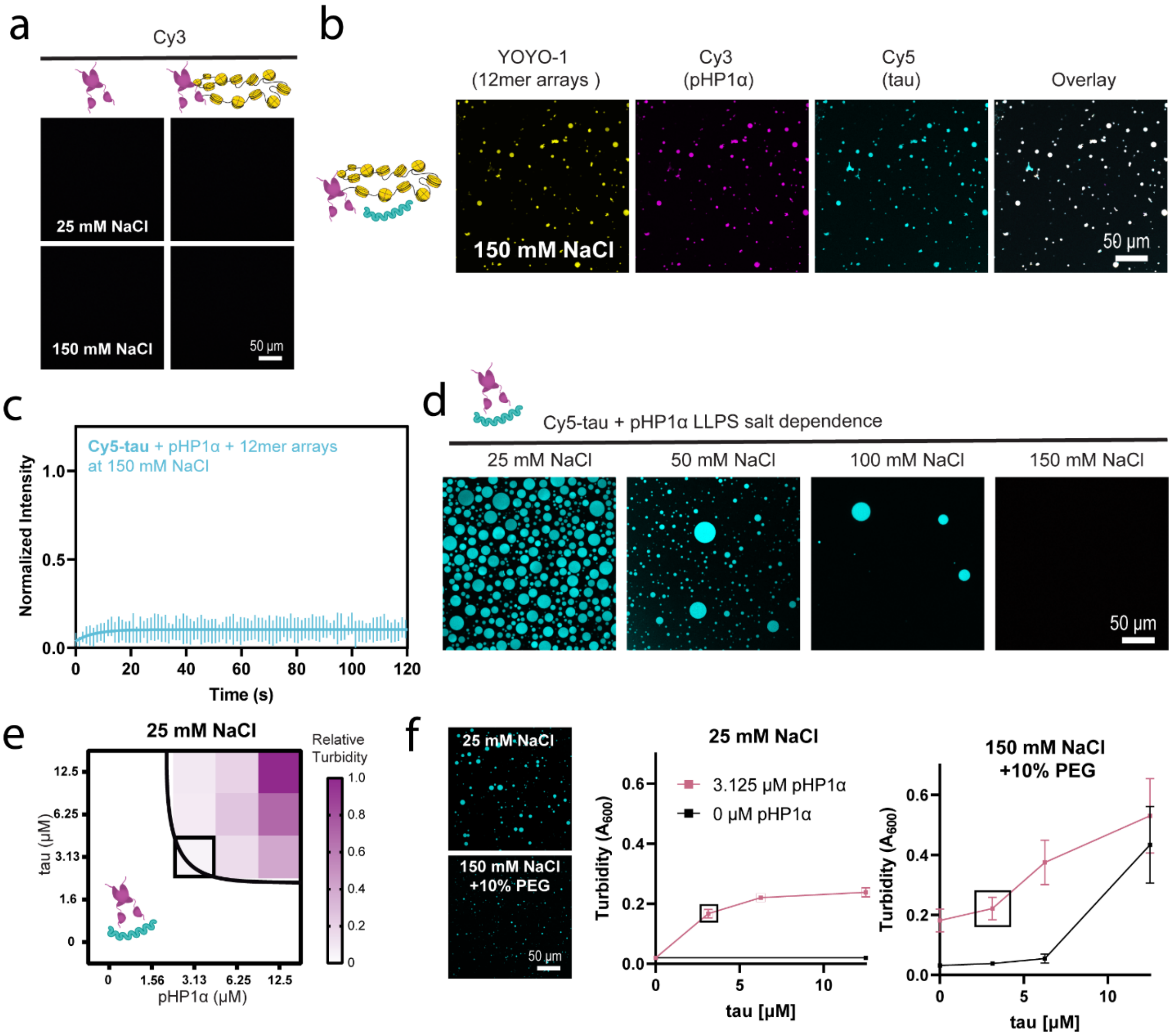


**Figure S14. LLPS behavior of tau with pHP1α.** a) Confocal microscopy images of 50 µM pHP1α with or without 80 nM 12mer arrays, visualized with 5% Cy3-pHP1α and 0.8 µM YOYO-1, in 20 mM HEPES buffer, 0.5 mM TCEP, pH 7.2, with varying NaCl concentration. At 150 mM NaCl, 12mer arrays are solid and compacted, but pHP1α does not associate with it (12mer array localization not shown). b) Confocal microscopy images of 50 µM tau, 50 µM pHP1α, and 80 nM 12mer arrays, spiked with 5% Cy5-tau, 5% Cy3-pHP1α, and labelled with 0.8 µM YOYO-1 (Scale bar = 50 µm), in the same buffer with 150 mM NaCl. c) FRAP analysis of tau mobility in samples containing 50 µM tau with pHP1α and 12mer arrays in the same high salt buffer. The standard deviation and the best monoexponential fit curve are plotted for the data set. d) Confocal microscopy images of 50 µM tau and 50 µM pHP1α in the same buffer, at varying salt concentrations. Images are visualized with 5% Cy5-tau and 5% Cy3-pHP1α (Scale bar = 50 µm). e) Phase diagram of tau-pHP1α in the same buffer with 25 mM NaCl, with turbidity measured by A600 plotted as indicated by the gradient scale. Values were normalized to the largest turbidity measurement. f) Turbidity (A600) of increasing concentrations of tau with or without 3 µM pHP1α, in the same buffer with 25 mM NaCl or 150 mM NaCl and 10% PEG-6000. On the left, boxed data points are shown as confocal microscopy images of 3 µM tau and 3 µM pHP1α at the indicated conditions.

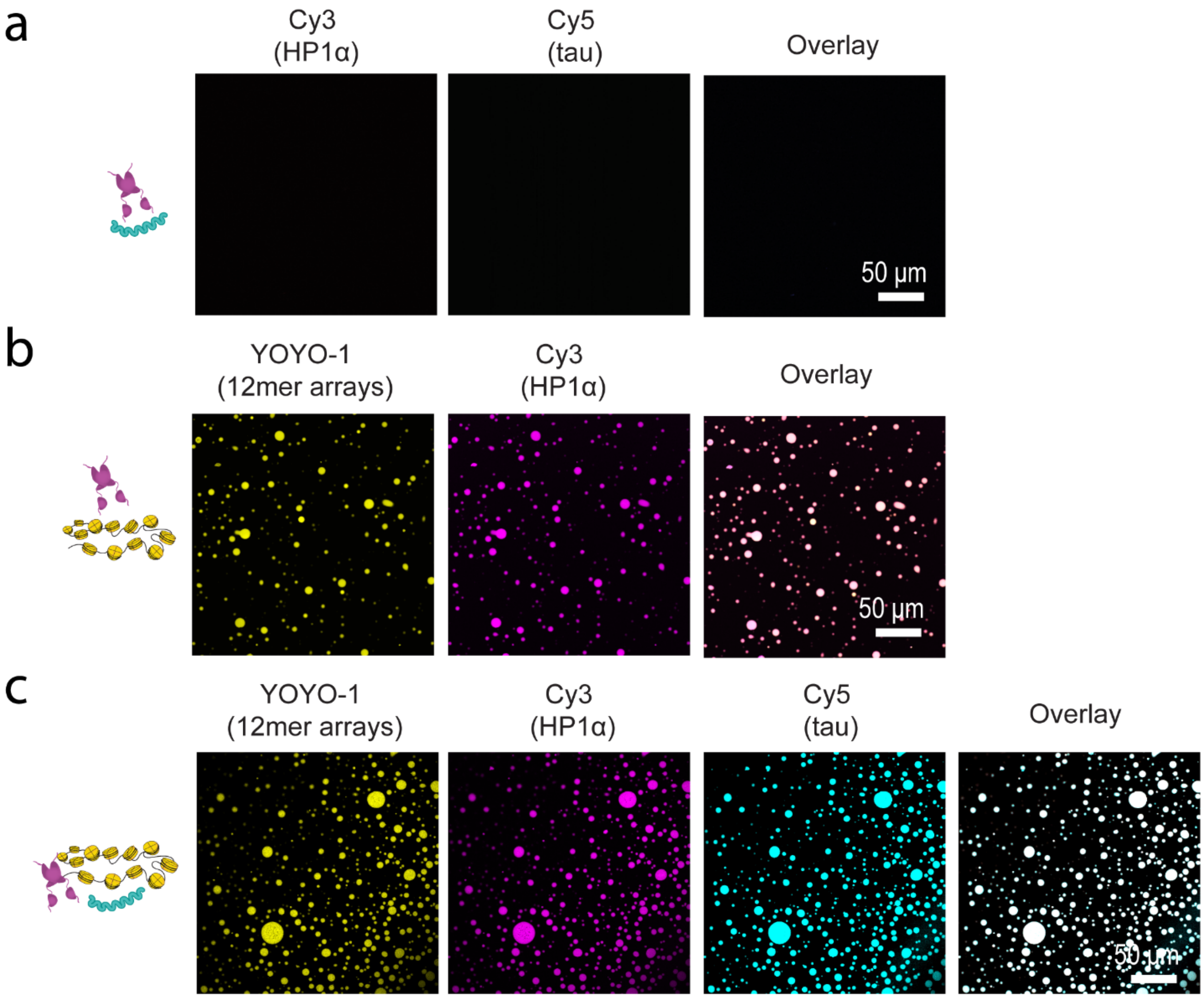


**Figure S15. Phase separation behavior of unmodified HP1α.** a) Confocal microscopy images of 50 μM HP1α and 50 μM tau in 20 mM HEPES buffer, 25 mM NaCl, 0.5 mM TCEP, pH 7. b) Confocal microscopy images of 50 μM HP1α and 80 nM 12mer arrays (1 μM equivalent of mononucleosomes) labelled with 0.8 μM YOYO-1 in the same low salt buffer. c) Confocal microscopy images of 50 μM tau, 50 μM HP1α, and 80 nM 12mer arrays (1 μM mononucleosomes) labelled with 0.8 μM YOYO-1 in in the same low salt buffer. Different components were visualized with 5% Cy5-tau, 5% Cy3-HP1α, and 0.8 μM YOYO-1 (Scale bar = 50 μm).

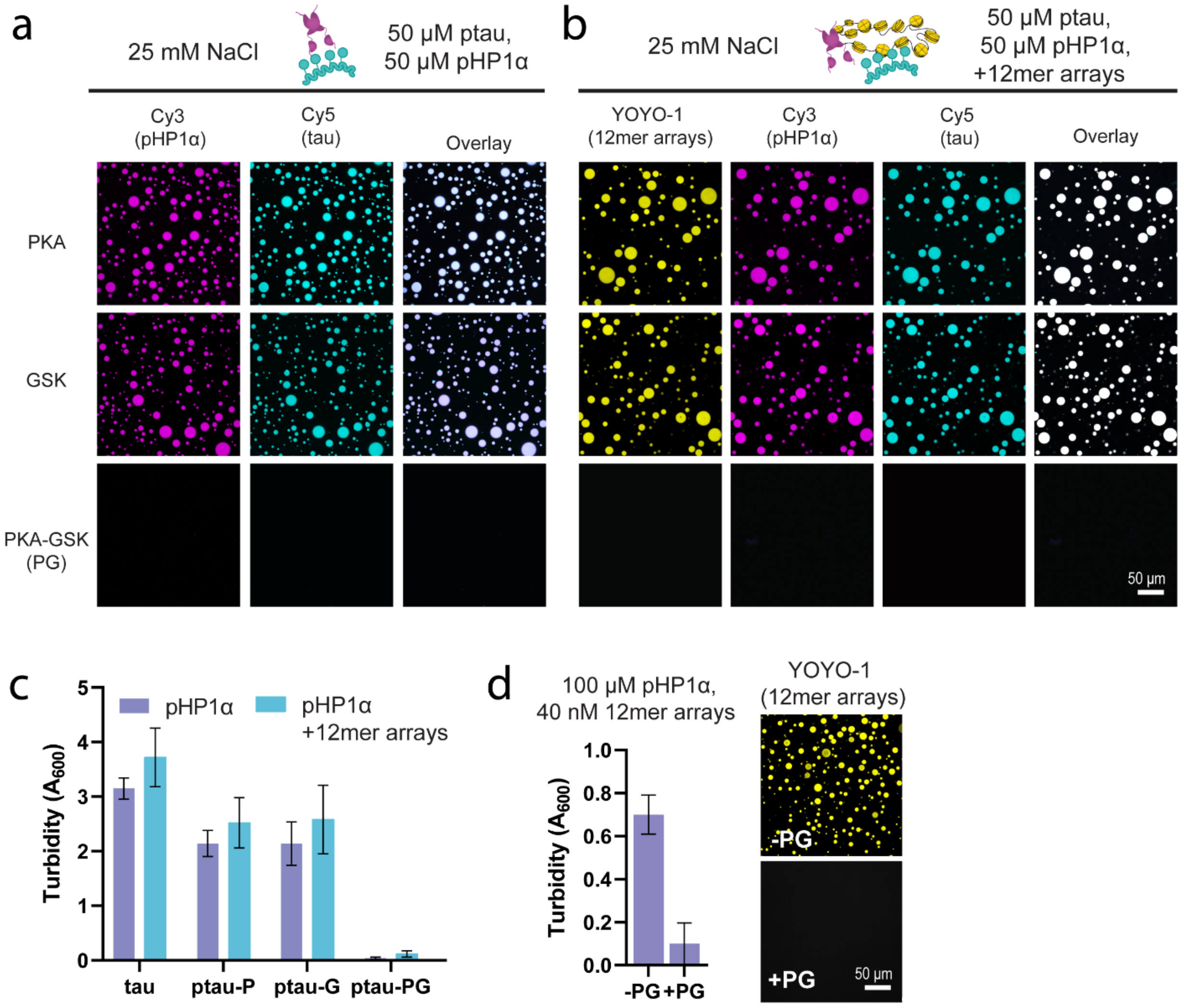


**Figure S16. LLPS behavior of phosphorylated tau with pHP1α and 12mer arrays.** a) Confocal microscopy images of 50 μM phosphorylated tau and 50 μM pHP1α, spiked with 5% Cy5-ptau and 5% Cy3-pHP1α (Scale bar = 50 μm). Studies were conducted in 20 mM HEPES buffer, 25 mM NaCl, 0.5 mM TCEP, pH 7.2. b) Confocal microscopy images of 50 μM phosphorylated tau, 50 μM pHP1α, 80 nM 12mer arrays (1 μM mononucleosome equivalent), spiked with 5% Cy5-ptau, 5% Cy3-pHP1α, and 0.8 μM YOYO-1 (Scale bar = 50 μm). Studies were conducted in the same low salt buffer. c) $A_{600}$ turbidity measurements of 50 μM phosphorylated tau with 50 μM pHP1α with and without 80 nM 12mer arrays, in the same low salt buffer. d) (left) Phase separation of 100 μM pHP1α and 40 nM 12mer arrays (0.5 μM mononucleosome equivalent) with and without 50 μM PKA-GSK phosphorylated tau, as measured by turbidity at $A_{600}$. (right) Confocal microscopy images of 100 μM pHP1α and 40 nM 12mer arrays (0.5 μM mononucleosome equivalent) with and without 50 μM PKA-GSK phosphorylated tau, visualized with 0.8 μM YOYO-1 (Scale bar = 50 μm). Both were conducted in the same low salt buffer.

**Table S1. Maximum recovery (%) and half-time (s) parameters obtained in different FRAP experiments.**

| LLPS condition | Fluorophore POV | Figure | Maximum recovery (%) | SE | 95% CI Range | Half-life ($t_{1/2}$) (s) | 95% CI Range |
|---|---|---|---|---|---|---|---|
| Tau + 12mer arrays | Cy5-tau | 1b | 39.9 | 0.4 | 39.14 to 40.69 | 17.0 | 15.30 to 19.01 |
| Tau + 2.1 kbp DNA | Cy5-tau | 1b | 57.4 | 0.4 | 56.56 to 58.23 | 20.0 | 18.65 to 21.57 |
| Tau + MNs | Cy5-tau | 1b | 33.9 | 0.3 | 33.23 to 34.68 | 23.3 | 21.34 to 25.61 |
| Tau + 10% PEG | Cy5-tau | 1b | 87.4 | 0.2 | 86.96 to 87.75 | 10.1 | 9.751 to 10.48 |
| Tau + pHP1α + 12mer arrays | Cy5-tau | 6c | 49.4 | 0.5 | 48.47 to 50.34 | 27.2 | 25.42 to 29.14 |
| Tau + pHP1α | Cy5-tau | 6c | 82.1 | 0.2 | 81.68 to 82.47 | 13.3 | 12.87 to 13.77 |

*POV – point of view
*SE – standard error*
*CI – confidence interval*
*MNs – mononucleosomes*

**Table S2. Dissociation constants ($K_d$) determined in this study.**

| Binding partners | $K_d$ (nM) | Method | NaCl (mM) | Figure |
|---|---|---|---|---|
| Tau + 22 bp DNA | 112.3 ± 3.7 | EMSA | 10 | S7c |
| Tau + 22 bp DNA | 6.3 ± 0.5 | FA | 10 | S7d |
| Tau + 177 bp DNA | 134.6 ± 8.7 | EMSA | 10 | 2b,c |
| Tau + 177 bp DNA | 188.3 ± 9.7 | EMSA | 150 | S7b |
| Tau + 177 bp MN | 106 ± 15 | EMSA | 10 | 2a,c |
| Tau + 147 bp MN | 83 ± 13 | EMSA | 10 | 2a,c |
| ΔMTBD-DBR + 177 bp DNA | 427 ± 150 | EMSA | 10 | S8b,c |
| ΔPRD-DBR + 177 bp DNA | 410 ± 52 | EMSA | 10 | S8b,c |
| PG-ptau + 22 bp DNA | 600 ± 158 | EMSA | 10 | S12b |
| PG-ptau + 22 bp DNA | 76 ± 16 | FA | 10 | S12c |
| PG-ptau + 177 bp DNA | 241 ± 14 | EMSA | 10 | 5f |
| PG-ptau + 177 bp DNA | 426 ± 40 | EMSA | 150 | S12d |
| PG-ptau + 177 bp MN | 166 ± 28 | EMSA | 10 | 5f |

*EMSA – electrophoretic mobility shift assay
*FA – fluorescence anisotropy
*PG-ptau – tau phosphorylated with a combination of PKA and GSK-3β
*MN - mononucleosome

**Table S3. Phosphorylation sites of PKA (P), GSK (G) or PKA-GSK (PG) phosphorylated tau detected in different replicates (1 to 4).**

| Residue | | Phosphosite occupancy (%) | | | | | | | | | |
|---|---|---|---|---|---|---|---|---|---|---|---|
| 2N4R | 1N4R | G1 | G2 | G3 | P1 | P2 | P3 | PG1 | PG2 | PG3 | PG4 |
| T63 | 62 | 0.00 | 0.00 | 0.00 | 0.00 | 0.00 | 0.00 | 0.00 | 0.00 | 0.00 | 0.00 |
| S68 | 67 | 0.00 | 0.00 | 0.00 | 0.00 | 0.00 | 0.00 | 0.00 | 0.00 | 0.00 | 0.00 |
| T69 | 68 | 0.00 | 0.00 | 0.00 | 0.00 | 0.00 | 0.00 | 0.00 | 0.00 | 0.00 | 0.00 |
| T71 | 70 | 0.00 | 0.00 | 0.00 | 0.00 | 0.00 | 0.00 | 0.00 | 0.00 | 0.00 | 0.01 |
| T111 | 81 | 0.19 | 0.05 | 0.05 | 0.00 | 0.00 | 0.00 | 0.00 | 0.02 | 0.00 | 0.00 |
| S113 | 83 | 0.06 | 0.02 | 0.00 | 0.00 | 0.00 | 0.00 | 0.11 | 0.01 | 0.00 | 0.00 |
| T123 | 93 | 0.12 | 0.00 | 0.00 | 0.00 | 0.00 | 0.00 | 0.00 | 0.00 | 0.00 | 0.06 |
| T149 | 119 | 0.00 | 0.00 | 0.00 | 0.00 | 0.00 | 0.00 | 0.00 | 0.00 | 0.00 | 43.89 |
| T153 | 123 | | | | 0.00 | 0.00 | 0.00 | 0.00 | 81.76 | 93.07 | 37.52 |
| T169 | 139 | 0.05 | 0.16 | 0.00 | 0.00 | 0.00 | 0.00 | 0.00 | 0.12 | 0.00 | 32.74 |
| T175 | 145 | 0.54 | 0.70 | 0.00 | 0.00 | 0.00 | 0.00 | 0.33 | 0.56 | 0.11 | 30.69 |
| T181 | 151 | 82.07 | 19.33 | 83.49 | 0.00 | 0.00 | 0.00 | 72.44 | 48.24 | 2.21 | 91.81 |
| S184 | 154 | 0.94 | 0.00 | 13.32 | 0.00 | 0.00 | 0.00 | 1.42 | 0.00 | 0.00 | 32.84 |
| S185 | 155 | 0.94 | 0.00 | 13.32 | 0.00 | 0.00 | 0.00 | 5.06 | 0.00 | 0.00 | 0.00 |
| S191 | 161 | 58.14 | 3.60 | 47.87 | 0.91 | 0.00 | 0.00 | 8.57 | 20.89 | 0.00 | 0.00 |
| S195 | 165 | 21.04 | 1.01 | 0.01 | 0.00 | 0.00 | 0.00 | 0.00 | 1.10 | 0.00 | 10.86 |
| Y197 | 167 | 1.30 | 0.00 | 0.00 | 0.00 | 0.00 | 0.00 | 0.00 | 0.02 | 0.00 | 0.00 |
| S198 | 168 | 0.70 | 0.02 | 2.74 | 0.48 | 0.00 | 0.00 | 0.86 | 0.00 | 1.28 | 90.34 |
| S199 | 169 | 32.32 | 0.04 | 0.00 | 0.17 | 0.00 | 0.00 | 2.09 | 0.33 | 0.00 | 14.96 |
| S202 | 172 | 22.25 | 1.84 | 0.04 | 0.04 | 0.03 | 0.00 | 6.64 | 6.78 | 8.07 | 12.69 |
| T205 | 175 | 3.49 | 0.08 | 0.02 | 0.00 | 0.00 | 0.00 | 0.77 | 1.54 | 0.00 | 75.11 |
| S208 | 178 | 0.25 | 0.26 | 0.97 | 0.00 | 0.00 | 0.00 | 0.00 | 0.02 | 1.28 | 69.97 |
| S210 | 180 | 7.70 | 13.16 | 12.40 | 10.59 | 99.13 | 99.00 | 91.32 | 91.91 | 92.49 | 91.61 |
| T212 | 182 | 0.01 | 0.00 | 0.23 | 0.00 | 0.00 | 28.52 | 48.19 | 77.12 | 83.24 | 68.18 |
| S214 | 184 | 0.07 | 0.01 | 0.08 | 8.69 | 6.10 | 0.00 | 12.05 | 28.24 | 14.52 | 96.53 |
| T217 | 187 | 0.76 | 4.23 | 0.57 | 1.77 | 0.00 | 0.12 | 0.00 | 0.33 | 0.00 | 50.90 |
| T220 | 190 | 0.00 | 0.00 | 0.02 | 0.00 | 0.00 | 0.00 | 0.00 | 0.98 | 0.00 | 11.01 |
| T231 | 201 | 95.97 | 94.11 | 0.00 | 0.00 | 0.00 | 0.00 | 97.32 | 91.28 | 54.76 | 94.13 |
| S235 | 205 | 68.36 | 30.80 | 81.68 | 9.63 | 0.08 | 0.00 | 70.57 | 2.59 | 0.00 | 58.36 |
| S237 | 207 | 83.56 | 59.23 | 0.00 | 0.00 | 0.00 | 0.00 | 81.13 | 0.00 | 11.85 | 0.00 |
| S241 | 211 | 0.00 | 0.00 | 0.00 | 0.19 | 0.06 | 0.00 | 0.17 | 0.00 | 0.00 | 0.00 |
| T245 | 215 | 0.01 | 0.01 | 0.00 | 0.35 | 0.01 | 0.00 | 0.28 | 0.02 | 0.00 | 0.02 |
| S258 | 228 | 0.00 | 1.28 | 0.00 | 0.00 | 0.00 | 0.00 | 10.73 | 1.47 | 0.00 | 0.27 |
| S262 | 232 | 0.06 | 0.27 | 0.05 | 0.01 | 0.00 | 0.00 | 0.45 | 0.16 | 0.00 | 1.42 |
| T263 | 233 | 0.69 | 0.00 | 0.00 | 0.00 | 0.00 | 0.00 | 2.90 | 0.00 | 0.00 | 0.21 |
| S285 | 255 | 0.00 | 0.00 | 0.00 | 0.00 | 0.00 | 0.00 | 0.00 | 0.01 | 0.00 | 0.00 |
| S289 | 259 | 0.00 | 0.01 | 0.01 | 0.00 | 0.00 | 0.00 | 0.13 | 0.05 | 0.00 | 0.06 |
| S305 | 275 | 1.59 | 1.34 | 0.73 | 15.08 | 31.19 | 0.99 | 21.15 | 7.84 | 0.83 | 7.68 |

| | | | | | | | | | | | |
|---|---|---|---|---|---|---|---|---|---|---|---|
| Y310 | 280 | 0.02 | 0.03 | 0.01 | 0.00 | 0.00 | 0.00 | 0.02 | 0.02 | 0.00 | 0.05 |
| T319 | 289 | | | | 0.00 | 0.00 | 0.00 | 0.00 | 0.00 | 0.00 | 76.40 |
| S320 | 290 | | | | 0.00 | 0.00 | 0.00 | 0.00 | 99.12 | 0.00 | 48.44 |
| S324 | 294 | 6.05 | 0.71 | 0.00 | 99.75 | 47.85 | 0.99 | 64.59 | 49.27 | 4.35 | 13.96 |
| S352 | 322 | 32.42 | 1.53 | 0.00 | 94.69 | 0.48 | 1.48 | 99.36 | 56.14 | 0.52 | 40.36 |
| S356 | 326 | 0.74 | 0.41 | 0.04 | 18.83 | 5.10 | 2.42 | 23.25 | 14.66 | 0.44 | 4.46 |
| T361 | 331 | 0.00 | 0.00 | 0.00 | 0.00 | 0.00 | 0.00 | 0.00 | 0.03 | 0.00 | 0.22 |
| T373 | 343 | 0.00 | 1.30 | 52.84 | 0.00 | 0.00 | 0.00 | 0.00 | 0.68 | 0.00 | 53.90 |
| T377 | 347 | 84.90 | 3.11 | 0.00 | 65.25 | 0.00 | 0.00 | 78.31 | 3.40 | 0.00 | 4.80 |
| T386 | 356 | 0.00 | 0.00 | 0.00 | 0.09 | 0.00 | 0.00 | 0.00 | 0.00 | 0.00 | 0.00 |
| Y394 | 364 | 16.07 | 0.19 | 0.00 | 0.04 | 0.00 | 0.00 | 16.96 | 4.55 | 0.33 | 3.78 |
| S396 | 366 | 30.31 | 3.04 | 1.43 | 0.05 | 0.00 | 0.00 | 44.26 | 21.04 | 0.64 | 2.36 |
| S400 | 370 | 61.44 | 11.69 | 3.02 | 0.03 | 0.00 | 0.00 | 48.54 | 55.20 | 0.31 | 1.73 |
| T403 | 373 | 32.92 | 7.13 | 10.69 | 0.00 | 0.00 | 0.00 | 22.49 | 18.48 | 6.12 | 8.65 |
| S404 | 374 | 32.69 | 1.36 | 0.28 | 0.00 | 0.00 | 0.00 | 56.38 | 36.88 | 6.12 | 8.70 |
| S409 | 379 | 0.25 | 0.00 | 0.00 | 78.13 | 0.00 | 0.00 | 44.42 | 0.00 | 0.00 | 0.56 |
| S412 | 382 | 0.27 | 0.00 | 0.00 | 0.00 | 0.00 | 0.00 | 0.00 | 0.00 | 0.00 | 0.00 |
| S413 | 383 | 0.00 | 0.00 | 0.00 | 0.07 | 0.00 | 0.00 | 0.00 | 0.00 | 0.00 | 0.00 |
| T414 | 384 | 0.00 | 0.00 | 0.00 | 0.08 | 0.00 | 0.00 | 0.00 | 0.00 | 0.00 | 0.00 |
| S416 | 386 | 0.10 | 0.00 | 0.00 | 78.13 | 0.00 | 0.00 | 8.04 | 0.00 | 0.00 | 2.23 |
| S422 | 392 | 0.04 | 0.00 | 0.00 | 0.44 | 0.00 | 0.00 | 0.00 | 0.00 | 0.00 | 0.00 |
| T427 | 397 | 0.00 | 0.00 | 0.00 | 0.67 | 0.00 | 0.00 | 0.00 | 0.00 | 0.00 | 0.00 |
| **Sites detected with over 10% occupancy** (2N4R nomenclature) | | **GSK**<br>T181, S191, S199, S210, T231, S235, S237, S352, T373, T377, S396, S400, T403, S404 | | | **PKA**<br>S210, S305, S324, S352, T377, S409, S416 | | | **PKA-GSK**<br>T149, T153, T181, S198, T205, S208, S210, T212, S214, T217, T319, S320, S324, T231, S235, S237, S352, S356, T373, T377, S396, S400, T403, S404, S409 | | | |
| **Special attributes** | | -Nuclear tau sites (T181, S404)<br>-Phosphorylation between PRD DNA-binding regions (T231, S235, S237)<br>-Trace (<10%) AT8 S202/T205 | | | -Trace (<10%) AT100 T212/S214 | | | -Nuclear tau sites (T181, T212, S404)<br>-AT8 - trace S202 and T205<br>-AT100 T212/S214<br>-Phosphorylation between PRD DNA-binding regions (T231) | | | |
